# MePMe-seq: Antibody-free simultaneous m^6^A and m^5^C mapping in mRNA by metabolic propargyl labeling and sequencing

**DOI:** 10.1101/2022.03.16.484494

**Authors:** Katja Hartstock, Anna Ovcharenko, Nadine A. Kueck, Petr Spacek, Nicolas V. Cornelissen, Sabine Hüwel, Christoph Dieterich, Andrea Rentmeister

## Abstract

Internal modifications of mRNA have emerged as widespread and versatile regulatory mechanism to control gene expression at the post-transcriptional level. Current insights rely on the ability to make a modified nucleoside amenable to sequencing. Most of the modifications are methylations involving the co-factor *S*-adenosyl-L-methionine (SAM), however, simultaneous detection of different methylation sites in the same sample has remained elusive. We present metabolic labeling with propargyl-selenohomocysteine (PSH) in combination with click chemistry to detect *N*^6^- methyladenosine (m^6^A) and 5-methylcytidine (m^5^C) sites in mRNA with single nucleotide precision in the same sequencing run (MePMe-seq). Intracellular formation of the corresponding SAM analogue leads to detectable levels of *N*^6^-propargyl-A (prop^6^A) and 5-propargyl-C (prop^5^C). MePMe-seq overcomes the problems of antibodies for enrichment and sequence-motifs for evaluation, limiting previous methodologies. The joint evaluation of m^6^A and m^5^C sites opens the door to study their interconnectivity and improve our understanding of mechanisms and functions of the RNA methylome.

## Main

Eukaryotic mRNA is canonically modified by addition of the 5’ cap and bears additional modifications at internal sites. The *N*^6^-methylation of adenosine is the most abundant and best studied internal modification of mRNA. It has been linked to cellular differentiation, cancer progression and development^1-6^. Most of the more than 12,000 sites are introduced by the METTL3-14 complex, whereas METTL16 is responsible for six additional validated sites in mRNA ^7-11^. Several reader proteins have been identified and mediate the effects of m^6^A in mRNA translation and degradation ^12-24^. m^5^C is the second most abundant internal modification in mammalian mRNA. Reported numbers range from a few hundred to 40,000 sites and different writer proteins (NSUN2, NSUN6 and TRDMT1) have been reported for mammalian cells ^25-28^. Several reader proteins have been identified, linking m^5^C to repair (via RAD52), export (via ALYREF), and proliferation (via YBX1 and ELAV) causing bladder cancer ^29-33^. In total, ten different internal modifications of eukaryotic mRNA have been described and mapped ^34^. In addition to m^6^A and m^5^C, these comprise the altered nucleobases inosine and pseudouridine, the acetylation ac^4^C, the oxidation 8-oxo-G and further methylations or derivatives thereof at either the nucleobase (m^1^A, m^7^G, hm^5^C) or the ribose (N_m_). The prevalence of methylation as mRNA modification mark is striking, suggesting that the responsible cofactor SAM plays a key role for their abundance and potential interconnectivity^35^.

Owing to the importance and abundance of m^6^A, multiple approaches have been developed to assign its positions on a transcriptome-wide level. Most methods rely on antibodies in combination with next generation sequencing (NGS). While early transcriptome-wide detection methods had limited resolution ^36-38^, crosslinking and bioinformatic analyses including search for the DRACH motif, improved the accuracy to single nucleotides ^39-41^. Concerns regarding bias of antibodies in recognizing the tiny nucleobase as epitope along with the inability to distinguish between m^6^A and m^6^A_m_ has prompted the development of antibody-free methods. DART-seq uses a YTH reader protein for m^6^A recognition and introduces adjacent C-to-U mutations by a fused deaminase ^42, 43^. MAZTER seq relies on the methylation-sensitive ribonuclease and bioinformatic alignment of cleaved versus uncleaved sequences at its target ACA sites^44, 45^. m^6^A-SEAL uses m^6^A-specific methyl oxidation by FTO for further derivatization and enrichment ^46^. m^6^A-REF-seq combines m^6^A demethylation by FTO and cleavage of ACA-sites with a m^6^A-sensitive RNA endonuclease ^47^. However, these methods rely on enzymes for modification, which bring about their own biases, such as preference for or even limitation to certain sequences. Furthermore, some of these approaches require transfection of cells.

Metabolic labeling with methionine (Met) analogues therefore presents an interesting alternative approach for m^6^A detection ^48, 49^. After feeding cells with PSH, modified adenosines in rRNA could be detected via click chemistry and enrichment ^49^. Label-Seq determined m^6^A-sites in mRNA by feeding allyl-selenohomcysteine followed by a highly specific cyclization reaction of the resulting *N*^6^- allyladenosine, causing mutations in RT identified by NGS. However, an antibody was required to enrich the allyl-modified mRNA and other modified nucleosides were not detected^48^.

For m^5^C, the second most abundant internal modification of mRNA, chemical conversion of C to U in bisulfite sequencing is most widely used for mapping ^50, 51^. This treatment risks damaging RNA and causing artifacts, necessitating careful und repeated controls to obtain reliable data^52^. Antibody-based methods with and without photo-crosslinking have been developed in analogy to m^6^A mapping methods and underlie the same limitations ^25, 53, 54^. The development of antibody-free methods includes progress towards nanopore sequencing and TAWO-seq, but the latter has not yet been implemented on a transcriptome-wide scale. ^55, 56^.

Taken together, both m^6^A and m^5^C as well as other methylations rely on the cofactor SAM, suggesting that the methyl-based modifications could be interconnected via SAM levels, and it would be important to study them in context^35^. However, current methodology has mainly focused on specific binding and detection of the modified nucleoside instead of the underlying and unifying process. The enrichment via antibodies or binding-proteins, or the specific modification by m^6^A-sensitive enzymes counteracts a more global look at possible links.

We therefore set out to develop metabolic labeling via the SAM pathway as methodology to detect more than one type of modified nucleoside by NGS. Such methodology should hinge on a SAM analogue that (1) can be efficiently made in genetically unaltered mammalian cells, (2) is accepted by promiscuous activity of several methyltransferases, and (3) provides a universal handle for efficient antibody-free enrichment of different nucleosides to (4) make modified nucleosides amenable to detection in NGS. The SAM analogue SeAdoYn presents an ideal metabolite, which is accepted by many methyltransferases and efficiently generated in genetically unaltered cells ^49, 57^. The propargyl group is above all bioorthogonal and specifically reacts with azides in a click reaction, providing a way to chemically enrich target RNA—completely independent of antibodies (Fig. 1a-e).

**Fig. 1:**
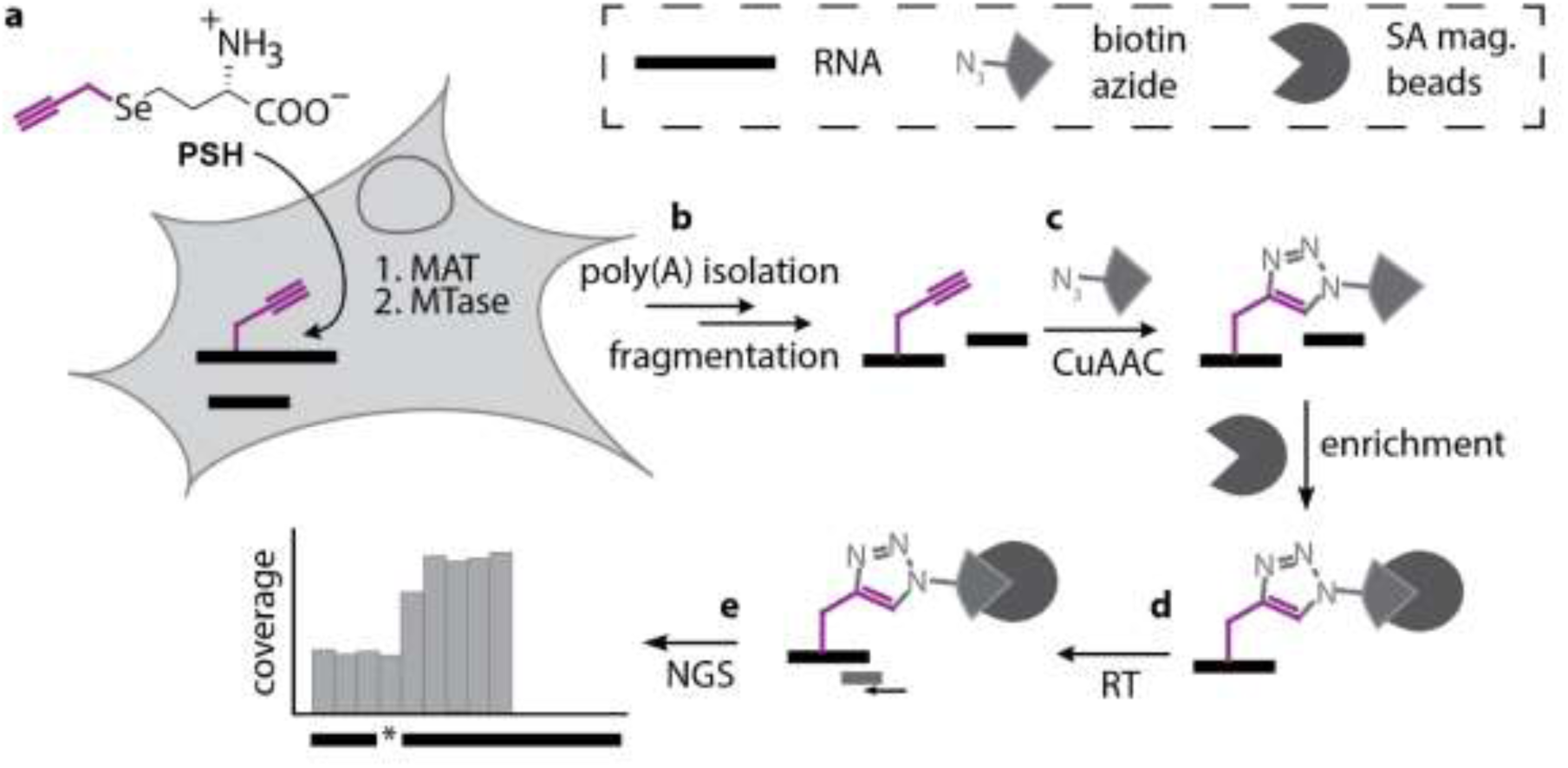
Scheme of MePMe-seq (<u>me</u>tabolic <u>p</u>ropargylation for <u>me</u>thylation <u>seq</u>uencing). **a**, Metabolic labeling of cells with PSH leads to methionine adenosyl transferase (MAT)-catalyzed formation of SAM-analogue and propargylation of methyltransferase (MTase) target sites. **b**, After cell lysis, poly(A)^+^ RNA is isolated and fragmented. **c**, Propargylated fragments react with biotin azide in a copper-catalyzed azide-alkyne cycloaddition (CuAAC) and are bound to streptavidin-coated magnetic beads. **d**, On-bead reverse transcription (RT) terminates at modified sites. **e**, Libraries for NGS are prepared. Modified sites are detected as steep coverage drops.

## Results

### Metabolic labeling and quantification of propargylated nucleosides

To optimize intracellular propargylation of RNA by metabolic labeling with PSH, we treated HeLa cells with different concentrations of PSH or Met (Supplementary Fig. 3a). For quantification, we performed mass spectrometric (MS) analysis by LC-QqQ-MS (liquid chromatography in tandem with triple quadrupole mass spectrometer) of RNA after digestion to nucleosides and referenced to the respective synthetic standards (Fig. 2a-c, Supplementary Fig. 1-2) ^58^. This analysis showed that the level of propargylated A (A_prop_/A) was increased at higher (up to 2.5 mM) PSH concentrations (Supplementary Fig. 3b). Under these conditions, 0.1 % of all As were A_prop_ (Supplementary Fig. 3b), and 2.2 % of A_m_ were substituted by A_prop_ (Supplementary Fig. 3c). In total RNA, the ratio of A_m_/A was ∼4 % and remained largely unaffected by metabolic PSH labeling, suggesting that the general cellular methylation itself is not perturbed (Supplementary Fig. 3d). 2.5 mM PSH were chosen for subsequent metabolic labeling and controls were treated identically but using Met instead of PSH. Under these conditions, cell viability was 81 % (Supplementary Fig. 3f).

**Fig. 2:**
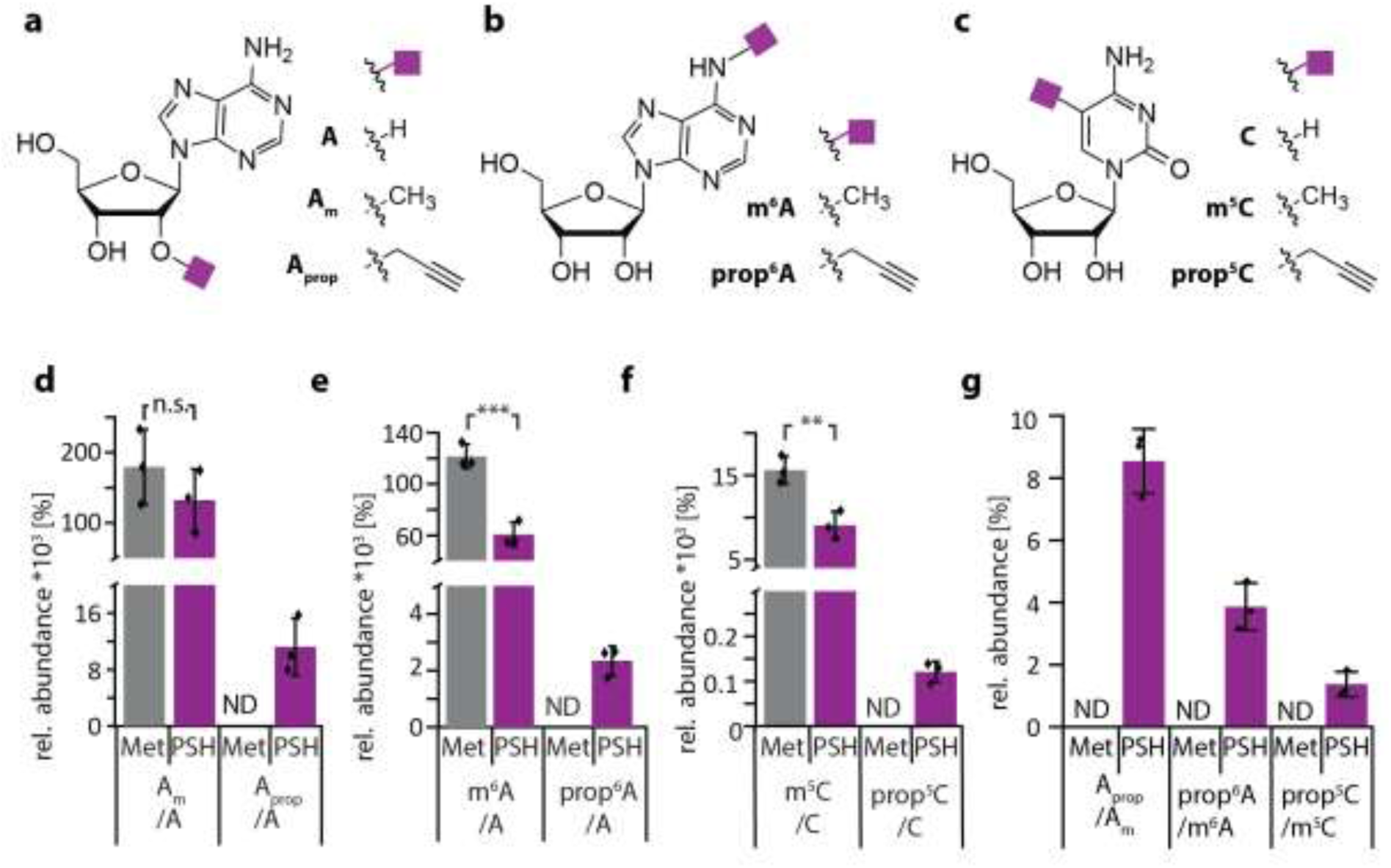
Metabolic labeling with PSH. **a-c**, Modified and unmodified nucleosides detected via LC-QqQ-MS. Structures of (**a**) adenosine (A), 2’-O-methyl adenosine (A_m_), 2′-O-propargyl adenosine (A_prop_), (**b**) *N*^6^-methyl adenosine (m^6^A), *N*^6^-propargyl adenosine, (**c**) cytidine (C), 5-methylcytidine (m^5^C) and 5-propargylcytidine (prop^5^C). **d-g**, Quantification of modified nucleotides in mRNA from HeLa cells treated with 2.5 mM PSH or methionine (Met) as control. (**d**) A_m_ and A_prop_ relative to A, (**e**) m^6^A and prop^6^A relative to A, (**f**) m^5^C and prop^5^C relative to C, and (**g**) each of the identified propargylated nucleosides relative to their methylated analogue. Quantification from dynamic MRM run on LC-QqQ-MS using external synthetic standards. Not detected (ND) means no signal with correct quantifier detected. Mean values and SD from n=3 biological replicates are shown. Statistical significance determined via unpaired two-sample two-tailed t-test (n.s. P>0.05; * P≤0.05; ** P≤0.01; *** P≤0.001).

Next, we analyzed propargylation and methylation levels of adenosine in mRNA. After two times of poly(A)-enrichment, we measured 0.011 % of A_prop_/A in RNA from labeled cells, whereas A_prop_ was not detectable in control cells (Fig. 2d). The A_m_/A ratio in mRNA was ∼20-fold lower than in total RNA, confirming successful poly(A)-enrichment (Fig. 2d, Supplementary Fig. 3d). The A_m_/A ratio was not significantly different in labeled and unlabeled cells (0.13–0.18 %, Fig. 2d) but the m^6^A/A levels were reduced from 0.12 % to 0.06 % by PSH labeling (Fig. 2e), suggesting a stronger effect of metabolic labeling on mRNAs, which have a shorter half-life than rRNAs. The observed ∼10-fold decrease in the A_prop_/A ratio in mRNA (0.011%, Fig. 2d) compared to total RNA (0.10%, Supplementary Fig. 3b) is in line with the abundance of the natural modifications in these samples, suggesting that metabolic labeling does not alter the ratio of modifications. In contrast, m^6^A is only slightly decreased by mRNA isolation. In mRNA from labeled cells, prop^6^A was unambiguously detected, whereas it was not detectable in control cells (Fig. 2e). Prop^6^A replaced 3.5 % of m^6^A (Fig. 2g) and was present in 2.3*10^−3^ % of all As (Fig. 2e). Taken together, metabolic labeling of HeLa cells with PSH leads to detectable levels of prop^6^A in mRNA suggesting that the propargyl group will be available for enrichment and detection of m^6^A sites.

### Transcriptome-wide analysis of m^6^A from metabolic labeling with PSH

Next, we investigated the transcriptome-wide identification of m^6^A and potentially other modified nucleosides at single nucleotide resolution. To this end, we isolated mRNA from cells after metabolic labeling with PSH, and reacted it with biotin-azide to enrich propargylated mRNA on streptavidin-coated magnetic beads (Fig. 1a). The conversion was complete (Supplementary Fig. 4) and after reverse transcription using Superscript (SSIV) under conditions optimized to yield 80 % of termination (Supplementary Fig. 5), we prepared libraries for NGS using an adapted iCLIP2 protocol ^59^. The reads were preprocessed (FASTQ processing, barcode filtering and quality control), mapped to the human genome (hg38) and duplicate reads removed based on the introduced unique molecule identifier (UMIs), resulting in 14.8 (rep1) and 15.5 (rep2) million reads for the PSH treated samples (see SupplementaryTable 1).

Visual inspection of the coverage profiles for known and validated m^6^A sites showed remarkably sharp edges one nucleotide downstream of m^6^A sites. This is exemplified by the six m^6^A positions in the hairpins of MAT2A, the m^6^A2515 and m^6^A2577 in MALAT1 as well as m^6^A1216 in β-actin (Suppl. Fig. S6), which are known targets of METTL16 and METTL3-14, respectively ^7, 60, 61^. These data indicate (1) that metabolic labeling results in METTL3-14 as well as METTL16-mediated propargylation at the *N*^6^- position of adenosine in poly(A)^+^ RNA and (2) that reverse transcription in isolated mRNA terminates one nucleotide downstream of clicked sites, allowing assignment of m^6^A sites in mRNA with single nucleotide precision in NGS data.

For systematic analysis of MePMe-seq data on the transcriptome-wide level, we used JACUSA2 ^62^. This improved version of JACUSA, a software for site-specific identification of RNA editing events from replicate sequencing data, is able to identify read termination events by calculating the arrest rate, i.e. the fraction of reads stopping at the position, from NGS data ^62^. The difference between arrest rates from sample and control (Δ_(RT arrest)_) was used to filter the terminations identified by the algorithm and remove false positives. Setting a sample read coverage threshold (> 35 for high stringency (HS) filter, > 20 for low stringency (LS) filter) and arrest score threshold (*Δ*_*RT arrest*_ >20 for HS filter; *Δ*_*RT arrest*_>15 for LS filter), resulted in calling a total number of 8,802 (rep1) and 7,124 (rep2) termination sites for all four nucleotides, if high stringency settings were used (Supplementary data 1). Filtering with low stringency called 26,673 (rep1) and 27,869 sites (rep2) termination sites, respectively (Supplementary Fig. 7-8). JACUSA2 is available at https://github.com/dieterich-lab/JACUSA2.

Initial inspection of these terminations revealed clustering at transcription start sites (TSS) (Supplementary Fig. 9). Accordingly, NGS coverage profiles showed strong enrichment of 5’ end fragments (i.e. ∼150 nt regions) for many transcripts and steep drop-offs within this region, which were called as arrests by JACUSA2 (Supplementary Fig. 10). Based on early literature on metabolic labeling with radioactive Met ^63-65^, which led to the identification of multiple methylation sites at the 5’ cap, it is reasonable to assume that metabolic PSH labeling will also target the canonical cap methylation sites, resulting in the observed clustering. To rule out effects from metabolic cap labeling, we excluded regions <5 nt upstream from the TSS from analysis of internal modification sites. Using this computational pipeline and high stringency filtering for data analysis, MePMe-seq identified 5,506 (rep1) and 3,714 (rep2) modified internal sites for all four nucleosides in mRNA from PSH labeled cells (Supplementary Fig. 7). The modified nucleotides were predominantly adenosine (A, 70 %), followed by cytidine (C, 16 %), uridine (U, 9 %) and guanosine (G, 5 %) (Fig. 3a).

**Fig. 3:**
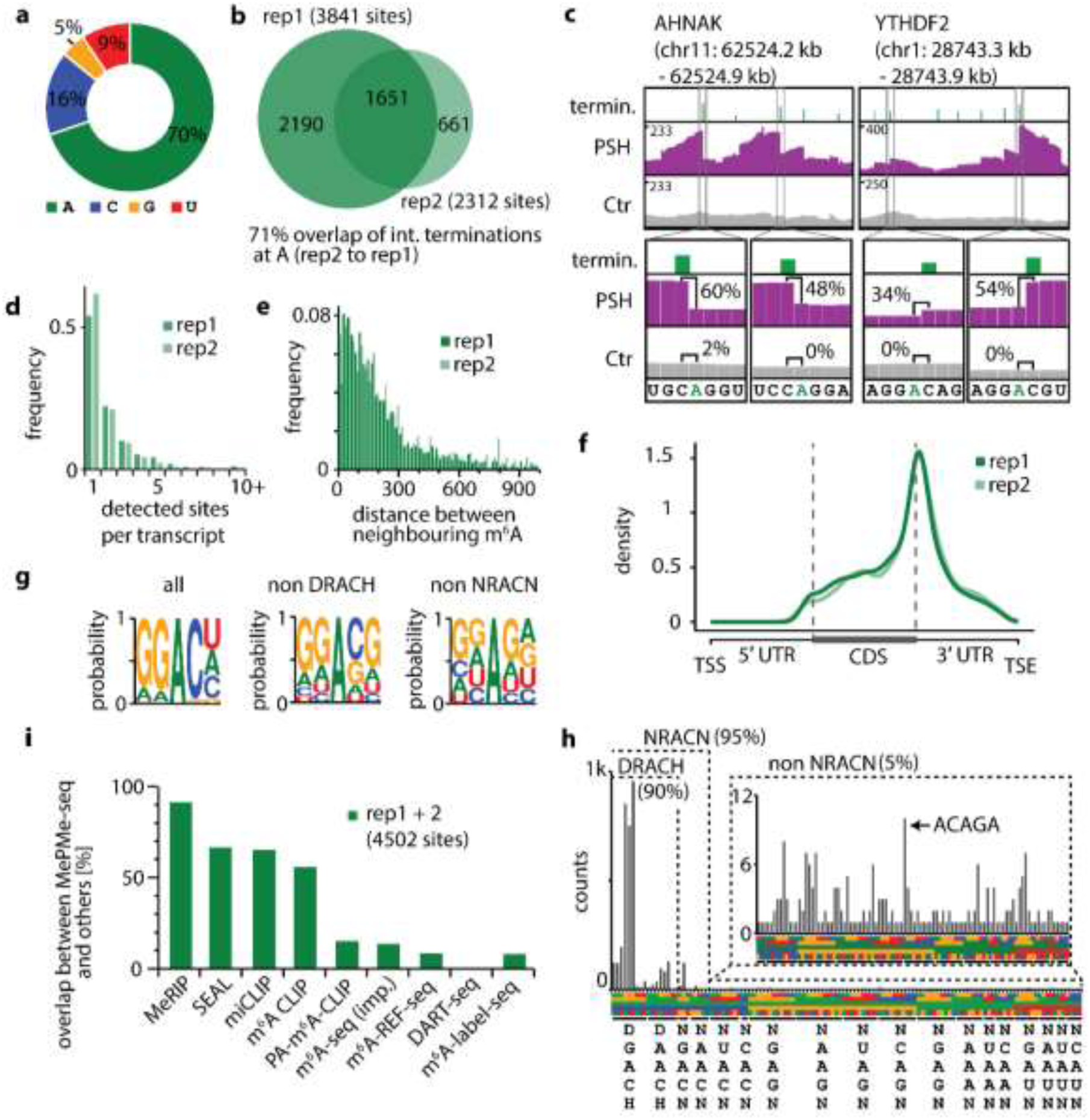
Detection of m^6^A sites using MePMe-seq. **a**, Distribution of all modified nucleotides identified in MePMe-seq. **b**, Overlap of m^6^A sites identified in n=2 independent experiments (% calculated from rep2 to rep1). **c**, Integrative genomics viewer (IGV) browser coverage tracks of MePMe-seq data for the indicated AHNAK and YTHDF2 mRNAs from cells labeled with PSH (purple) or methionine as control (gray). Green bars indicate terminations identified by JACUSA2 applying high stringency filtering. Numbers (%) for calculated arrest rate at indicated positions are shown. **d**, Frequency of m^6^A sites per transcript. **e**, Frequency of distance between neighboring m^6^A positions located on the same transcript. Cutoff at 1,000 nt (for cutoff at 5,000 nt see Supplementary Fig. 13). **f**, Metatranscript analysis showing a density plot of the distribution of prop^6^A sites detected by MePMe-seq. **g**, Consensus motif for sequences surrounding identified m^6^A (HS filtering) for all 5-mers (all), if DRACH sequences are excluded (non DRACH) or if NRACN sequences are excluded (non NRACN). Representative example of n=2 biologically independent samples is shown (2^nd^ replicate: Supplementary Fig. 14). **h**, Sequence motifs surrounding identified m^6^A sites (HS filtering), sorted by consensus motif DRACH, NRACN or non-NRACN, respectively. Arrow indicates ACAGA-motif, which is part of METTL16 motif. **i**, Overlap of all 4,502 m^6^A sites identified in MePMe-seq with sites identified by MeRIP, SEAL, miCLIP, m^6^A CLIP, PA-m^6^A-CLIP, m^6^A-seq improved (imp.), m^6^A-REF-seq, DART-seq and m^6^A-label-seq ^46, 48, 66, 67^.

We first focused on m^6^A as the most abundant modification in mRNA. In two biological replicates, MePMe-seq identified 3,841 (rep1) and 2,312 (rep2) internal As as MTase target sites in mRNA from HeLa cells using high stringency filtering (Fig. 3b). Of the modified As, 1,651 (71 %) were identified in both replicates, indicating very good reproducibility (Fig. 3b). With low stringency filtering, the number of called m^6^A sites from the same replicates was much higher, namely 13,865 and 10,693 (Supplementary Data 2), and the overlap was still good (45 %, Supplementary Fig. 11). Nevertheless, for all subsequent analyses, we relied on hits identified with high stringency filtering. This is most likely an underestimation of sites (Supplementary Fig. 11).

Inspection of the hits in the IGV showed drop-offs at the m^6^A sites identified by JACUSA2 in mRNA from labeled cells that are not observed in mRNA from control cells. This is illustrated for the AHNAK and the YTHDF2 mRNAs (Fig. 3c, Supplementary Fig. 12s), which are known for their high m^6^A content ^48, 66,67^. The drop-offs are remarkably sharp and correspond to termination one nucleotide downstream of the modified A, in line with our thorough *in vitro* evaluation (Supplementary Fig. 5). Arrest rates ranged from 34–60 %. These results demonstrate that MePMe-seq in combination with JACUSA2 analysis enables reliable calling of m^6^A sites with single-nucleotide precision.

Next, we looked at the abundance of m^6^A sites in individual transcripts. MePMe-seq identified methylated A in 1,834 different transcripts (Supplementary Data 2) (1,311 in rep2). In 54 % of these transcripts, a single methylated A was found (for rep2: 61 %) (Fig. 3d). In all other transcripts (i.e. 46 % in rep1 and 39 % in rep2, respectively), more than one methylated A was present, including some with >10 m^6^A sites (Fig. 3d, Supplementary Data 2). Some of the highest m^6^A densities were found on AHNAK, PLEC and YTHDF2 mRNAs (Fig. 3c, Supplementary Figs. 8, 12), in line with previous reports ^48^. We were particularly interested in these clustered m^6^A sites that currently pose a challenge to most of the m^6^A mapping methods. MePMe-seq identified a total of 80, 25, and 12 m^6^A sites for AHNAK, PLEC, and YTHDF2, respectively (56, 18, 10, in rep2). We calculated the distance between neighboring m^6^As on the same transcript and found that they tend to cluster in short distances (Fig. 3e, Supplementary Fig. 13, emphasizing the importance of precise assignment. In summary, MePMe-seq showed remarkable precision in assigning the position of m^6^A sites and identified m^6^A sites in very close proximity (<10 nt) to each other.

We looked at the distribution of m^6^A sites by performing a metagene analysis of all modified As detected by MePMe-seq. The density plot shows enrichment at the 3’ end of the coding sequence (CDS) and around the stop codons (Fig. 3f). This result is in line with the m^6^A distribution reported by various methods, confirming that metabolic PSH labeling in combination with MePMe-seq identifies natural m^6^A sites ^36, 37, 39^-^42, 48^. The metabolic propargylation does not seem to introduce bias, except for the heavily and canonically methylated 5’ cap region which had to be excluded from analysis.

Comparing 5-mer sequences around the identified methylated internal adenosines revealed DRACH as the prevailing motif (Fig. 3g, Supplementary Fig. 14) with an abundance of 90 % (most abundant: GG**A**CU 25 %, GG**A**CA 22 %, GG**A**CC 20 %, AG**A**CU 5 %, others < 5 % abundance, Fig. 3h), which has been reported previously as the main consensus motif for *N*^6^-methylation of A via METTL3-14 ^36, 37, 39, 41, 42, 48^. Interestingly, 10 % of the m^6^A sites identified by MePMe-seq are located in non-DRACH motifs (Fig. 3h). These are composed of NRACN sequences (5 % in total), which are closely related to the DRACH motif and non-NR**A**CN motifs (5 % in total) (Fig. 3h). The non-NRACN motifs do not share a consensus motif, but G is preferred over other nucleotides directly downstream of A (Fig. 3g). Within the non-NR**A**CN hits, the sequence AC**A**GA is most abundant (Fig. 3h). This sequence is part of the motif targeted by METTL16 ^68^. Of note, MePMe-seq identified all currently known methylation sites of METTL16, i.e. six sites in the 3’-UTR of MAT2A-mRNA, as well as the U6 snRNA (Supplementary Fig. 15). These non-DRACH sites escape antibody-based approaches and approaches relying on bioinformatics searches for the DRACH motif, like m^6^A-CLIP^40^. MePMe-seq is thus able to accurately detect m^6^A in non-DRACH contexts and provide data about the interconnectivity of different methylations in an unbiased manner ^66, 67^.

### Overlap with datasets from other m^6^A-mapping methods

m^6^A sites have been mapped previously using antibody-dependent and antibody-independent methods ^36-42, 44^-^48, 69, 70^. To compare MePMe-seq results with m^6^A sites found in previous studies, data originating from various methodologies were assembled from the databases REPIC ^66^, ATLAS ^67^ and publications ^46, 48^. However, in a pairwise comparison of published datasets from individual miCLIP experiments, the detected m^6^A sites differed significantly, even for the same cell line (Supplementary Table 2). We therefore combined the hits reported in different experiments to obtain an unbiased and more comprehensive reference dataset ^44^.

We found that 90 % of the m^6^A sites identified by MePMe-seq matched the reported MeRIP hits (Fig. 3i). 55-67 % of the MePMe-seq sites overlapped with m^6^A sites identified using SEAL, miCLIP or m^6^A CLIP. A lower fraction (8–15 %) of the MePMe-seq sites were found in other antibody-free single nucleotide resolution techniques, i.e. PA-m^6^A-seq, m^6^A-REF-seq, m^6^A-label-seq (Fig. 3i). Only 18 sites (0.4 %) of m^6^A sites identified by MePMe-seq were also reported in DART-seq. This fraction increases when the exact sites are extended: 5 % of m^6^A sites identified in MePMe-seq are in close proximity (±50 nt) to sites identified in DART-seq and the overlap between the techniques increases up to ∼11 % if an uncertainty range of ±150 nt is allowed (Supplementary Fig. 17, Supplementary data 2). Of note, the overlap of m^6^A sites detected by MePMe-seq is higher than the range obtained by comparison of other single nucleotide resolution methods with CLIP (10-45 %) and better than comparison between each other (0.3-7.7 %) (Supplementary Table 3, Supplementary data 3), suggesting that m^6^A sites reported by MePMe-seq are highly reliable.

### Independent validation of MePMe-seq m^6^A sites

To independently validate m^6^A sites identified by MePMe-seq, we performed SELECT, an elongation- and ligation-based qPCR amplification method with single-nucleotide resolution ^71^. We evaluated eight putative m^6^A sites in poly(A)^+^ RNA, five of them in a DRACH motif and three in a non-DRACH motif (Fig. 4a). Comparing the normalized Δ*C*_q_ values of qPCRs of samples with and without FTO treatment, a Δ*C*_q_>1 for + FTO indicated the presence of m^6^A. We found that all five chosen DRACH sites indeed contained m^6^A. This includes an m^6^A site in the mRNA coding for the serine/arginine repetitive matrix protein 2 (SRRM2) that was not reported before. Of the three tested non-DRACH sites, WDR6 and CTNNB1 mRNAs were confirmed to contain m^6^A. These sites have been reported before via MeRIP and SEAL, but not in any of the single-base resolution techniques. The putative non-DRACH m^6^A site in FLNB, however, could not be validated by SELECT. As FTO has sequential and structural preferences ^72^ it is conceivable that this m^6^A site is not preferably demethylated and therefore not detectable via SELECT. To test this hypothesis, we tried to validate a known non-DRACH m^6^A site located in the 3’ UTR of MAT2A forming a hairpin structure. Indeed, SELECT failed to detect this well-known non-DRACH m^6^A site, most likely as a result of its hairpin structure and lack of FTO-mediated demethylation (Supplementary Fig. 17).

**Fig. 4:**
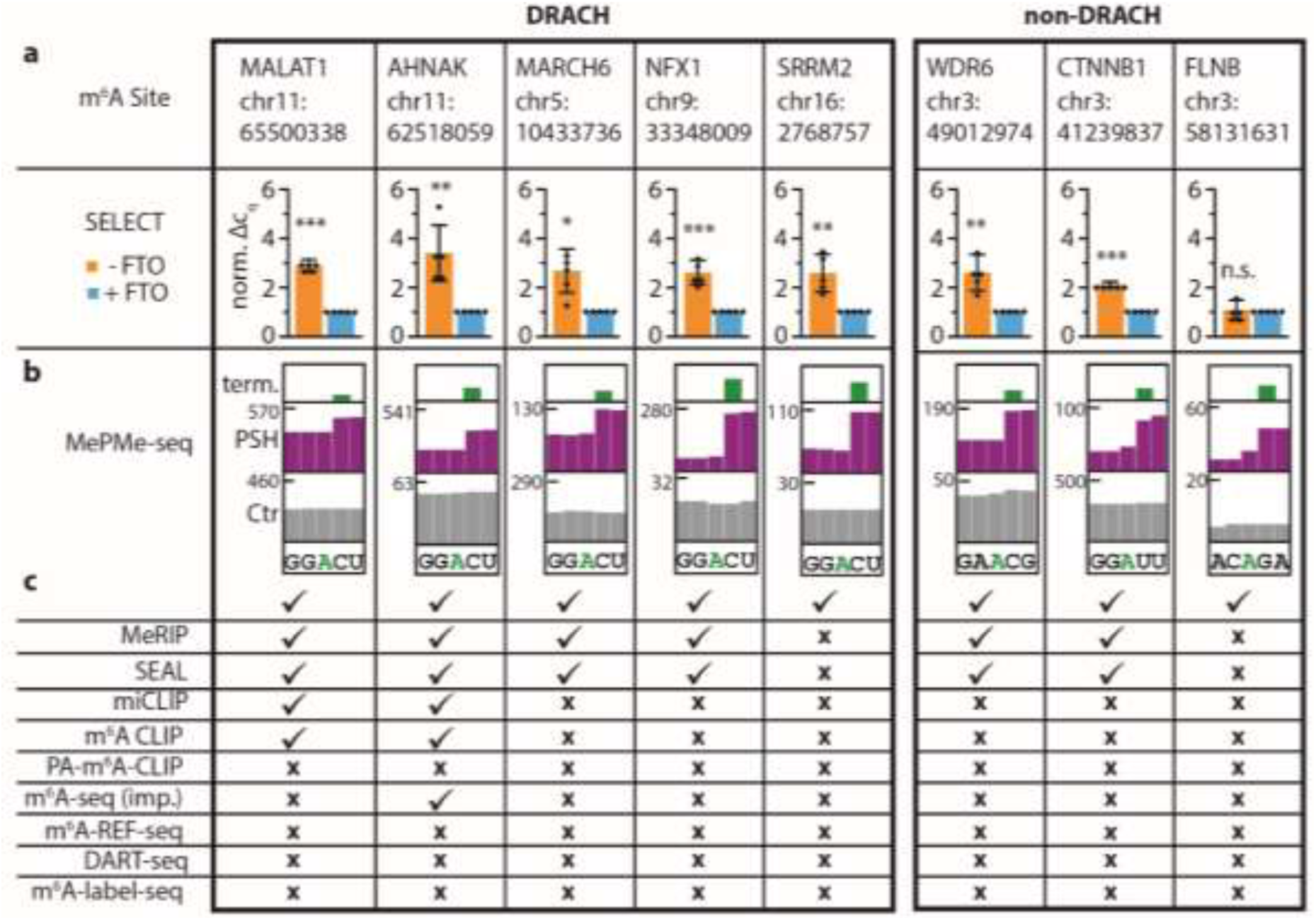
Validation of m^6^A sites identified in MePMe-seq via SELECT in HeLa poly(A)^+^ RNA. **a**, The normalized Δ*C*_q_ values of SELECT qPCR measurements are shown for five sites located in a DRACH motif and three sites located in a non-DRACH motif. **b**, IGV browser coverage tracks of MePMe-seq data for the same sites from cells grown with PSH (purple) or methionine (gray) as control. Green bars indicate terminations identified by JACUSA2. **c**, Comparison with MeRIP, SEAL, miCLIP, m^6^A CLIP, PA-m^6^A-CLIP, improved (imp.) m^6^A-seq, m^6^A-REF-seq, DART-seq or m^6^A-label-seq sequencing datasets^46, 48, 66, 67^. Checkmark for sites present, x for sites not present in dataset (data obtained from literature). Mean values and SD from n=5 biological replicates are shown. Statistical significance determined via one-sample one-tailed t-test (n.s. P>0.05; * P≤0.05; ** P≤0.01; *** P≤0.001).

In summary, all of the five tested m^6^A sites in a DRACH motif – including a previously unreported site – were confirmed by SELECT. In addition, two of the three putative m^6^A sites detected by MePMe-seq in non-DRACH sites were confirmed by SELECT, indicating that MePMe-seq is the first method with single nucleotide resolution able to detect m^6^A-sites in non-DRACH motifs. Since SELECT relies on FTO, bias originating from the enzyme’s substrate preference has to be considered. Therefore, it is conceivable that non-DRACH sites reported by MePMe-seq are true sites, even if confirmation by SELECT is not possible.

### METTL16-dependent labeling

MePMe-seq relies on intracellular formation of the SAM analogue SeAdoYn and therefore detects m^6^A sites originating from different MTases. While most m^6^A sites are METTL3-14 dependent, METTL16 is an emerging player in the RNA modification landscape of the human cell ^73^. METTL16 has been shown to bind a number of RNAs, including mRNAs and lncRNAs ^8, 73^, however methylation was only confirmed for six sites in MAT2A mRNA and U6 snRNA ^7^.

To pinpoint METTL16-dependent m^6^A sites, we isolated poly(A)^+^ RNA from untreated HeLa cells and propargylated it *in vitro* using recombinantly produced METTL16 and SeAdoYn (Fig. 5a, Supplementary Fig. 18). The *in vitro* propargylated mRNA was then processed as described above to enrich biotinylated RNA and determine the modification sites via termination in NGS. Visual inspection of the few known METTL16 target sites revealed steep coverage drops precisely one nucleotide upstream of the targeted adenosine in all cases, i.e. the hairpins in the 3’ UTR of the MAT2A-mRNA and the U6 snRNA (Fig. 5b, Supplementary Fig. 15, Supplementary data 4). These drops were exclusively found in the modified sample but not in a control sample and matched sites found by metabolic labeling, confirming that these are METTL16-dependent target sites that are also installed in intact cells.

**Fig. 5:**
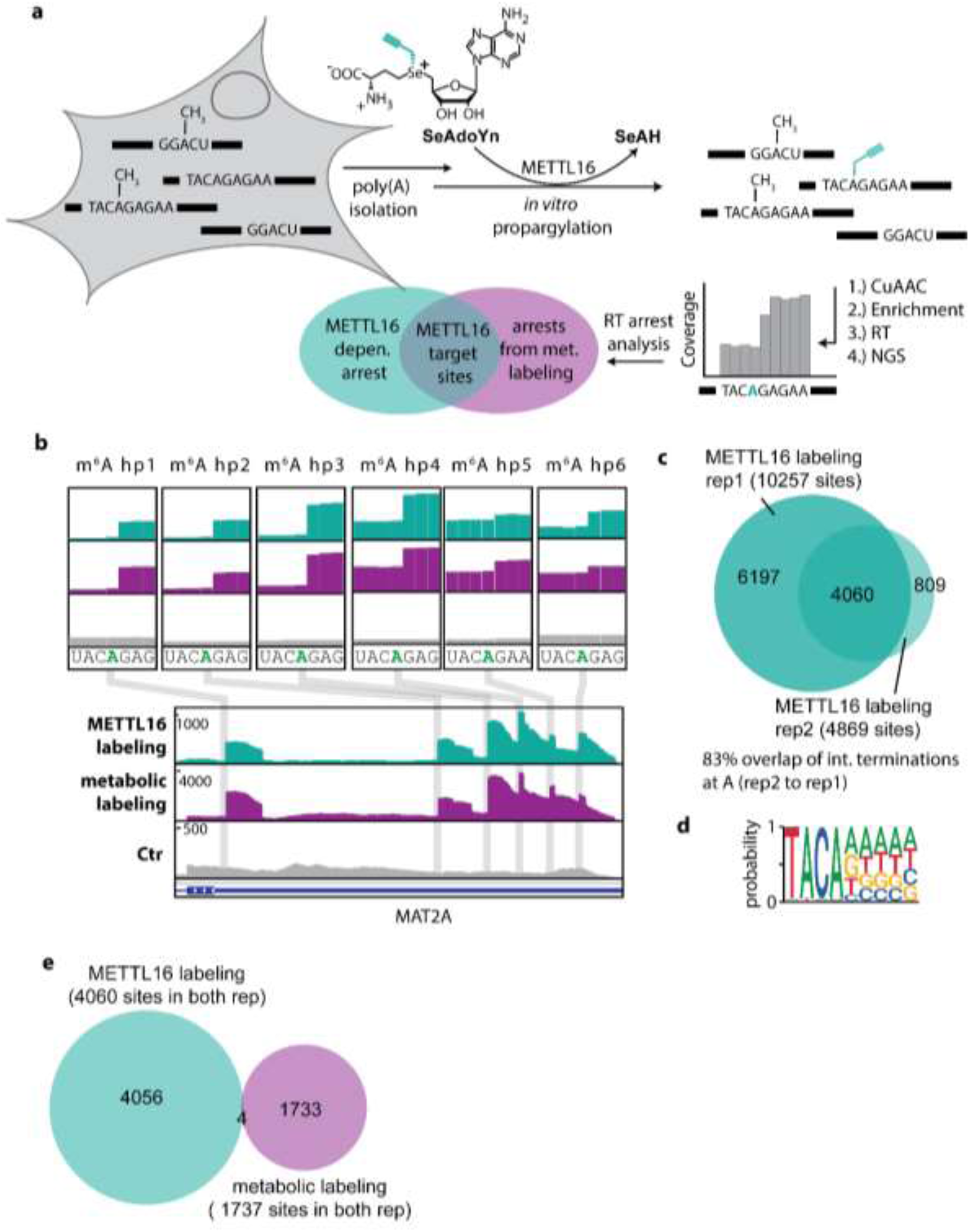
METTL16-dependent propargylation. **a**, Scheme illustrating METTL16-dependent labeling in combination with MePMe-seq to identify METTL16 target sites. Isolated mRNA is propargylated *in vitro* using METTL16 and analyzed by NGS. To eliminate false positive hits from *in vitro* off-target effects of METTL16, only hits observed also in MePMe-seq are identified as METTL16 targets. **b**, IGV browser coverage tracks for MAT2A-mRNA mapped by METT16-labeling *in vitro* (cyan) or MePMe-seq (purple), or control (gray). **c**, Overlap of identified m^6^A sites in n=2 independent METTL16-labeling experiments. **d**, Consensus motif for sequences surrounding identified As after METTL16-dependent labeling *in vitro*. **e**, Overlap of identified m^6^A sites in METTL16-dependent *in vitro* and metabolic labeling for sites present in n=2 independent experiments with HS filtering.

Sequencing and evaluation yielded 10,257 and 4,869 putative METTL16 target sites in two independent replicates using high stringency filtering (Fig. 5c, Supplementary data 5). Of these sites, 4,060 were found in both replicates (i.e. 83 % overlap), indicating very good reproducibility. Low stringency filtering yielded 40,307 and 38,552 sites respectively, but only 29 % (11,134 sites) overlap (Supplementary Fig. 19) and was not considered in further analyses. Within these hits, we inspected previously reported interaction sites of METTL16, such as STUB1, RBM3, MYC, NT5DC2, GNPTG, GMIP and MALAT1 ^8, 74^, for which it is unclear, whether they are also methylated by METTL16. Interestingly, we detected several of these sites in both replicates for MYC, RBM3, NT5DC2 and MALAT1 (Supplementary Fig. 20, Supplementary Table 4), providing evidence that METTL16 is indeed able to modify them *in vitro*. We observed multiple METTL16-dependent sites in the cancer-associated MALAT1 lncRNA, however, A8290 was not methylated *in vitro* (Supplementary Fig. 21). This is of particular interest, as A8290 was shown to interact with METTL16 but could not be validated as methylation target ^73, 75^. Analysis of the sequence motif adjacent to the m^6^A sites resulting from *in vitro* METTL16 labeling, identified a TACAD (Fig. 5d) motif, containing the reported METTL16 consensus motif TACA motif ^68^.

The large number of METTL16 sites identified by *in vitro* labeling is in stark contrast to the small number of confirmed sites. *In vitro* modification of RNA has also been used in other methods ^76^, however, we wondered whether the non-natural conditions could lead to off-target modification by METTL16 *in vitro*. To unambiguously identify METTL16 sites, we therefore matched the data from *in vitro* METTL16 labeling with the data from metabolic labeling. Hits identified in both approaches should be relevant METTL16 sites in cells. For this comparison, we used sites appearing in both replicates of *in vitro* METTL16 labeling (4,060 hits) and MePMe-seq (1,737 hits) and identified only four overlapping sites as hits (Fig. 5e). This indeed suggests that a large fraction of hits from *in vitro* METTL16 labeling result from off-target effects. Analysis of the four hits showed that these are the previously reported METTL16 target sites in the 3’-UTR of MAT2A ^68^. Two of the reported METTL16 hits in mRNA escaped this assignment using the stringent filtering conditions but were detected in either rep1 or rep2 of MePMe-seq (using high stringency filtering; Supplementary data 3).

When we inspected A8290 from MALAT1, which was not modified by METTL16 *in vitro*, we found that this site was not called by metabolic labeling either (Supplementary Fig.21, Supplementary Table 4). However, several m^6^A sites in close proximity to the putative METTL16 target site in MALAT1 could be clearly assigned owing to the high precision by MePMe-seq. Based on the combined analysis of *in vitro* and metabolic labeling we can now exclude A8290 in MALAT1 as a target of METTL16.

In summary, the *in vitro* modification data of METTL16 show that real and off-targets are detected within the consensus motif TACAD when applied *in vitro* at high concentrations. As several of the METTL16-dependent *in vitro* sites coincide with the interaction sites identified by CRAC, it could mean that METTL16 binds and – with SeAdoYn – can modify them. It cannot be excluded that additional proteins/RNAs as cofactors facilitate METTL16-dependent methylation in cells. We could show that the combination of *in vitro* and metabolic labeling provides a reliable protocol to assign the m^6^A sites to a certain methyltransferase and determine its target sites with single nucleotide precision.

### MePMe-seq identifies m^5^C sites in mRNA

Metabolic labeling with radioactive Met indicated that m^5^C is the second most abundant methylated nucleoside in poly(A)^+^-RNA, apart from the 5’ cap ^63, 65^. Following this trend, MePMe-seq identified 16 % of terminations one nucleotide upstream of cytidines (Fig. 3a). A total number of 1,305 sites matched a modified C. Specifically, 875 sites were found in rep1, 726 sites in rep2, and 325 (44 %) in both replicates, using high stringency filtering (Fig. 6a). These data suggest that cytosines in mRNA become propargylated during metabolic labeling and can be detected in MePMe-seq. We therefore compared previously reported m^5^C and C_m_ sites with our termination data ^67^. MePMe-seq identified sites in Furin- and Peroxidasin (PXDN) mRNA that matched m^5^C positions identified by bisulfite sequencing ^67^, suggesting that metabolic PSH labeling can be used to detect also m^5^C sites. None of the reported C_m_ positions matched terminations in our datasets, indicating that MePMe-seq detects 5-methylated rather than 2’-O-methylated Cs.

Visual inspection of the sequencing data for FURIN- and PXDN-mRNAs shows sharp drop-offs not only for m^6^A sites but also for m^5^C sites (Fig. 6b). Furthermore, these mRNAs appear enriched compared to the control samples that had not been treated with PSH, supporting installation of propargyl groups as handles for click chemistry and biotin-based isolation of modified sites (Fig. 6b). Terminations downstream of cytosine occurred in most cases only once per transcript (Fig. 6c) and were located mainly at the end of the 5’ UTR (Fig. 6d). The frequency and distribution for m^5^C sites are thus markedly different from m^6^A (Fig. 2f), in line with previous reports about m^5^C ^30, 77-79^. Analysis of the sequence context did not reveal a clear consensus motif (Fig. 6e, Supplementary Fig. 22). This result is in accordance with literature for type I m^5^C sites ^50, 80, 81^. The known motif for type II m^5^C sites (**C**UCCA) can be found in the results but is not prevalent, in line with their low abundance (Supplementary Fig. 23) ^81^.

**Fig. 6:**
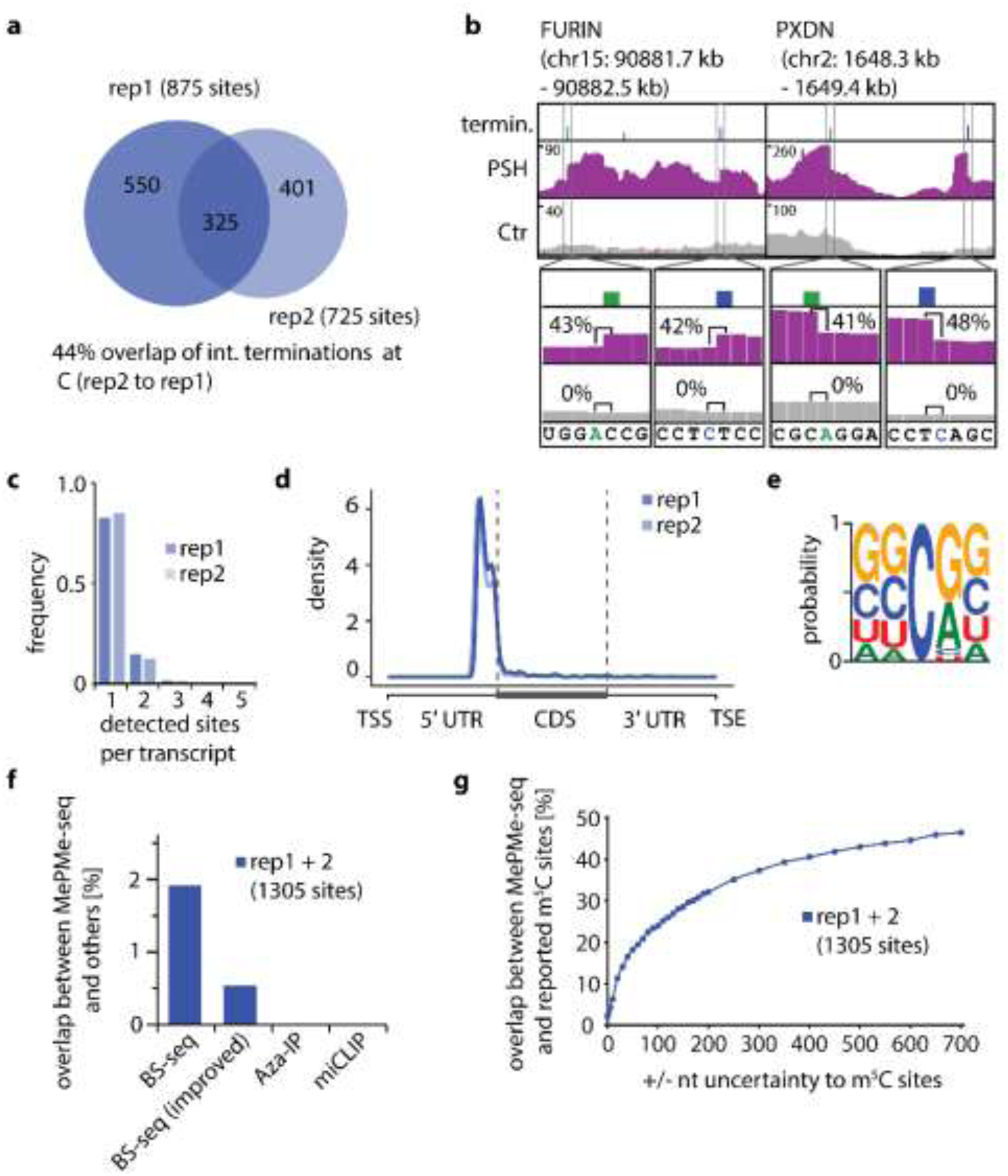
Detection of m^5^C sites using MePMe-seq. **a**, Overlap of identified m^5^C sites between n=2 MePMe-seq experiments (44 % calculated from rep2 to rep1). **b**, IGV browser coverage tracks of MePMe-seq data for the indicated FURIN and PXDN mRNAs from cells labeled with PSH (purple) or methionine as control (gray). Blue and green bars indicate terminations at C and A, respectively, identified by JACUSA2. Numbers (%) denote the calculated arrest rate in that position. **c**, Frequency of m^5^C sites per transcript. **d**, Metatranscript analysis showing a density plot of the distribution of m^5^C sites detected by MePMe-seq. **e**, Consensus motif for sequences surrounding identified m^5^C sites. Representative example of n=2 biologically independent samples. **f**, Overlap of m^5^C identified in MePMe-seq (combined sites from n=2 independent experiments with HS filtering) with sites identified by bisulfite sequencing (BS-seq), improved BS-seq (Imp.), Aza-IP and miCLIP ^67^. **g**, Overlap of m^5^C sites identified in MePMe-seq (combined sites from n=2 independent experiments) with all m^5^C sites from ATLAS database, ^67^ when increasing uncertainty region around the site is applied.

Similar to sequencing data for m^6^A, reported m^5^C sites vary strongly between different reports as can be seen in a pairwise comparison of results from different base resolution techniques (Supplementary Table 5, 6), retrieved from ATLAS database ^67^. The overlap of m^5^C sites identified by MePMe-seq with reported sites is also very low. However, if an uncertainty window of ± 50 nt is allowed, 20 % of the identified sites overlap with sites from previous studies. The imperfections of different m^5^C mapping methods are a well-known problem in the field, demanding for m^5^C mapping methods independent of bisulfite treatment ^28, 79, 82^. Of note, the m^5^C sites overlapping between MePMe-seq and individual bisulfite experiments were reported in multiple experiments. For example m^5^C in NECTIN2, FURIN, TRAF7, PXDN, THOC7 and ZNF106 was present in 5–17 experiments (according to the ATLAS database ^67^), thus independently confirming their existence.

To validate the intracellular formation of prop^5^C in response to metabolic labeling with PSH, we developed an LC-QqQ-MS method to detect and quantify prop^5^C in mRNA. To this end, we synthesized prop^5^C (Supplementary Fig. 1) and determined the mass spectrometric parameters required for quantification process (Supplementary Table 7). After PSH feeding, LC-QqQ-MS analyses of mRNA confirmed formation of prop^5^C, based on the MS/MS fragmentation (MRM transition 282.2 *→* 150.1) that was used for quantification (Fig. 2f). This MS/MS fragment was absent in mRNA from control HeLa cells that had been fed with Met instead (Fig. 2f, g, Supplementary Fig. 24). Two additional MS/MS peaks (MRM transition 282.1 *→* 121.9, 282.1 *→* 80) used for independent confirmation were less pronounced (Supplementary Fig.2, for details see Supplementary Note 3). From the same cells, m^5^C levels were measured. In control cells, m^5^C/C ratio in mRNA was 16*10E-3 %, in line with previous reports ^78^, in PSH-labeled cells this ratio was reduced to 9*10E-3 % (Fig. 3f), similar to the reduction observed for m^6^A/A (Fig. 2e). The ratio of prop^5^C/C was 0.12*10E-3 %, which means that 1.4 % of m^5^C was substituted for prop^5^C (Fig. 2g). Taken together the LC-QqQ-MS data indicate that PSH feeding leads to formation of prop^5^C in addition to prop^6^A and A_prop_ and can be detected in mRNA. The data suggest that the termination bands observed opposite C in MePMe-seq indeed represent natural target sites of cytosine methyltransferases.

## Conclusions

We showed that PSH can be used for metabolic labeling to detect different methylation sites in mRNA with single nucleotide precision. The bioorthogonal propargyl group replaces 1.5–3.5 % of methyl groups in m^6^A and m^5^C, respectively and provides two key features, (i) it enables enrichment of the modified sites via click reaction with biotin and (ii) it induces termination when reverse transcription is performed on beads. MePMe-seq proved powerful for the precise assignment of m^6^A sites in mRNA even if multiple sites were in closer proximity. Importantly, it is completely independent of antibodies and sequence-searches, such as the DRACH motif that is used in many cases. MePMe-seq is therefore able to find non-DRACH motifs, such as METTL16 sites and previously unreported sites. The combined analysis of metabolic labeling and *in vitro* labeling proves suitable to assign m^6^A sites as target sites to a specific methyltransferase in cells, as demonstrated for METTL16.

MePMe-seq is the first method enabling simultaneous mapping of different types of RNA modifications. This technological advance will be important to study the interconnectivity of the different modifications. We showed this for m^6^A and m^5^C but it is conceivable that further improvements might allow to get insights into methylated uridines and guanines.

## Methods

### Chemical synthesis and characterization

PSH and SeAdoYn were synthesized as previously described^49^. Synthesis and characterization of prop^5^C is described in Supplementary Notes 1.

### *In vitro* transcription

*N*^6^-Propargyl-containing RNAs were produced by *in vitro* transcription using T7 RNA polymerase (RNAP). T7 primer was prehybridized with T7 template (10 μM each) before transcription was performed in 1× CARO buffer (120 mM Tris, 6 mM Spermidine, 0.03 % (v/v) Triton-X, 4.5 % (w/v) PEG6000, 15 mM DTT, pH 8.1), 15 mM Mg(OAc)_2_, 3.3 mM per NTP (UTP, GTP and CTP), 0.4 mM ATP (or modified ATP), 1 mg/mL T7 RNAP and 0.1 U PPase for 4 h at 37 °C. The template DNA was digested using 5 U/μL DNase I. The resulting test RNA was purified via PAGE (15 % denat. PAA gel, 1× TBE), the desired band was cut out, RNA was eluted and precipitated. For biotinylated RNA, *in vitro* transcription with *N*^6^-propargyl-ATP and PAGE purification were followed by CuAAC (150 mM biotin-N_3_, 60 mM sodium phosphate buffer, 25 mM THPTA, 5 mM CuSO_4_ (THPTA and CuSO_4_) were pre-mixed) 100 mM sodium ascorbate) and an additional PAGE purification.

### Primer extension assays for SSIII/IV

RT-primer (20 μM) was radioactively labeled with [γ-^32^P]-ATP using 0.4 U/μL T4 PNK (30min, 37°C). Excess [γ-^32^P]-ATP was removed by MicroSpin G-25 columns. The test RNA (0.3 μM) with or without *N*^6^-modified adenosine was hybridized with 2 μM [^32^P]-labeled RT-primer (5 min, 65°C followed by 5 min, 0 °C) and incubated with 0.1 U/μL reverse transcriptase (SSIII or SSIV) and 0.1 mM dNTP for 30 min. The reaction was stopped by addition of 0.1 M HClO_4_. The RNA was digested via alkaline hydrolysis by addition of 185 mM NaOH (80 °C, 10min), neutralized with HCl and analyzed via PAGE. Termination bands were quantified using ImageJ and normalized to the total amount of primer extension products. If bioconjugation via CuAAC was performed, the biotinylated RNA was bound to 400μg M-280 Streptavidine Dynabeads (Life Technologies) according to the manufacturer’s protocol and washed thoroughly before RT was performed directly on beads as described above.

### Determination of clicking efficiency

20 ng/μL test RNA were mixed with 150 mM biotin-N_3_ and 60 mM phosphate buffer. Then, 25 mM THPTA and 5 mM CuSO_4_ (pre-mixed) were added, the reaction was started by adding 100 mM NaAsc and incubated for 30 min at 37 °C. The reaction was stopped by adding 10 mM EDTA. The RNA was precipitated in EtOH, suspended in ddH_2_O and digested to nucleosides, using P1 nuclease and FastAP. For this, the RNA (140 ng/μL) was digested to nucleotides using 5 mU/μL P1 nuclease and 1× P1 nuclease buffer (1 h, 50 °C) and dephosphorylated to nucleosides by adding 0.05 U/μL FastAP and incubating for 1 h at 37 °C. 100 mM HClO_4_ was added to denature the enzymes and after centrifugation, the clear supernatant was used for HPLC or LC-MS analysis.

### Metabolic labeling of HeLa cells

HeLa cells were cultured in MEM Earl’s with 10 % (v/v) fetal bovine serum (FCS, PAN), 1 % (v/v) non-essential amino acids (NEAS, PAN) and 1 % (v/v) glutamine solution (200 mM in PBS, PAN) in a humidified incubator at 37 °C and 5 % CO_2_. The cells (10 mL of a 2*10^5^ cells·mL^-1^ suspension) were seeded into cell culture dishes (Ø = 10 cm) and cultivated for 1 d. The medium was removed and the cells were starved in Met-deficient medium (Gibco) supplemented with 1 % (v/v) glutamine solution and 1 mM L-cysteine for 30 min to deplete intracellular Met. PSH was added at different concentrations and the cells were cultivated for another 16 h. For negative controls, L-Met was added instead of PSH. The cells were washed with PBS, trypsinized and suspended in 20 mL of supplemented MEM Earl′s. The suspension was centrifuged (800 rpm, 10 min), the medium was removed, and the pellet was stored at –80 °C.

### Isolation of total RNA from HeLa cells

Cell pellets from HeLa cells (∼ 2.5–3.0 × 10^5^ cells) were lysed mechanically by pipetting in 5 mL lysis buffer (10 mM TrisHCl (pH 8.0), 150 mM NaCl, 0.5 mM NP40, adjusted to pH 7.5 with HCl). Total RNA was purified by two consecutive phenol-chloroform extractions (4:1 and 2:1) followed by extraction with 1 mL CHCl_3_, back-extraction with NaCl (0.9 %) and EtOH precipitation. The resulting total RNA was treated with 1 U DNase I (Thermo Fisher) in 1× DNase buffer (Thermo Fisher) for 30 min at 37 °C, followed by phenol-CHCl_3_ (5:1) extraction and EtOH precipitation.

### Isolation of Poly(A)+ mRNA from HeLa cells

Extraction of poly(A)^+^ RNA directly from HeLa cell pellets (1–5×10^6^ cells) was performed with Sera-Mag Oligo(dT)-Coated Magnetic Particles according to manufacturer’s instructions. Extraction was performed once for SELECT assays and twice for LC-MS/QqQ and NGS.

### Quantification of nucleic acids via spectrophotometric assay using TECAN / Nanodrop

To quantify purified nucleic acids and assess their purity, absorbance at 260 nm and 280 nm were measured with a TECAN Infinite M1000 Pro in combination with NanoQuant plate or with NanoDrop Spectrophotometer. Purity was determined by the 260/280 nm absorbance ratios.

### Quantification of nucleic acids via NanoQuant plate adapted PicoGreen assay

Concentration of NGS libraries was determined using Quant-iT™ PicoGreen™ dsDNA Assay Kit (Invitrogen) following the procedure for fluorescence-based DNA quantification in small volumes by TECAN on NanoQuant Plate™ (TECAN). Measurement was repeated trice for technical replicates from the same mix. Raw values were blank corrected and technical replicates were averaged. Concentrations were determined by comparison to lambda DNA standards.

### Digestion of total RNA / poly(A)^+^ RNA for LC-MS-QqQ analysis

Digestions of isolated RNA was performed as above for determination of clicking efficiency, using 0.1 U nuclease P1 per 1 μg RNA.

### Quantification of modified and unmodified nucleosides via LC-QqQ-MS

For quantification of required analytes, the calibration curves of external standards were measured. We used commercially available nucleosides for all standards (see Supplementary Table 11) except for 5-propargylcytidine (prop^5^C). Prop^5^C was synthesized and characterized (see Supplementary Note 1).

Quantification of modified and unmodified nucleosides was performed on a LC-QqQ-MS system (for details about instrument and software see Supplementary note 2). Digested, dephosphorylated nucleoside mix was separated with a linear gradient from buffer A (20 mM NH_4_OAc in ddH_2_O (pH 6.0) to buffer B (acetonitrile) (see Supplementary Table 6) at 40 °C with 0.8 mL/min. The principle of measuring, optimizing mass spectrometric parameters, and quantifying modified nucleosides was described before in great detail by Muthmann et al.^58^. For best performance, mass spectrometric parameters such as fragmentor voltage (FV) and collision energy (CE) were optimized individually for each product ion. The optimized MS conditions are listed in Supplementary Table 7.

Mass measurements for several nucleosides were performed in parallel in dynamic MRM mode. The retention time (RET.) of all analytes and respective RET windows are listed in Supplementary Table 7. For quantification, pairs of precursor ion and product ion were detected, using the most abundant MRM transition - fragment of ribose loss or ribose derivative loss ^58^ - as quantifier. The second most abundant or specific MRM transition was used as a qualifier and the ratio between quantifier and qualifier signals was used as extra evidence for analyte identity. In the cases of prop^6^A and prop^5^C, two qualifiers were used. The quantification of prop^5^C is described in more detail in Supplementary note 3.

### NGS library preparation

NGS libraries from mRNA were prepared using a modified version of the iCLIP2 protocol^59^. Detailed procedures can be found in Supplementary Note 4.

### Illumina sequencing

The libraries were sequenced on Illumina platform as paired-end with the read length 150 (PE150). The received amount of data per sample was 5G which corresponds to ∼33 million reads.

### Preprocessing of raw sequencing data

The quality filtering, trimming and UMI preprocessing were performed in one step with fastp software^83^. Since the first 15 bases of read 1 had the structure (UMI1)_5 nt_(Experimental barcode)_6 nt_(UMI2)_4 nt_, the entire 15-mer was treated as UMI. The barcode-based demultiplexing was performed manually with standard command-line tools, in order to retain only the reads containing the correct experimental barcode. The alignment was performed with hisat2^84^, with the disabled soft-clipping, as recommended by Busch et al.^85^. The deduplication step was performed with UMI-tools^86^ with the default directional method to define duplicates, as recommended for highly over-amplified libraries^85^.

### Termination calling via JACUSA2 and filtering

Differential reverse transcription termination signatures between treated and control samples were analyzed with our software package JACUSA2 ^62^ https://github.com/dieterich-lab/JACUSA2 using run mode “rt-arrest”. As one replicate of MePMe-seq CTR failed, MePMe-seq PSH rep1 was compared in further analysis with the libraries from untreated HeLa mRNA (METTL16 CTR). We used the following set of parameters: rt-arrest -m 0 -c 4 -p 10 -P1 FR-SECONDSTRAND -P2 FR-SECONDSTRAND, which disable quality clipping of read alignments (-m), require a minimal coverage of 4 reads across all samples (-c), run with 10 CPU threads (-p 10) and set library orientation to second strand in cDNA synthesis (-P1, -P2). JACUSA2 outputs genomic site coordinates, read coverage and termination rates per sample. The difference between arrest rates from sample and control (Δ_(RT arrest)_) was used to filter the terminations identified by the algorithm and to remove false positives. We retained high and low confident predictions by setting a sample read coverage threshold (> 35 for high stringency (HS) filter, > 20 for low stringency (LS) filter) and arrest score threshold (*Δ*_*RTarrest*_>20 for HS filter; *Δ*_*RT arrest*_>15 for LS filter). We annotated the reported sites with genomic features such as gene locus, exon class or intron, distance to next TSS in 5’direction and the sequence context using bedtools 2.29.2.

### Metatranscript analysis

Genomic coordinates were converted into cDNA coordinates using R language (version 4.0.2, https://www.R-project.org/)^87^ and the Bioconductor package ensembldb (version 2.14.1, PMID: 30689724). We always selected the longest transcript per locus to which genomic coordinates mapped to. For every protein-coding transcript, we computed the respective cDNA coordinate of a given termination site and reported the length of this matching transcript (in nt), the length of the coding sequence, five and three prime untranslated region (UTR, in nt). Subsequently, and using this information, we used the same strategy as in MetaPlotR (PMID: 28158328) to visualize the read termination site distribution along a metatranscript profile (5’UTR+CDS+3’UTR). Graphical density plots were generated using the ggplot2 library (version 3.3.2)^88^.

### SELECT

SELECT was used to independently validate selected putative m^6^A sites identified via MePMe-seq. Experiments were performed with poly(A)^+^ RNA (one time enriched via oligo(dT) magnetic beads) from HeLa cells. RNA was split and methylations were removed by treatment with FTO (+ FTO) from one half but not the other (-FTO) in 50 mM MES-buffer (pH 5.5), 283 μM (NH_4_)_2_Fe(SO_4_)_2_, 300μM α-ketoglutarate, 2 mM ascorbic acid, 50 ng/μL poly(A)^+^ RNA, 0.2 U/μL RiboLock, 0.1 μg/μL FTO (+FTO) or ddH2O (-FTO) for 30 min at 37°C. Reaction was stopped by adding 20 mM EDTA (95°C, 5 min)^71^. The poly(A)^+^ RNAs were purified immediately via RNA clean & concentrator kit by Zymo Research. For the SELECT assay poly(A)^+^ RNA (+ FTO and – FTO, respectively) was mixed with 85 pmol dTTP, 1× CutSmart buffer (50 mM potassium acetate, 20 mM Tris-acetate, 10 mM magnesium acetate,100 μg/mL BSA, pH 7.9, NEB), 20 U RiboLock, 80 fmol up primer and 80 fmol down primer in 15 μL ddH_2_O. Depending on the transcript expression level either 50 ng or 100 ng of + FTO and – FTO poly(A)^+^ RNA were used. The primer were annealed to the RNA (90 °C for 1 min; –10 °C/min for 4 min; 40 °C for 6 min; keep at 4 °C) and the enzyme mixture (0.01 U Bst 2.0 DNA polymerase, 0.5 U SplintR ligase, 10 nmol ATP and 1× CutSmart buffer in 5 μL ddH_2_O) was added. The reaction was carried out at 40 °C for 20 min, denatured at 80 °C for 20 min and kept at 4 °C. For the qPCR 1 μL of the reaction mixture was added to 9 μL of the *PowerUp™ SYBR*^*®*^ *Green Master Mix*. The data was analyzed with Bio-Rad CFX Maestro 1.0 software. The obtained results were normalized by N-site.

### Recombinant production of MTAN

Recombinant MTAN was expressed as described previously^89^.

### Recombinant production of FTO

For the recombinant production of FTO, *E. coli* BL21 (DE3) cells were transformed with a pET28a vector encoding FTO. The cells were grown in LB medium at 37 °C to an OD_600_ of 1, kept at room temperature for 30 min, induced by adding 1 mM IPTG followed by expression overnight (16 °C). The cells were lysed in binding buffer (50 mM sodium phosphate buffer (pH 8), 300 mM NaCl, 50 mM imidazole) and the enzyme was purified in two steps using the ÄKTApurifier system. In a first purification step, the lysate was loaded on a 1 mL HisTrap FF column. The column was washed with 4 % elution buffer (50 mM sodium phosphate buffer (pH 8), 300 mM NaCl, 500 mM imidazole) in binding buffer at 1 mL/min in 6 column volumes (CV), followed by a linear gradient from 4–100 % elution buffer in 10 CV. The gradient was held at 21 % elution buffer when FTO started to elute and then continued the linear gradient to 100 %. In a second step, the enzyme was purified using a Superdex 200 increase column with running buffer (50 mM sodium phosphate buffer (pH 8), 300 mM NaCl, 10 % glycerol) at 0.45 mL/min.

### Recombinant expression and purification of GST-METTL16

Recombinant GST-METTL16 was expressed as described previously^90^. For construct refer to Supplementary Fig. 55. For RNA-free preparation, an anion exchange purification step was added (HiTrap Q, GE Healthcare).

### *In vitro* propargylation of mRNA with METTL16

To propargylate isolated poly(A)^+^-RNA, 4 μg of RNA were incubated in 30 μL volume with 1 mM SeAdoYn, 10 μM METTL16 (RNA free), 1 U/ μL Ribolock and 0.4 μM MTAN in 10× METTL16 activity buffer (100 mM HEPES-KOH, pH 7.4, 1 M NaCl) in 30 μL for 1 h at 37 °C. RNA was purified with RNA Clean & Concentrator™-5 kit (Zymo Research Europe GmbH). 100 ng of reaction product were used for analysis on a denaturing agarose gel.

### Data acquisition

For comparison with MePMe-seq sites m^6^A, Nm and m^5^C datasets were downloaded from ATLAS^67^ and REPIC^66^ database. Additional datasets were acquired from the respective publications^46, 48^. Coordinates that were not in hg38 were converted using browser based LiftOver tool (available at https://genome.ucsc.edu/cgi-bin/hgLiftOver ^91^).

## Acknowledgements

A.R. thanks the DFG for funding within the priority programme SPP1784 (RE 2796/3-2). A. O. was supported by a CiM-IMPRS fellowship. At the WWU Münster, we would like to thank Ann-Marie Lawrence Dörner and the analytical facilities of the Organic Chemistry Institute for excellent technical assistance. We would like to thank students involved in the project for technical assistance, namely Fabrice Becker, Wiebke Teich and Greta Charlotte Dahm. Group members Dr. Nils Muthmann, Nils Klöcker, Melissa van Dülmen and Florian Weissenböck are acknowledged for fruitful discussions. Dr. Julian König (IMB Mainz) is gratefully acknowledged for sharing his library preparation protocol and for technical tips. We thank Prof. Chengqui Yi (Peking University) for providing the plasmid for FTO expression.

## Author contributions

K.H., A.O. and A.R. conceived the project. K.H. designed, optimized and performed MePMe-seq. A.O. designed and performed *in vitro* labeling with METTL16. K.H. and N.A.K. prepared sequencing libraries. N.A.K. and K.H. designed, performed and analyzed SELECT experiments. P.S. designed and performed chemical syntheses of prop^5^C. A.O., N.A.K. and N.V.C. performed chemical syntheses of PSH and SeAdoYn. P.S. developed, optimized and performed LC-QqQ-MS measurements with contributions of K.H.. S.H. performed cell culture experiments. C.D., A.O. and K.H. analyzed and evaluated NGS data and performed statistical analyses. A.R. supervised the project and contributed to the design of experiments. All authors discussed the results. A.R. and K.H. wrote the manuscript with contributions from all coauthors. All authors read and approved the final manuscript.

## Data availabilitiy

Mapped data for both replicates of MePMe-seq and in vitro METTL16 labeling in HeLa cells will be available from the NCBI Sequence Read Archive (SRA) after publication or can be provided for review upon request. Processed data, including JACUSA2 sites from both replicates of MePMe-seq (Supplementary data 1), identified m^6^A sites from MePMe-seq (Supplementary data 2), identified m^5^C sites from MePMe-seq (Supplementary data 6), JACUSA2 sites from both replicates of in vitro METTL16 labeling (Supplementary data 4) and identified m^6^A sites from *in vitro* METTL16 labeling (Supplementary data 5) are provided as Supplementary data.

## Code availability

JACUSA2 is available at https://github.com/dieterich-lab/JACUSA2.

## Supplementary information

Supplementary information (tables 1-11, Figs. 1-55, Notes 1-4)

Supplementary data 1-7

