## Supplementary Material for "MePMe-seq: Antibody-free simultaneous m^6^A and m^5^C mapping in mRNA by metabolic propargyl labeling and sequencing"

Supplementary Figure 38: COSY spectrum of compound 3. ... 53
Supplementary Figure 39: HSQC spectrum of compound 3. ... 54
Supplementary Figure 40: HMBC spectrum of compound 3. ... 55
Supplementary Figure 41: $^{1}$H-NMR spectrum of compound 4. ... 56
Supplementary Figure 42: $^{13}$C-NMR spectrum of compound 4. ... 57
Supplementary Figure 43: COSY spectrum of compound 4. ... 58
Supplementary Figure 44: HSQC spectrum of compound 4. ... 59
Supplementary Figure 45: HMBC spectrum of compound 4. ... 60
Supplementary Figure 46: $^{1}$H-NMR spectrum of compound 5. ... 61
Supplementary Figure 47: $^{13}$C-NMR spectrum of compound 5. ... 62
Supplementary Figure 48: COSY spectrum of compound 5. ... 63
Supplementary Figure 49: HSQC spectrum of compound 5. ... 64
Supplementary Figure 50: HMBC spectrum of compound ... 65
Supplementary Figure 52: The representation of all points of calibration (cc1 to cc7) curve of prop$^{5}$C. ... 67
Supplementary Figure 53: The signal intensity of different MRM transition ions of prop$^{5}$C. ... 68
Supplementary Figure 54: Quantifier peaks of calibration curve of prop$^{5}$C ... 69
Supplementary Figure 55: Sequence of METTL16-construct used in this work. ... 70
Supplementary Notes ... 71
Supplementary Note 1: Synthesis of 5-propargylcytidine (prop$^{5}$C) ... 71
Supplementary Note 2: Details about the LC-QqQ-MS instrument and software setup ... 75
Supplementary Note 3: Details to prop$^{5}$C quantification via LC-QqQ-MS ... 76
Supplementary Note 4: NGS library preparation for MePMe-seq ... 77
Supplementary References ... 79

Supplementary Tables:

Supplementary Table 1: Number of reads after NGS sequencing and details from NGS data processing. Libraries were prepared from mRNA isolated from HeLa labeled with 2.5 mM PSH (MePMe-seq PSH) or 2.5 mM Met (MePMe-seq CTR) or mRNA isolated from untreated HeLa cells (METTL16 CTR) and labeled *in vitro* using METTL16 (METTL16 SAMPLE). MePMe-seq PSH rep1 was compared in further analysis with the libraries from untreated HeLa mRNA (METTL16 CTR).

| Sample | MePMe-seq PSH rep1 | MePMe-seq PSH rep2 | MePMe-seq CTR rep2 | METTL16 SAMPLE rep 1 | METTL16 SAMPLE rep 2 | METTL16 CTR rep1 | METTL16 CTR rep2 |
|---|---|---|---|---|---|---|---|
| Barcode | ATTGGC | GCTCAT | AAGCTA | TACAAG | ACATCG | AAGCTA | CGTGAT |
| Total read pairs | 27,247,772 | 26,521,975 | 19,459,453 | 28,723,676 | 26,608,786 | 28,877,411 | 20,738,712 |
| Good quality reads (FASTQ) | 96.37 % | 93.30 % | 80.52 % | 88.30 % | 94.13 % | 92.40 % | 92.96 % |
| Alignment rate | 94.41 % | 89.38 % | 88.00 % | 92.99 % | 88.84 % | 94.66 % | 89.51 % |
| Mapped reads removed in deduplication | 61.55 % | 46.62 % | 68.03 % | 71.34 % | 84.81 % | 58.97 % | 75.32 % |
| Reads after UMI deduplication | 14,782,330 | 15,463,622 | 4,658,002 | 7,167,391 | 3,430,028 | 10,590,131 | 3,592,481 |

Supplementary Table 2: Pairwise comparison of miCLIP experiments from human samples. Shown are the numbers of overlapping identified $m^6A$ sites between the indicated experiments and the corresponding percentage (referred to the total number of sites in the respective experiment).

| Experiment | | 31279658 HEK293T Control | 30867593 HepG2 Control | 28920958 MOLM13 Control | 26593424 HepG2 Control | 26121403 HEK293 sysy_antinbody_CITS | 26121403 HEK293 Control | 31328227 HCT116 Control |
|---|---|---|---|---|---|---|---|---|
| 31279658 HEK293T Control | sites | 43652 | 518 | 6633 | 710 | 3450 | 3454 | 2514 |
| | % of all | 100.0 | 1.2 | 15.2 | 1.6 | 7.9 | 7.9 | 5.8 |
| 30867593 HepG2 Control | sites | 518 | 3424 | 227 | 79 | 105 | 137 | 109 |
| | % of all | 15.1 | 100.0 | 6.6 | 2.3 | 3.1 | 4.0 | 3.2 |
| 28920958 MOLM13 Control | sites | 6633 | 227 | 20516 | 488 | 1599 | 1528 | 1217 |
| | % of all | 32.3 | 1.1 | 100.0 | 2.4 | 7.8 | 7.4 | 5.9 |
| 26593424 HepG2 Control | sites | 710 | 79 | 488 | 12208 | 160 | 273 | 296 |
| | % of all | 5.8 | 0.6 | 4.0 | 100.0 | 1.3 | 2.2 | 2.4 |
| 26121403 HEK293 sysy_antinbody_CITS | sites | 3450 | 105 | 1599 | 160 | 6215 | 957 | 588 |
| | % of all | 55.5 | 1.7 | 25.7 | 2.6 | 100.0 | 15.4 | 9.5 |
| 26121403 HEK293 Control | sites | 3454 | 137 | 1528 | 273 | 957 | 7370 | 706 |
| | % of all | 46.9 | 1.9 | 20.7 | 3.7 | 13.0 | 100.0 | 9.6 |
| 31328227 HCT116 Control | sites | 2514 | 109 | 1217 | 296 | 588 | 706 | 6368 |
| | % of all | 39.5 | 1.7 | 19.1 | 4.6 | 9.2 | 11.1 | 100.0 |

Supplementary Table 3: Pairwise comparison of m6A sequencing techniques with base resolution. Shown are the numbers of overlapping identified m6A sites between the indicated techniques and the corresponding percentage (referred to the total number of sites in the respective technique). Results from multiple experiments using the same technique on human tissues/cell types were pooled.

| Technique | | MePMe-seq (HS) | m6A-label-seq | DART-seq | m6A-REF-seq | PA- m6A seq | m6A-seq (improved protocol) | m6A CLIP (human samples) | miCLIP (human samples) |
|---|---|---|---|---|---|---|---|---|---|
| MePMe-seq (HS) | sites | 4502 | 359 | 18 | 373 | 679 | 612 | 2505 | 2941 |
| | % of all | 100.0 | 8.0 | 0.4 | 8.3 | 15.1 | 13.6 | 55.6 | 65.3 |
| m6A-label-seq | sites | 359 | 2497 | 8 | 89 | 110 | 159 | 717 | 805 |
| | % of all | 14.4 | 100.0 | 0.3 | 3.6 | 4.4 | 6.4 | 28.7 | 32.2 |
| DART-seq | sites | 18 | 8 | 8543 | 167 | 309 | 29 | 881 | 1318 |
| | % of all | 0.2 | 0.1 | 100.0 | 2.0 | 3.6 | 0.3 | 10.3 | 15.4 |
| m6A-REF-seq | sites | 373 | 89 | 167 | 11474 | 448 | 295 | 1657 | 2309 |
| | % of all | 3.3 | 0.8 | 1.5 | 100.0 | 3.9 | 2.6 | 14.4 | 20.1 |
| PA- m6A -seq | sites | 679 | 158 | 309 | 448 | 19687 | 828 | 3467 | 5309 |
| | % of all | 3.4 | 0.8 | 1.6 | 2.3 | 100.0 | 4.2 | 17.6 | 27.0 |
| m6A -seq improved (pooled human samples) | sites | 612 | 159 | 29 | 295 | 828 | 10741 | 3944 | 5133 |
| | % of all | 5.7 | 1.5 | 0.3 | 2.7 | 7.7 | 100.0 | 36.7 | 47.8 |
| m6A CLIP (pooled human samples) | sites | 2505 | 717 | 881 | 1657 | 3467 | 3944 | 50092 | 22701 |
| | % of all | 5.0 | 1.4 | 1.8 | 3.3 | 6.9 | 7.9 | 100.0 | 45.3 |
| miCLIP (pooled human samples) | sites | 2941 | 805 | 1318 | 2309 | 5309 | 5133 | 22701 | 80104 |
| | % of all | 3.7 | 1.0 | 1.6 | 2.9 | 6.6 | 6.4 | 28.3 | 100.0 |

Supplementary Table 4: List of $m^6A$ sites for representative transcripts found via *in vitro* METTL16 labeling and comparison with MePMe-seq. Shown are sites in MALAT1, MYC, NT5DC2 and RBM transcripts that were found in both replicates of either technique.

| coordinates (hg38) | | | | | | Sequence context | | | | | | | | |
|---|---|---|---|---|---|---|---|---|---|---|---|---|---|---|
| arrest Chr | arrest Start | arrest End | Strand | modification | gene | -2 | -1 | 0 | 1 | 2 | METTL16 rep1 HS | METTL16 rep2 HS | MePMe-seq rep1 HS | MePMe-seq rep2 HS |
| 11 | 65503012 | 65503013 | + | 65503012 | MALAT1 | A | C | A | C | T | true | true | | |
| 11 | 65502223 | 65502224 | + | 65502223 | MALAT1 | A | C | A | A | T | true | true | | |
| 11 | 65502935 | 65502936 | + | 65502935 | MALAT1 | A | C | A | A | T | true | true | | |
| 11 | 65505045 | 65505046 | + | 65505045 | MALAT1 | A | C | A | T | A | true | true | | |
| 11 | 65503390 | 65503391 | + | 65503390 | MALAT1 | A | C | A | C | T | true | true | | |
| 11 | 65501369 | 65501370 | + | 65501369 | MALAT1 | A | C | A | G | T | true | true | | |
| 11 | 65504575 | 65504576 | + | 65504575 | MALAT1 | A | C | A | T | T | true | true | | |
| 11 | 65499371 | 65499372 | + | 65499371 | MALAT1 | A | C | A | G | A | true | true | | |
| 11 | 65504068 | 65504069 | + | 65504068 | MALAT1 | A | C | A | C | T | true | true | | |
| 11 | 65504324 | 65504325 | + | 65504324 | MALAT1 | A | C | A | G | A | true | true | | |
| 11 | 65504814 | 65504815 | + | 65504814 | MALAT1 | A | C | A | T | G | true | true | | |
| 8 | 127741111 | 127741112 | + | 127741111 | MYC | G | G | A | C | T | | | true | true |
| 8 | 127740891 | 127740892 | + | 127740891 | MYC | G | G | A | C | T | | | true | true |
| 8 | 127740621 | 127740622 | + | 127740621 | MYC | G | G | A | C | A | | | true | true |
| 8 | 127738657 | 127738658 | + | 127738657 | MYC | G | G | A | C | T | | | true | true |
| 8 | 127736472 | 127736473 | + | 127736472 | MYC | A | C | A | A | C | true | true | | |
| 8 | 127740845 | 127740846 | + | 127740845 | MYC | A | C | A | T | C | true | true | | |
| 8 | 127741209 | 127741210 | + | 127741209 | MYC | A | C | A | C | A | true | true | | |
| 3 | 52529150 | 52529151 | - | 52529152 | NT5DC2 | A | C | A | A | G | true | true | | |
| 3 | 52528192 | 52528193 | - | 52528194 | NT5DC2 | A | C | A | A | G | true | true | | |
| 3 | 52528903 | 52528904 | - | 52528905 | NT5DC2 | A | C | A | A | C | true | true | | |
| X | 48577066 | 48577067 | + | 48577066 | RBM3 | A | C | A | G | A | true | true | | |

Supplementary Table 5: Pairwise comparison of m$^5$C sequencing experiments. Shown are the numbers of overlapping identified m$^5$C sites between the indicated experiments and the corresponding percentage (referred to the total number of sites in the respective experiment). Results from experiments in the same study on human tissues/cell types were pooled.

| Experiment | | 23604283; Aza-IP | 27356879 miCLIP | 22344696 BS-seq | 27356879 BS-seq | 28418038 (all) BS-seq | 30526041 BS-seq | 30872485 RBS seq | 31061524 (all) BS-seq | 31358969 BS-seq | MePMe-seq (HS) |
|---|---|---|---|---|---|---|---|---|---|---|---|
| 23604283; Aza-IP | sites | 597 | 4 | 2 | 12 | 5 | 119 | 4 | 1 | 0 | 0 |
| | % of all | 100.0 | 0.7 | 0.3 | 2.0 | 0.8 | 19.9 | 0.7 | 0.2 | 0.0 | 0.0 |
| 27356879 miCLIP | sites | 4 | 361 | 1 | 86 | 361 | 37 | 0 | 0 | 94 | 0 |
| | % of all | 1.1 | 100.0 | 0.3 | 23.8 | 100.0 | 10.2 | 0.0 | 0.0 | 26.0 | 0.0 |
| 22344696 BS-seq | sites | 2 | 1 | 5246 | 5 | 342 | 16 | 118 | 81 | 289 | 2 |
| | % of all | 0.0 | 0.0 | 100.0 | 0.1 | 6.5 | 0.3 | 2.2 | 1.5 | 5.5 | 0.0 |
| 27356879 BS-seq | sites | 12 | 86 | 5 | 2357 | 94 | 73 | 5 | 2 | 212 | 0 |
| | % of all | 0.5 | 3.6 | 0.2 | 100.0 | 4.0 | 3.1 | 0.2 | 0.1 | 9.0 | 0.0 |
| 28418038 (all) BS-seq | sites | 5 | 361 | 342 | 94 | 40554 | 65 | 495 | 1040 | 4397 | 22 |
| | % of all | 0.0 | 0.9 | 0.8 | 0.2 | 100.0 | 0.2 | 1.2 | 2.6 | 10.8 | 0.1 |
| 30526041 BS-seq | sites | 119 | 37 | 16 | 73 | 65 | 12442 | 14 | 2 | 82 | 0 |
| | % of all | 1.0 | 0.3 | 0.1 | 0.6 | 0.5 | 100.0 | 0.1 | 0.0 | 0.7 | 0.0 |
| 30872485 RBS seq | sites | 4 | 0 | 118 | 5 | 495 | 14 | 2159 | 161 | 327 | 2 |
| | % of all | 0.2 | 0.0 | 5.5 | 0.2 | 22.9 | 0.6 | 100.0 | 7.5 | 15.1 | 0.1 |
| 31061524 (all) BS-seq | sites | 1 | 0 | 81 | 2 | 1040 | 2 | 161 | 3096 | 503 | 7 |
| | % of all | 0.0 | 0.0 | 2.6 | 0.1 | 33.6 | 0.1 | 5.2 | 100.0 | 16.2 | 0.2 |
| 31358969 BS-seq | sites | 0 | 94 | 289 | 212 | 4397 | 82 | 327 | 503 | 20488 | 8 |
| | % of all | 0.0 | 0.5 | 1.4 | 1.0 | 21.5 | 0.4 | 1.6 | 2.5 | 100.0 | 0.0 |
| MepMe-seq (HS) | sites | 0 | 0 | 2 | 0 | 22 | 0 | 2 | 7 | 8 | 1305 |
| | % of all | 0.0 | 0.0 | 0.2 | 0.0 | 1.7 | 0.0 | 0.2 | 0.5 | 0.6 | 100.0 |

Supplementary Table 6: Pairwise comparison of bisulfide sequencing experiments from human samples. Shown are the numbers of overlapping identified m$^5$C sites between the indicated experiments and the corresponding percentage (referred to the total number of sites in the respective experiment). Blue regions mark experiments performed in the same study but on different tissues/cell types, orange regions mark experiments performed on HeLa cells in different studies.

| Experiment | | 22344696 | 27356879 GSE66011 | 28418038 GSE74771 | 28418038 GSE74772 | 28418038 GSE74773 | 28418038 GSE74774 | 28418038 GSE74775 | 28418038 GSE93752 | 30526041 GSE122413 | 30872485 GSE90963 | 31061524 GSE122260 | 31061524 GSE122261 | 31061524 GSE122263 | 31061524 GSE122264 | 31061524 GSE122265 | 31061524 GSE122266 | 31061524 GSE122267 | 31061524 GSE122268 | 31061524 GSE122269 | 31358969 GSE133672 |
|---|---|---|---|---|---|---|---|---|---|---|---|---|---|---|---|---|---|---|---|---|---|
| 22344696 | sites | 5246 | 5 | 140 | 159 | 130 | 137 | 150 | 259 | 16 | 118 | 24 | 68 | 18 | 11 | 20 | 17 | 20 | 7 | 44 | 289 |
| | % of all | 100.0 | 0.1 | 2.7 | 3.0 | 2.5 | 2.6 | 2.9 | 4.9 | 0.3 | 2.2 | 0.5 | 1.3 | 0.3 | 0.2 | 0.4 | 0.3 | 0.4 | 0.1 | 0.8 | 5.5 |
| 27356879 GSE66011 | sites | 5 | 2357 | 88 | 9 | 3 | 4 | 5 | 3 | 73 | 5 | 2 | 1 | 2 | 2 | 2 | 2 | 2 | 2 | 2 | 212 |
| | % of all | 0.2 | 100.0 | 3.7 | 0.4 | 0.1 | 0.2 | 0.2 | 0.1 | 3.1 | 0.2 | 0.1 | 0.0 | 0.1 | 0.1 | 0.1 | 0.1 | 0.1 | 0.1 | 0.1 | 9.0 |
| 28418038 GSE74771 | sites | 140 | 88 | 11235 | 3851 | 3078 | 2856 | 3196 | 1817 | 48 | 210 | 149 | 298 | 111 | 51 | 84 | 84 | 57 | 53 | 177 | 1990 |
| | % of all | 1.2 | 0.8 | 100.0 | 34.3 | 27.4 | 25.4 | 28.4 | 16.2 | 0.4 | 1.9 | 1.3 | 2.7 | 1.0 | 0.5 | 0.7 | 0.7 | 0.5 | 0.5 | 1.6 | 17.7 |
| 28418038 GSE74772 | sites | 159 | 9 | 3851 | 13803 | 3202 | 3012 | 3942 | 2144 | 9 | 238 | 133 | 276 | 116 | 46 | 78 | 78 | 55 | 41 | 176 | 2353 |
| | % of all | 1.2 | 0.1 | 27.9 | 100.0 | 23.2 | 21.8 | 28.6 | 15.5 | 0.1 | 1.7 | 1.0 | 2.0 | 0.8 | 0.3 | 0.6 | 0.6 | 0.4 | 0.3 | 1.3 | 17.0 |
| 28418038 GSE74773 | sites | 130 | 3 | 3078 | 3202 | 9367 | 2646 | 2940 | 1703 | 15 | 227 | 117 | 274 | 109 | 50 | 84 | 83 | 56 | 48 | 159 | 1797 |
| | % of all | 1.4 | 0.0 | 32.9 | 34.2 | 100.0 | 28.2 | 31.4 | 18.2 | 0.2 | 2.4 | 1.2 | 2.9 | 1.2 | 0.5 | 0.9 | 0.9 | 0.6 | 0.5 | 1.7 | 19.2 |
| 28418038 GSE74774 | sites | 137 | 4 | 2856 | 3012 | 2646 | 8770 | 2945 | 1688 | 9 | 228 | 84 | 217 | 85 | 46 | 82 | 54 | 53 | 39 | 125 | 1848 |
| | % of all | 1.6 | 0.0 | 32.6 | 34.3 | 30.2 | 100.0 | 33.6 | 19.2 | 0.1 | 2.6 | 1.0 | 2.5 | 1.0 | 0.5 | 0.9 | 0.6 | 0.6 | 0.4 | 1.4 | 21.1 |
| 28418038 GSE74775 | sites | 150 | 5 | 3196 | 3942 | 2940 | 2945 | 11220 | 2059 | 7 | 291 | 119 | 383 | 123 | 69 | 111 | 90 | 91 | 57 | 227 | 2261 |
| | % of all | 1.3 | 0.0 | 28.5 | 35.1 | 26.2 | 26.2 | 100.0 | 18.4 | 0.1 | 2.6 | 1.1 | 3.4 | 1.1 | 0.6 | 1.0 | 0.8 | 0.8 | 0.5 | 2.0 | 20.2 |
| 28418038 GSE93752 | sites | 259 | 3 | 1817 | 2144 | 1703 | 1688 | 2059 | 6734 | 7 | 404 | 160 | 681 | 134 | 74 | 133 | 116 | 100 | 66 | 278 | 2488 |
| | % of all | 3.8 | 0.0 | 27.0 | 31.8 | 25.3 | 25.1 | 30.6 | 100.0 | 0.1 | 6.0 | 2.4 | 10.1 | 2.0 | 1.1 | 2.0 | 1.7 | 1.5 | 1.0 | 4.1 | 36.9 |
| 30526041 GSE122413 | sites | 16 | 73 | 48 | 9 | 15 | 9 | 7 | 7 | 12442 | 14 | 2 | 2 | 2 | 2 | 2 | 2 | 2 | 2 | 2 | 82 |
| | % of all | 0.1 | 0.6 | 0.4 | 0.1 | 0.1 | 0.1 | 0.1 | 0.1 | 100.0 | 0.1 | 0.0 | 0.0 | 0.0 | 0.0 | 0.0 | 0.0 | 0.0 | 0.0 | 0.0 | 0.7 |
| 30872485 GSE90963 | sites | 118 | 5 | 210 | 238 | 227 | 228 | 291 | 404 | 14 | 2159 | 57 | 144 | 48 | 38 | 57 | 40 | 34 | 34 | 86 | 327 |
| | % of all | 5.5 | 0.2 | 9.7 | 11.0 | 10.5 | 10.6 | 13.5 | 18.7 | 0.6 | 100.0 | 2.6 | 6.7 | 2.2 | 1.8 | 2.6 | 1.9 | 1.6 | 1.6 | 4.0 | 15.1 |

| Experiment | | 22344696 | 27356879 GSE66011 | 28418038 GSE74771 | 28418038 GSE74772 | 28418038 GSE74773 | 28418038 GSE74774 | 28418038 GSE74775 | 28418038 GSE93752 | 30526041 GSE122413 | 30872485 GSE90963 | 31061524 GSE122260 | 31061524 GSE122261 | 31061524 GSE122263 | 31061524 GSE122264 | 31061524 GSE122265 | 31061524 GSE122266 | 31061524 GSE122267 | 31061524 GSE122268 | 31061524 GSE122269 | 31358969 GSE133672 |
|---|---|---|---|---|---|---|---|---|---|---|---|---|---|---|---|---|---|---|---|---|---|
| 31061524 GSE122260 | sites | 24 | 2 | 149 | 133 | 117 | 84 | 119 | 160 | 2 | 57 | 257 | 187 | 108 | 46 | 86 | 74 | 49 | 46 | 136 | 128 |
| | % of all | 9.3 | 0.8 | 58.0 | 51.8 | 45.5 | 32.7 | 46.3 | 62.3 | 0.8 | 22.2 | 100.0 | 72.8 | 42.0 | 17.9 | 33.5 | 28.8 | 19.1 | 17.9 | 52.9 | 49.8 |
| 31061524 GSE122261 | sites | 68 | 1 | 298 | 276 | 274 | 217 | 383 | 681 | 2 | 144 | 187 | 1232 | 180 | 89 | 164 | 131 | 125 | 81 | 324 | 410 |
| | % of all | 5.5 | 0.1 | 24.2 | 22.4 | 22.2 | 17.6 | 31.1 | 55.3 | 0.2 | 11.7 | 15.2 | 100.0 | 14.6 | 7.2 | 13.3 | 10.6 | 10.1 | 6.6 | 26.3 | 33.3 |
| 31061524 GSE122263 | sites | 18 | 2 | 111 | 116 | 109 | 85 | 123 | 134 | 2 | 48 | 108 | 180 | 514 | 68 | 115 | 77 | 67 | 63 | 166 | 136 |
| | % of all | 3.5 | 0.4 | 21.6 | 22.6 | 21.2 | 16.5 | 23.9 | 26.1 | 0.4 | 9.3 | 21.0 | 35.0 | 100.0 | 13.2 | 22.4 | 15.0 | 13.0 | 12.3 | 32.3 | 26.5 |
| 31061524 GSE122264 | sites | 11 | 2 | 51 | 46 | 50 | 46 | 69 | 74 | 2 | 38 | 46 | 89 | 68 | 218 | 84 | 80 | 104 | 55 | 89 | 69 |
| | % of all | 5.0 | 0.9 | 23.4 | 21.1 | 22.9 | 21.1 | 31.7 | 33.9 | 0.9 | 17.4 | 21.1 | 40.8 | 31.2 | 100.0 | 38.5 | 36.7 | 47.7 | 25.2 | 40.8 | 31.7 |
| 31061524 GSE122265 | sites | 20 | 2 | 84 | 78 | 84 | 82 | 111 | 133 | 2 | 57 | 86 | 164 | 115 | 84 | 351 | 94 | 87 | 94 | 167 | 115 |
| | % of all | 5.7 | 0.6 | 23.9 | 22.2 | 23.9 | 23.4 | 31.6 | 37.9 | 0.6 | 16.2 | 24.5 | 46.7 | 32.8 | 23.9 | 100.0 | 26.8 | 24.8 | 26.8 | 47.6 | 32.8 |
| 31061524 GSE122266 | sites | 17 | 2 | 84 | 78 | 83 | 54 | 90 | 116 | 2 | 40 | 74 | 131 | 77 | 80 | 94 | 381 | 84 | 65 | 137 | 80 |
| | % of all | 4.5 | 0.5 | 22.0 | 20.5 | 21.8 | 14.2 | 23.6 | 30.4 | 0.5 | 10.5 | 19.4 | 34.4 | 20.2 | 21.0 | 24.7 | 100.0 | 22.0 | 17.1 | 36.0 | 21.0 |
| 31061524 GSE122267 | sites | 20 | 2 | 57 | 55 | 56 | 53 | 91 | 100 | 2 | 34 | 49 | 125 | 67 | 104 | 87 | 84 | 369 | 56 | 124 | 76 |
| | % of all | 5.4 | 0.5 | 15.4 | 14.9 | 15.2 | 14.4 | 24.7 | 27.1 | 0.5 | 9.2 | 13.3 | 33.9 | 18.2 | 28.2 | 23.6 | 22.8 | 100.0 | 15.2 | 33.6 | 20.6 |
| 31061524 GSE122268 | sites | 7 | 2 | 53 | 41 | 48 | 39 | 57 | 66 | 2 | 34 | 46 | 81 | 63 | 55 | 94 | 65 | 56 | 276 | 96 | 62 |
| | % of all | 2.5 | 0.7 | 19.2 | 14.9 | 17.4 | 14.1 | 20.7 | 23.9 | 0.7 | 12.3 | 16.7 | 29.3 | 22.8 | 19.9 | 34.1 | 23.6 | 20.3 | 100.0 | 34.8 | 22.5 |
| 31061524 GSE122269 | sites | 44 | 2 | 177 | 176 | 159 | 125 | 227 | 278 | 2 | 86 | 136 | 324 | 166 | 89 | 167 | 137 | 124 | 96 | 1162 | 226 |
| | % of all | 3.8 | 0.2 | 15.2 | 15.1 | 13.7 | 10.8 | 19.5 | 23.9 | 0.2 | 7.4 | 11.7 | 27.9 | 14.3 | 7.7 | 14.4 | 11.8 | 10.7 | 8.3 | 100.0 | 19.4 |
| 31358969 GSE133672 | sites | 289 | 212 | 1990 | 2353 | 1797 | 1848 | 2261 | 2488 | 82 | 327 | 128 | 410 | 136 | 69 | 115 | 80 | 76 | 62 | 226 | 20488 |
| | % of all | 1.4 | 1.0 | 9.7 | 11.5 | 8.8 | 9.0 | 11.0 | 12.1 | 0.4 | 1.6 | 0.6 | 2.0 | 0.7 | 0.3 | 0.6 | 0.4 | 0.4 | 0.3 | 1.1 | 100.0 |

Supplementary Table 7: MRM transition ions and optimized parameters used for the LC-MS analysis of nucleosides. MRM was conducted in positive mode.

| Analytes | MRM transition ions [m/z] | type | Method | RT [min] | RT windows [min] | FV [V] | CE [V] | CAV [V] |
|---|---|---|---|---|---|---|---|---|
| A | 268.1 → 136.0 | Quantifier | 1 | 3.9 | 0.6 | 100 | 9 | 9 |
| A | 268.1 → 119.0 | Qualifier | 1 | 3.9 | 0.6 | 100 | 45 | 9 |
| C | 244.0 → 111.9 | Quantifier | 1 | 2.2 | 0.7 | 85 | 5 | 9 |
| C | 244.0 → 95.1 | Qualifier | 1 | 2.2 | 0.7 | 85 | 41 | 9 |
| $A_m$ | 282.1 → 136.1 | Quantifier | 1 | 4.17 | 0.6 | 109 | 9 | 9 |
| $A_m$ | 282.1 → 69.1 | Qualifier | 1 | 4.17 | 0.6 | 109 | 25 | 9 |
| $m^6A$ | 282.1 → 150.1 | Quantifier | 1 | 4.29 | 0.7 | 120 | 13 | 9 |
| $m^6A$ | 282.1 → 108.0 | Qualifier | 1 | 4.29 | 0.7 | 120 | 65 | 9 |
| $A_{prop}$ | 306.1 → 136.1 | Quantifier | 2 | 4.4 | 0.5 | 109 | 9 | 9 |
| $A_{prop}$ | 306.1 → 136.1 | Qualifier | 2 | 4.4 | 0.5 | 109 | 49 | 9 |
| $prop^6A$ | 306.1 → 174.0 | Quantifier | 2 | 4.7 | 0.6 | 121 | 9 | 9 |
| $prop^6A$ | 306.1 → 148.0 | Qualifier | 2 | 4.7 | 0.6 | 121 | 33 | 9 |
| $prop^6A$ | 306.1 → 108.0 | Qualifier | 2 | 4.7 | 0.6 | 121 | 57 | 9 |
| $m^5C$ | 258.1 → 126.0 | Quantifier | 2 | 3.5 | 0.4 | 96 | 4 | 9 |
| $m^5C$ | 258.1 → 108.9 | Qualifier | 2 | 3.5 | 0.4 | 104 | 15 | 9 |
| $prop^5C$ | 282.1 → 150.1 | Quantifier | 2 | 3.9 | 0.4 | 76 | 0 | 9 |
| $prop^5C$ | 282.1 → 121.9 | Qualifier | 2 | 3.9 | 0.4 | 100 | 25 | 9 |
| $prop^5C$ | 282.1 → 80.0 | Qualifier | 2 | 3.9 | 0.4 | 100 | 41 | 9 |

Supplementary Table 8: Elution gradient and chromatographic parameters.

| Time [min] | Buffer B [%] |
|---|---|
| 0.0 | 0 |
| 1.0 | 0 |
| 7.0 | 60 |
| 7.2 | 100 |
| 8.8 | 100 |
| 9.0 | 0 |
| 12.0 | 0 |

Supplementary Table 9: Oligonucleotide sequences used in this study. (*IIIIII* = index sequence | N = randomized nucleotides | /5phos/ = phosphate | /rApp/ = 5′-adenylation | /3ddC/ = 3′-dideoxy-C)

| Name | Oligonucleotide sequence | Source |
|---|---|---|
| T7 template | GCACAGAGCAGCAAGAGGCAGTCCCGGGAGAGCGCCTATAGTGAGTCGTATTA | Biolegio |
| T7 primer | TAATACGACTCACTATAGG | Biolegio |
| RT-primer | GCACAGAGCAGCAAGAG | Biolegio |
| poly(dT)-Oligo | TTTTTTTTTT | Biolegio |
| L3-Adapter | /rApp/AGATCGGAAGAGCGGTTCAG/ddC/ | IDT |
| RT primer 2 | GGATCCTGAACCGCT | Biolegio |
| L##clip2.0 | /5Phos/NNNN*IIIIII*NNNNNAGATCGGAAGAGCGTCGTG/3ddC/ | IDT |
| P5Solexa_s | ACACGACGCTCTTCCGATCT | Biolegio |
| P3Solexa_s | CTGAACCGCTCTTCCGATCT | Biolegio |
| P5Solexa | AATGATACGGCGACCACCGAGATCTACACTCTTTCCCTACACGACGCTCTTCCGATCT | Biolegio |
| P3Solexa | CAAGCAGAAGACGGCATACGAGATCGGTCTCGGCATTCCTGCTGAACCGCTCTTCCGATCT | Biolegio |
| MALAT1_11:65.500.272_down | /5phos/CACATTTTTCAAACTAAGCTACTcagaggctgagtcgctgcat | Biolegio |
| MALAT1_11:65.500.272_up | tagccagtaccgtagtgcgtgAATTACTTCCGTTACGAAAGTCCT | Biolegio |
| MALAT1_1165.500.338_down | /5phos/CCAATGCAAAAACATTAAGTcagaggctgagtcgctgcat | Biolegio |
| MALAT1_11:65.500.338_up | tagccagtaccgtagtgcgtgGGATTTAAAAAATAATCTTAACTCAAAG | Biolegio |
| FLNB_3:58131631_down | /5phos/GTGAATCCAAGAACATATCATTGGCGAGcagaggctgagtcgctgcat | Biolegio |
| FLNB_3:58131631_up | tagccagtaccgtagtgcgtgGAGGGAAAACCCCATTTGCCACTTC | Biolegio |
| FLNB_3:58131742_down | /5phos/AATTTCCCCCAAACCATTTCTGCCCcagaggctgagtcgctgcat | Biolegio |
| FLNB_3:58131742_up | tagccagtaccgtagtgcgtgGAATCTGATCCCAGACCCATGACC | Biolegio |
| CTNNB1_3:41239837_down | /5phos/CCATTTGTATTGTTACTCCTCGcagaggctgagtcgctgcat | Biolegio |
| CTNNB1_3:41239837_up | tagccagtaccgtagtgcgtgTCTTCACTTCTTGAGTCACTCCCAAAA | Biolegio |
| CTNNB1_3:41239763_down | /5phos/CAGCTTGGTTAGTGTGTCAGGCACTcagaggctgagtcgctgcat | Biolegio |
| CTNNB1_3:41239763_up | tagccagtaccgtagtgcgtgAGTTTACTTCAATTGTTCCCATAGGAAAC | Biolegio |
| AHNAK_11:62518109_up | tagccagtaccgtagtgcgtgCCTTTCAGGTTCACATCCACACC | Biolegio |
| AHNAK_11:62518109_down | /5phos/GGCCCCTTCAGTTCGCCAGAcagaggctgagtcgctgcat | Biolegio |
| AHNAK_11:62518059_down | /5phos/CCACATTCGGTGCTGAAAcagaggctgagtcgctgcat | Biolegio |
| AHNAK_11:62518059_up | tagccagtaccgtagtgcgtgTTGGTCCTTCCAAGTTAAAG | Biolegio |
| WDR6_3:49012974_up | tagccagtaccgtagtgcgtgGGCAGCAGGTACCGACAACG | Biolegio |
| WDR6_3:49012974_down | /5phos/TCCTTGACAAAGATGGCCTTGCCcagaggctgagtcgctgcat | Biolegio |
| WDR6_3:49012969_up | tagccagtaccgtagtgcgtgCAGCAGGTACCGACAACGTTCCT | Biolegio |
| WDR6_3:49012969_down | /5phos/GACAAAGATGGCCTTGCCAGAGGcagaggctgagtcgctgcat | Biolegio |
| MARCH6_5:10433736_up | tagccagtaccgtagtgcgtgTGGGAAAAATCTCCAAAGAGAGAAG | Biolegio |
| MARCH6_5:10433736_down | /5phos/CCACAAAAAAGGACATGTAAAGGGcagaggctgagtcgctgcat | Biolegio |
| MARCH6_5:10433720_up | tagccagtaccgtagtgcgtgGAGAGAAGTCCACAAAAAAGGACA | Biolegio |
| MARCH6_5:10433720_down | /5phos/GTAAAGGGGAAGGTCAAGTTGTTGcagaggctgagtcgctgcat | Biolegio |
| NFX1_9:33348009_up | tagccagtaccgtagtgcgtgCTTTCCCCAGAGTCCCCAAAG | Biolegio |
| NFX1_9:33348009_down | /5phos/CCATTGTATCATTCTCATGCCTTTGCcagaggctgagtcgctgcat | Biolegio |
| NFX1_9:33348002_up | tagccagtaccgtagtgcgtgCCAGAGTCCCCAAAGTCCATTG | Biolegio |
| NFX1_9:33348002_down | /5phos/ATCATTCTCATGCCTTTGCGTCCcagaggctgagtcgctgcat | Biolegio |
| SRRM2_16:2768757_up | tagccagtaccgtagtgcgtgGGATCAAGGTTAAACCTCAAGAATG | Biolegio |

| | | |
|---|---|---|
| SRRM2_16:2768757_down | /5phos/CCCTGATGGAGGAATAAGGCcagaggctgagtcgctgcat | Biolegio |
| SRRM2_16:2768735_up | tagccagtaccgtagtgcgtgCCCTGATGGAGGAATAAGGCC | Biolegio |
| SRRM2_16:2768735_down | 5phos/CTGGGCTCAGGTGTGGAAGcagaggctgagtcgctgcat | Biolegio |
| MAT2A_2:85544902_down | 5phos/GTACTTACGCCATACCCCcagaggctgagtcgctgcat | Biolegio |
| MAT2A_2:85544902_up | tagccagtaccgtagtgcgtgCTGAGGTGATGGCTTCTC | Biolegio |
| MAT2A_2:85544538_up | tagccagtaccgtagtgcgtgACTAGGAGTTGGTGAGGTTTGGC | Biolegio |
| MAT2A_2:85544538_down | 5phos/AGTGCTGAAACCATGCATAGGATTGcagaggctgagtcgctgcat | Biolegio |
| SELECT PCR fwd | ATGCAGCGACTCAGCCTCTG | Biolegio |
| SELECT PCR rev | TAGCCAGTACCGTAGTGCGTG | Biolegio |

Supplementary Table 10: Reagents, chemicals and special solvents used in synthesis.

| Chemicals | CAS number | Vendor | Cat.No. |
|---|---|---|---|
| 1*H*-Imidazole | 288-32-4 | AppliChem | A1073,1000 |
| 4-Methylbenzene-1-sulfonyl chloride | 98-59-9 | Acros Organics | 139031000 |
| 5-Methyluridine [m5U] | 1463-10-1 | Sigma-Aldrich | 535893-25G |
| Acetic acid (LC-MS grade) | 64-19-7 | Merck | 5.33001.0050 |
| Ammonia solution (7N in methanol) | 7664-41-7 | Acros Organics | 428381000 |
| Ammonium acetate (LC-MS grade) | 631-61-8 | VWR | 84885.180 |
| Ammonium chloride | 12125-02-9 | Acros Organics | 123340010 |
| Ammonium fluoride | 12125-01-8 | Sigma-Aldrich | 216011-100G |
| Azobisisobutyronitrile | 78-67-1 | Sigma-Aldrich | 441090-25G |
| Copper(I) cyanide | 544-92-3 | Merck | 8.41811.0100 |
| Ethynylmagnesium bromide (0.5M in THF) | 4301-14-8 | Sigma-Aldrich | 346152-100ML |
| *N*-Bromosuccinimide | 128-08-5 | Alfa Aesar | A15922 |
| *N*-Methylpiperidine | 626-67-5 | Acros Organics | 127480100 |
| *tert*-Butyldimethylsilyl chloride | 18162-48-6 | Sigma-Aldrich | 190500-100G |
| Triethylamine | 121-44-8 | Carl Roth | X875.4 |

| Special solvents | CAS number | Vendor | Cat.No. |
|---|---|---|---|
| Acetonitrile (HPLC grade) | 75-05-8 | Honeywell | 34851-2.5L |
| Acetonitrile (LC-MS grade) | 75-05-8 | Fisher Scientific | A/0638/17 |
| Dichloromethane (dried over molecular sieve) [DCM] | 75-09-2 | Acros Organics | 348465000 |
| Dimethylformamide (dried over mol. sieve) [DMF] | 68-12-2 | Acros Organics | 348431000 |
| Tetrahydrofuran (dried over mol. sieve) [THF] | 109-99-9 | Acros Organics | 348451000 |

Supplementary Table 11: Commercially available compounds used as standards for LC-QqQ-MS quantification

| Standards for LC-QqQ-MS quantification | CAS number | Vendor | Cat.No. |
|---|---|---|---|
| 2′-(*O*-Propargyl)-adenosine [$A_{prop}$] | 151390-97-5 | Jena Bioscience | CLK-RP-3401-10 |
| 2′-*O*-Methyladenosine [$A_m$] | 2140-79-6 | Cayman Chemical | 16936 |
| 5-Methyl-cytidine [m5C] | 2140-61-6 | Jena Bioscience | N-RP-1832-1G |
| Adenosine [A] | 58-61-7 | Sigma-Aldrich | A9251-25G |
| Cytidine [C] | 65-46-3 | Sigma-Aldrich | C122106-1G |
| $N^6$-Methyladenosine [$m^6A$] | 1867-73-8 | Carbosynth | NM32281 |
| $N^6$-Propargyl-adenosine [$prop^6A$] | 67005-97-4 | Jena Bioscience | CLK-N004-5 |

Supplementary Figures:

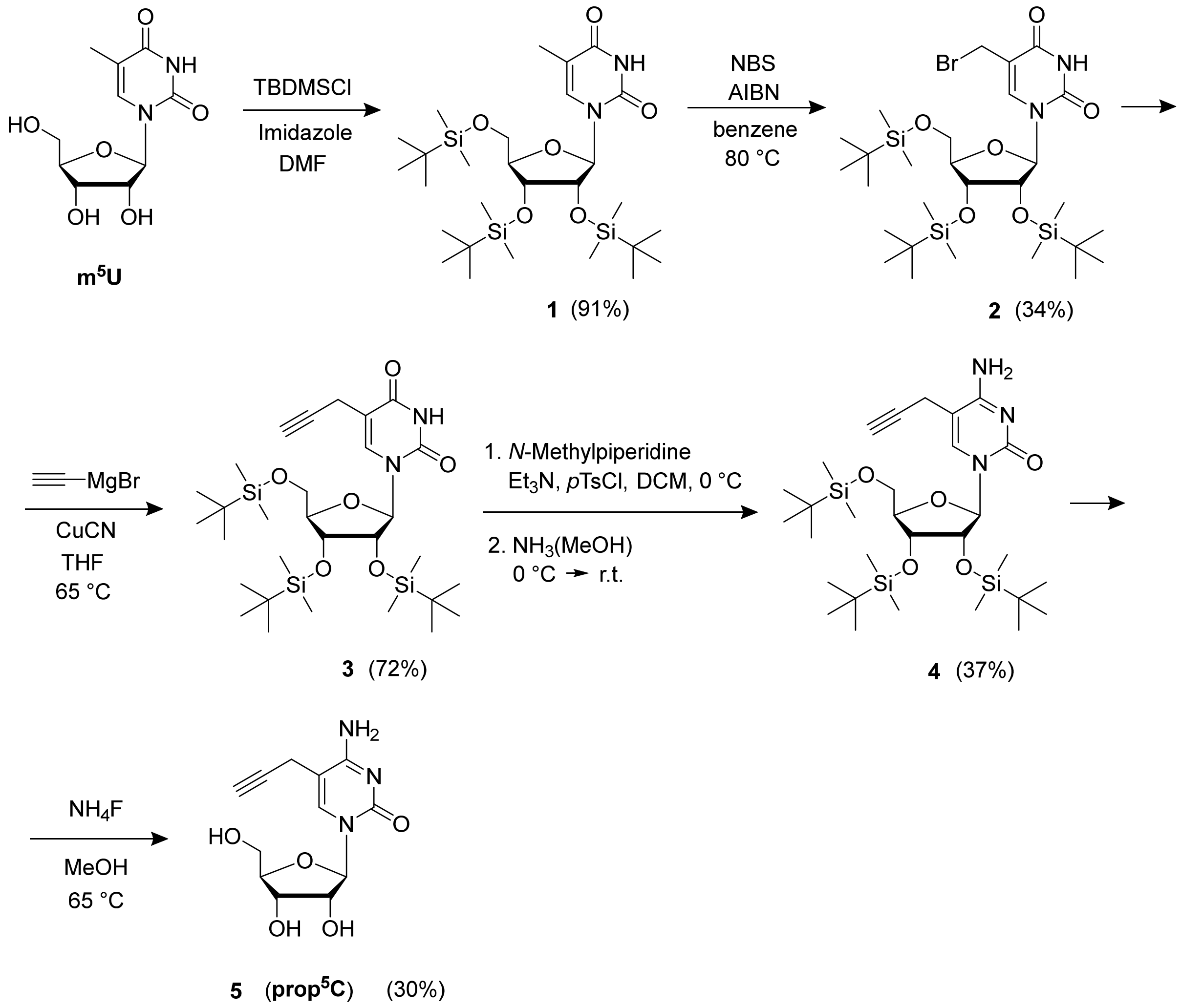

Supplementary Figure 1. Synthesis of prop$^5$C.

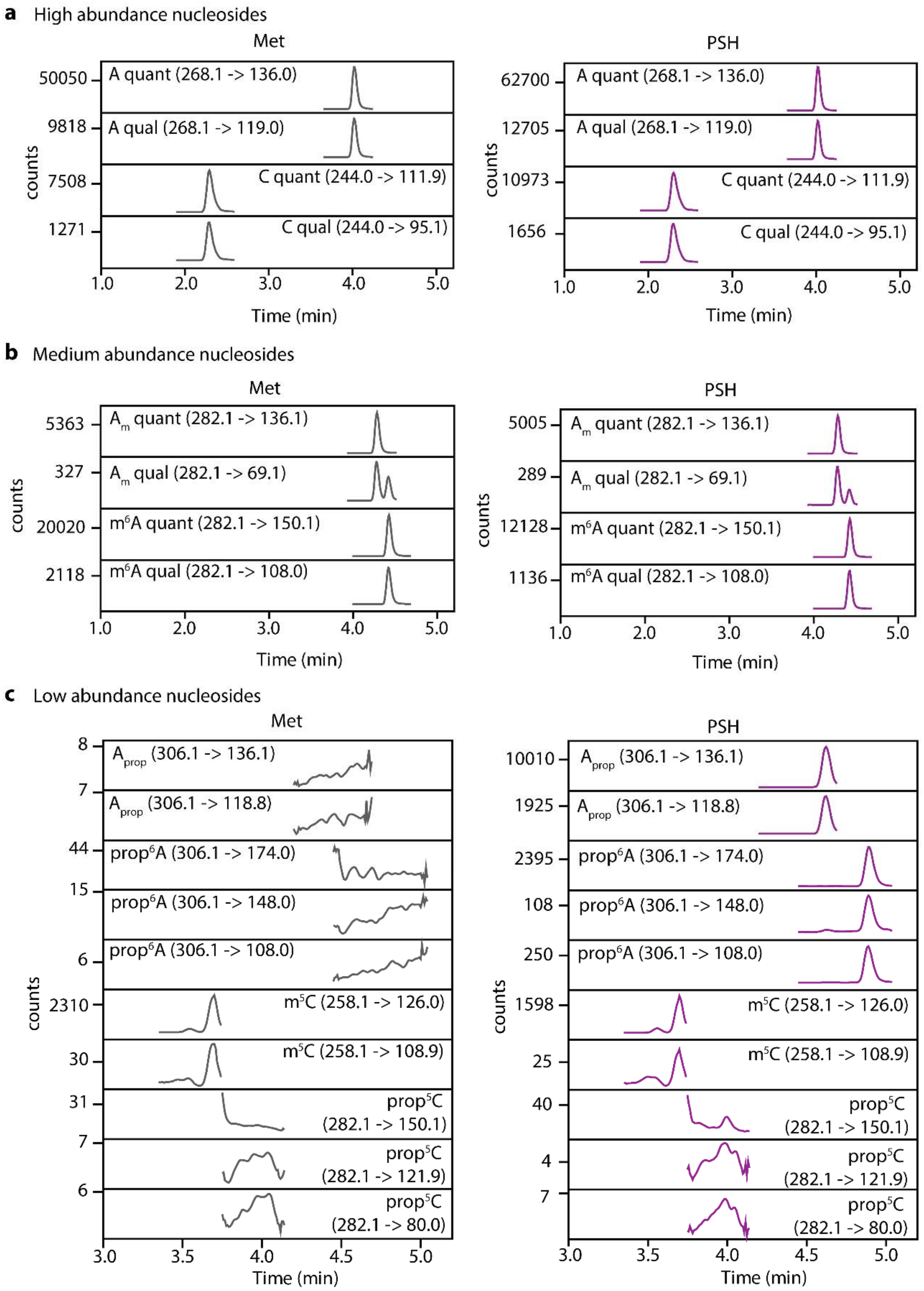

Supplementary Figure 2. Detection and quantification of modified nucleosides in mRNA. a-c, Examples runs of LC-QqQ-MS analysis with dynamic MRM to detect and quantify high abundant (a), medium abundant (b), or low abundant (c) modified nucleosides in digested, dephosphorylated, poly(A)$^+$ RNA from HeLa cells metabolically labeled with 2.5 mM PSH (purple) or Met (gray) as control. Shown are the extracted ion counts vs time for the retention time windows of the different analytes as well as the indicated fragmentation.

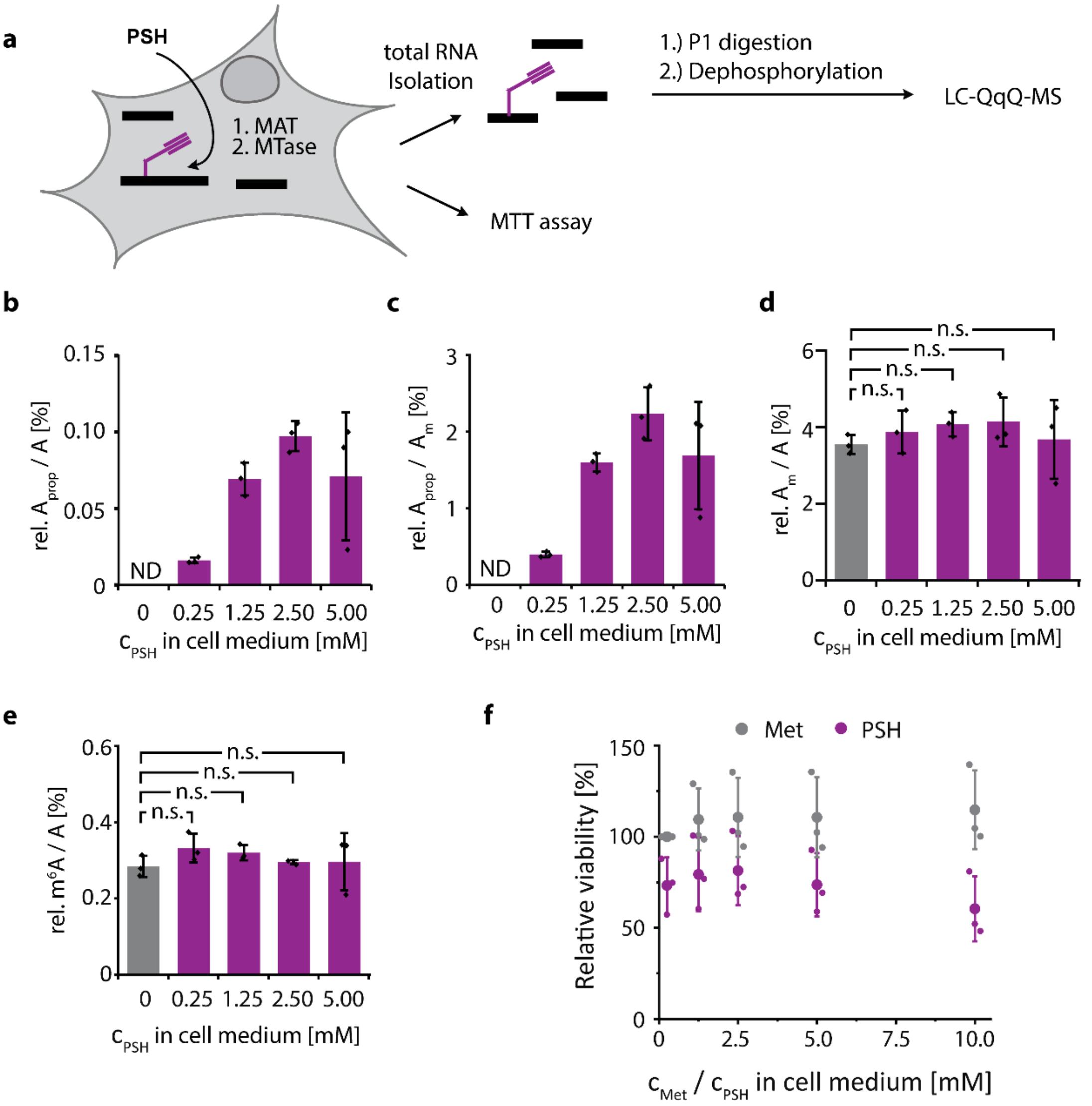

Supplementary Figure 3. Optimization of metabolic labeling of HeLa cells with PSH. a, Scheme. b-c, Quantification of modified nucleotides in total RNA from HeLa cells treated with indicated concentrations of PSH. (b) $A_{prop}$ relative to A, (c) $A_{prop}$ relative to $A_m$, (d) $A_m$ relative to A and (e) $m^6A$ relative to A. Quantification from dynamic MRM run on LC-QqQ-MS using external synthetic standards. Not detected (ND) means no signal with correct quantifier detected.. f, Cell viability from MTT assay for HeLa cells treated with different amounts of PSH or methionine in cell media for 16 h. Vitality of HeLa cells after metabolic labeling relative to cells in standard conditions (0.25 mM Met), determined by MTT assay. Mean values ± SD from n=3 biological replicates are shown. Statistical significance determined via independent two-tailed t-test (n.s. P>0.05; * P≤0.05; ** P≤0.01; *** P≤0.001).

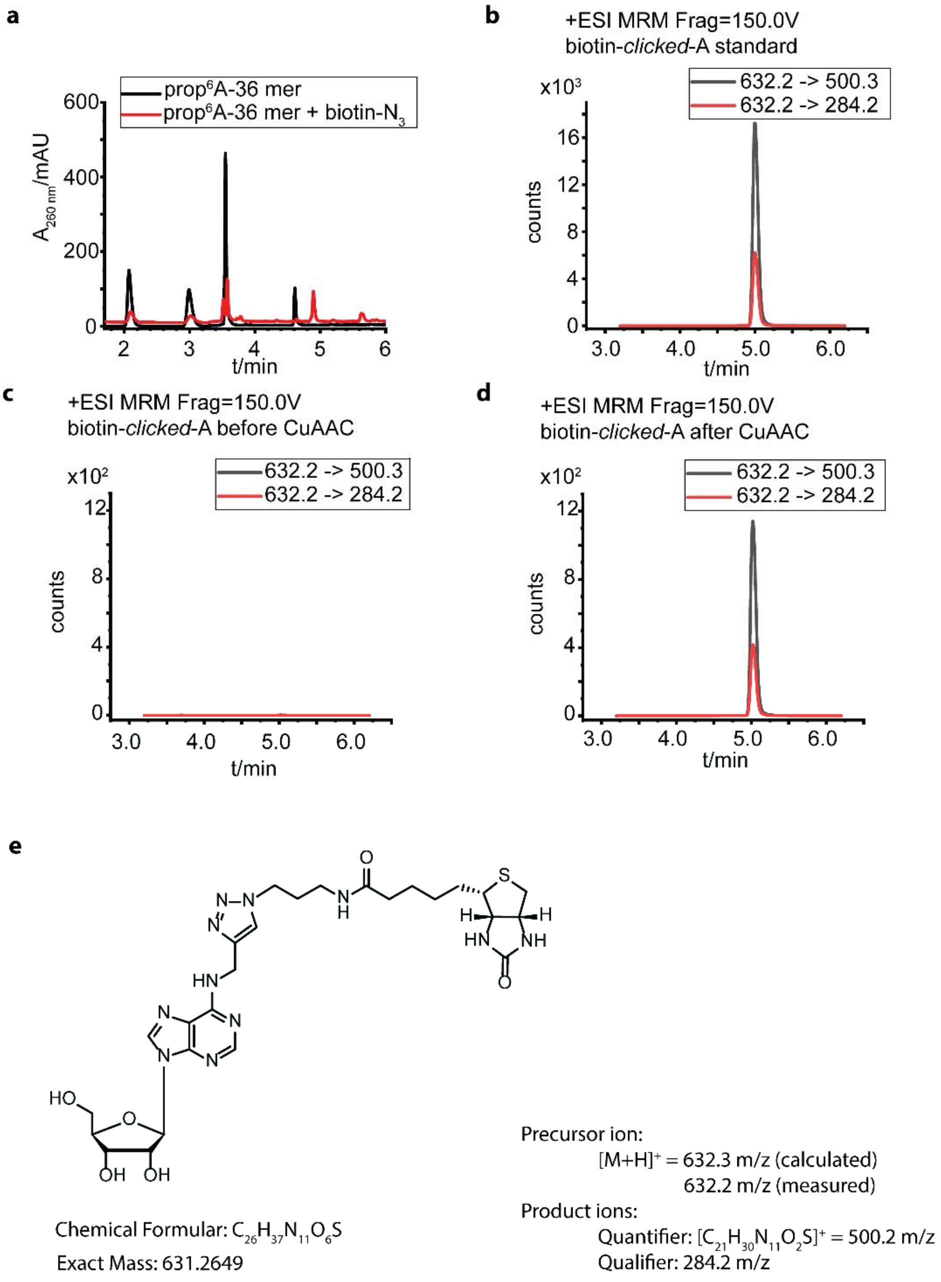

Supplementary Figure 4. Efficiency of CuAAC reaction on RNA containing N6-propargyl-adenosine. a, HPLC run of prop$^6$A-containing RNA (36mer) before and after CuAAC with biotin azide. b-d, LC-QqQ-MS analysis of expected click-product shown in (e). m/z=632.2 correspond to biotin clicked A (e), m/z=500.3 was the most abundant fragment corresponding to the ribose loss and m/z=284.2 is the second most abundant fragment observed from biotin-clicked A standard. Fragmentation of $N^6$-biotinylated adenosine (b) was used to verify product formation and observed after CuAAC (d) but not in a control without biotin azide (c). e, Structure of biotin product after CuAAC.

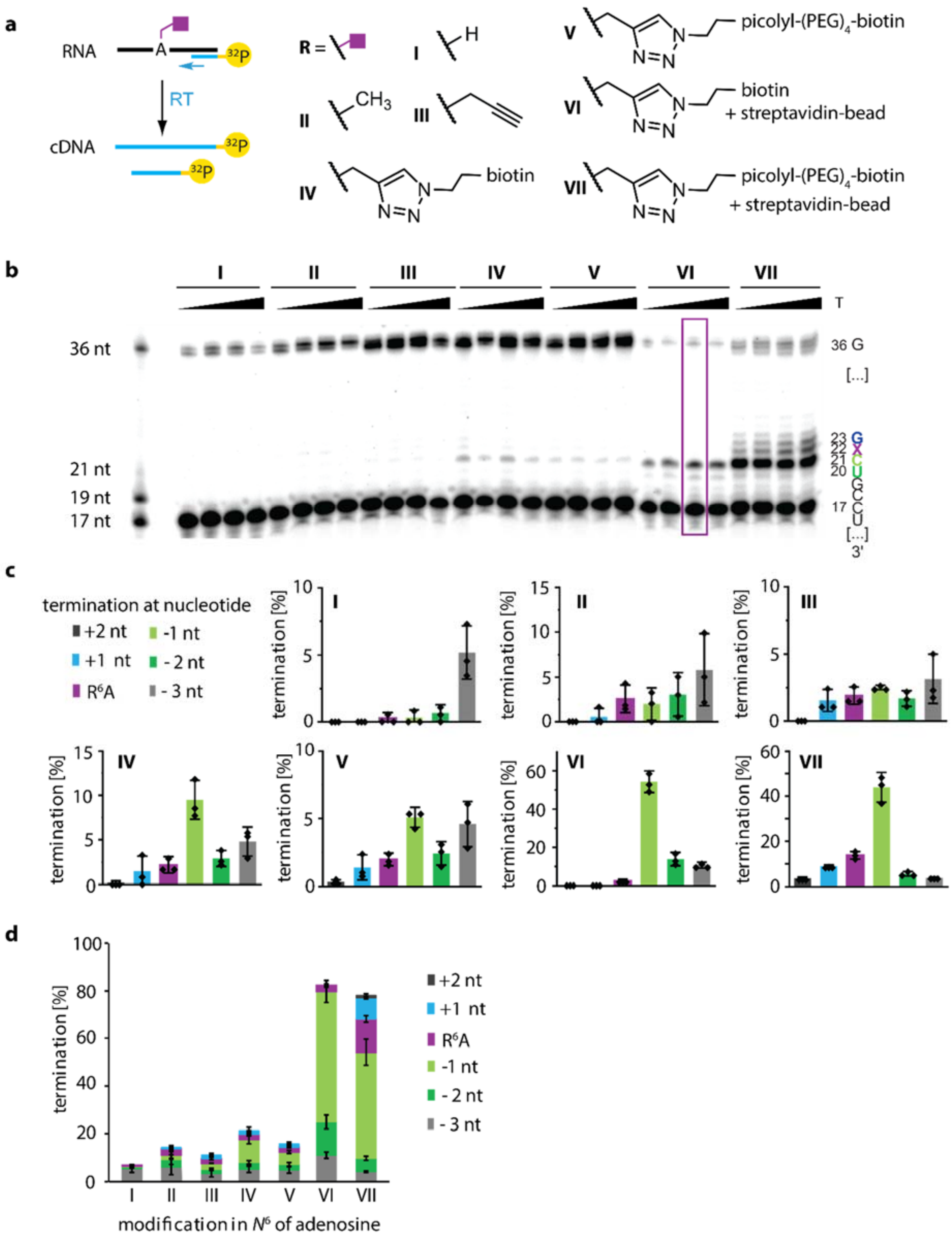

Supplementary Figure 5: Optimizing termination of reverse transcription (RT) with SuperScript SSIV at $N^6$-modified adenosines.a, Schematic illustration of reverse transcription to test the effect of indicated $N^6$-modifications (purple square). b, Denat. PAGE (15 % PAA, 1× TBE) showing primer extension assay of modified RNA with SS IV at different temperatures (25 °C, 37 °C, 45 °C, 50 °C and 57 °C). Shown is one representative gel of n=3 biological replicates. Purple box indicates conditions chosen for library preparation. c, Quantification of primer extension experiments. Termination was quantified for indicated positions from PAGE analyses and normalized to the total cDNA yield. d, Stacked data of individual positions shown in (c) illustrate overall termination. Mean values ± SD of n=3 indpendent experiments are shown.

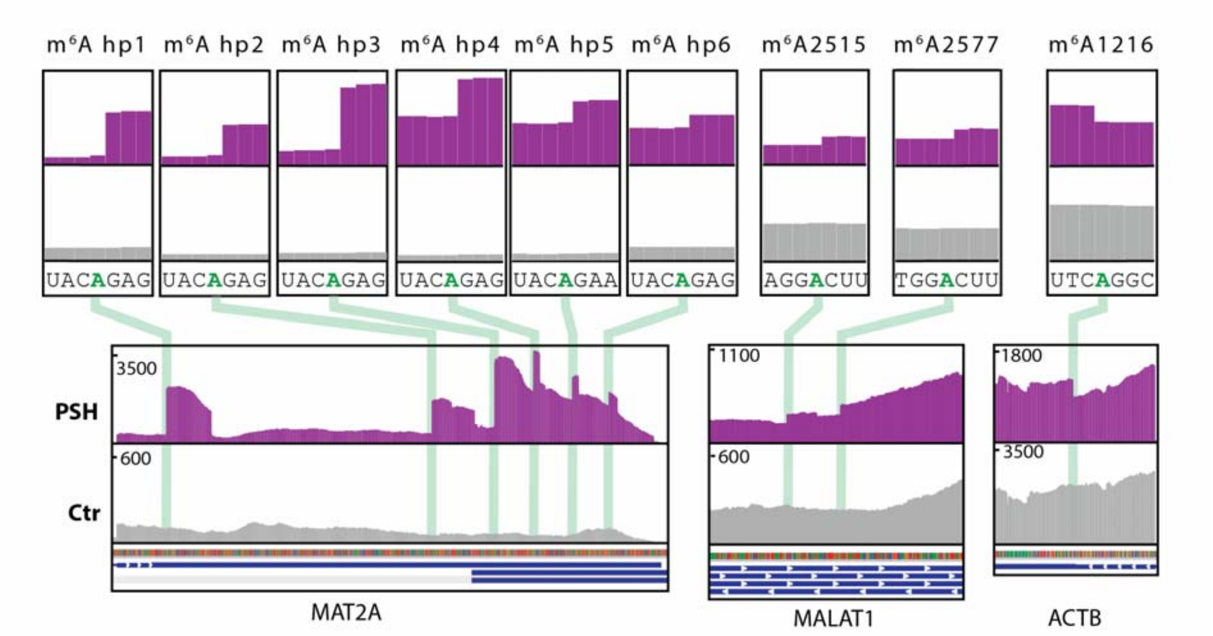

Supplementary Figure 6: Integrative genomics viewer (IGV) browser coverage tracks of MePMe-seq data for MAT2A, MALAT1 and ACTB RNAs. Cells labeled with PSH (purple) or methionine as control (gray). One representative example of n=2 biological replicates is shown.

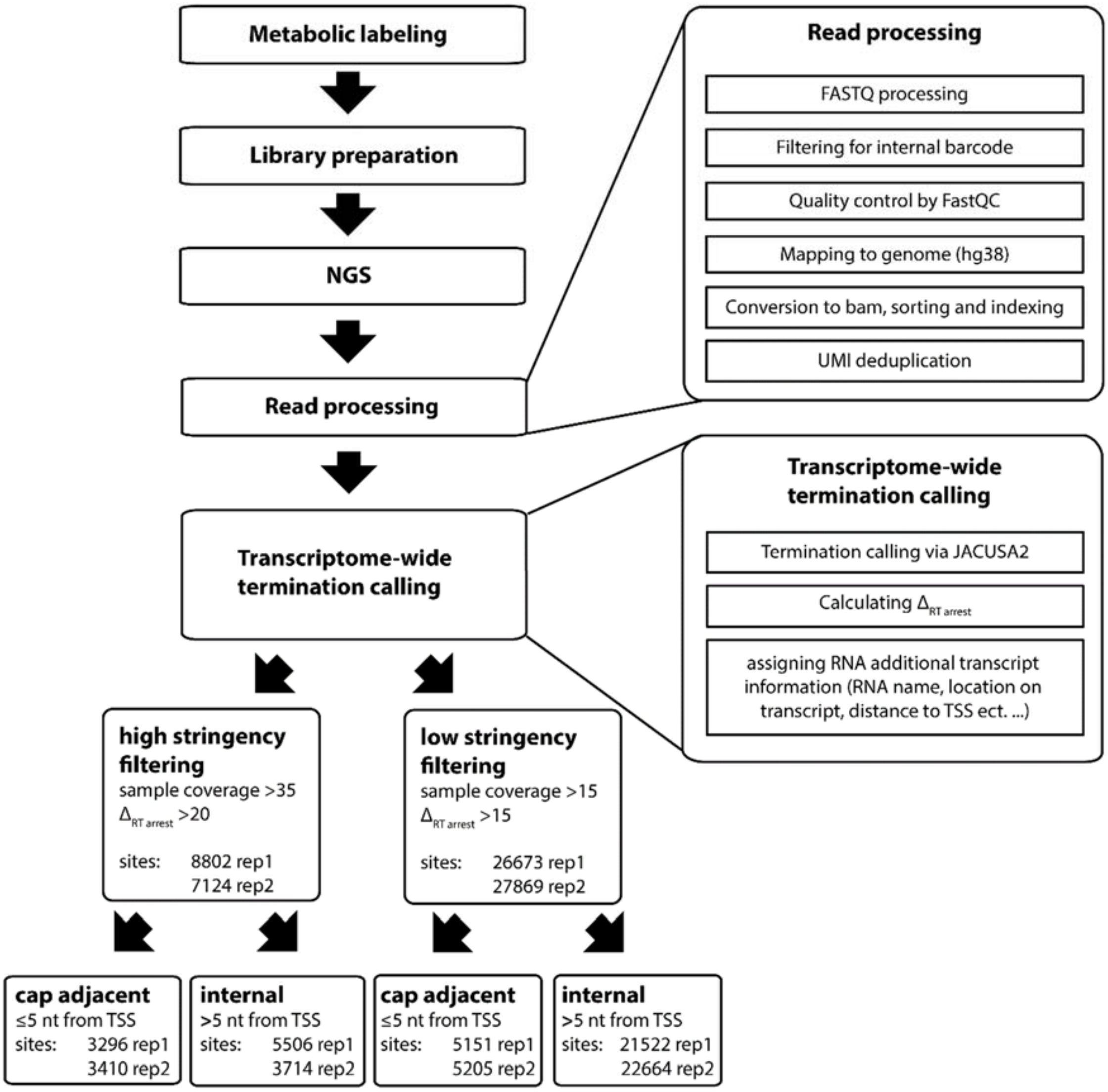

Supplementary Figure 7: Flow chart illustrating steps of bioinformatic analysis performed for MePMe-seq, indicating filter settings applied to the data sets and the number of sites identified when the indicated filter settings are applied.

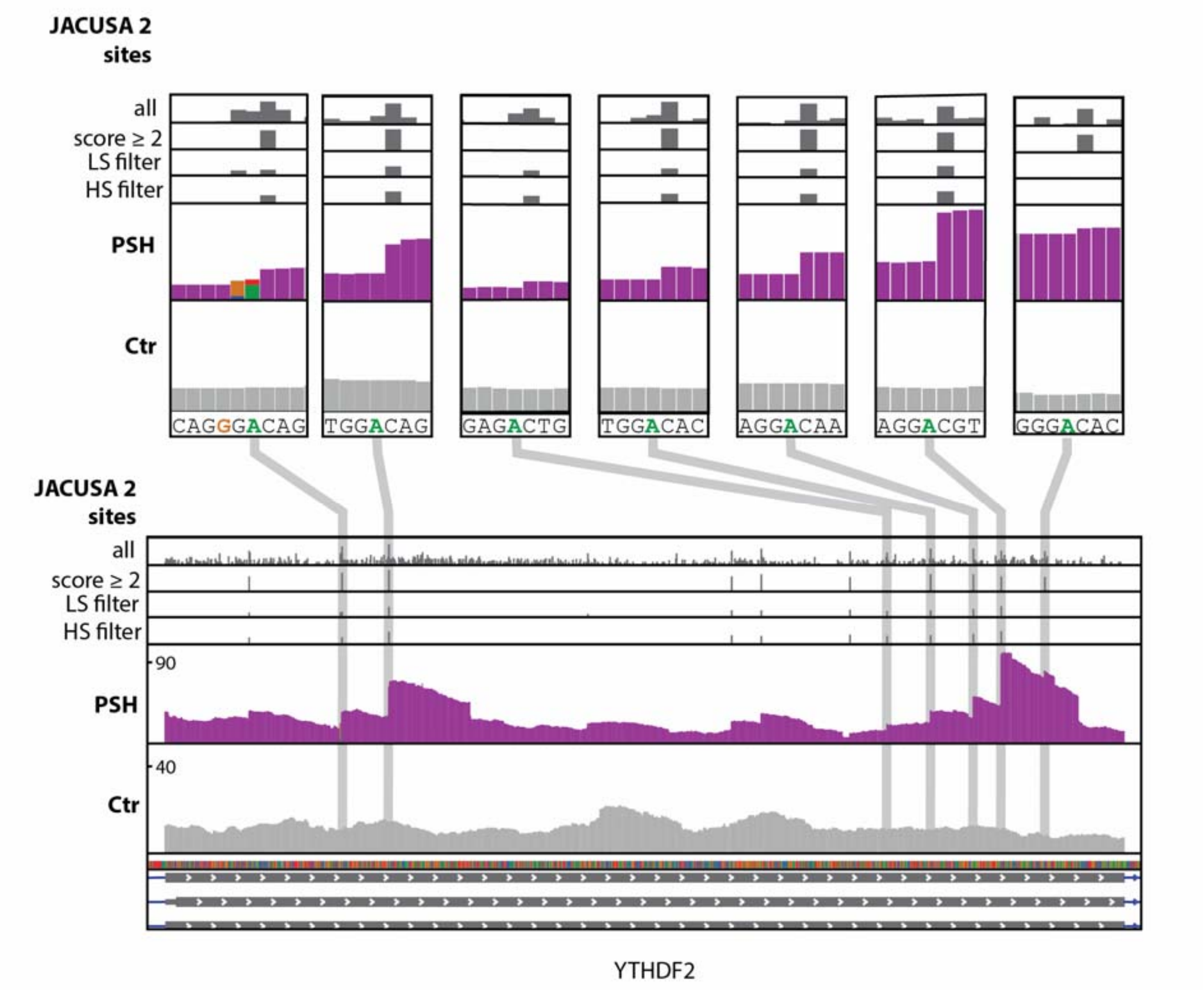

Supplementary Figure 8: IGV browser coverage tracks of MePMe-seq data for YTHDF2 mRNA. Cells labeled with PSH (purple) or methionine as control (gray). Gray bars represent positions called by JACUSA2 as terminations under the indicated filtering conditions. One representative example of n=2 biological replicates is shown.

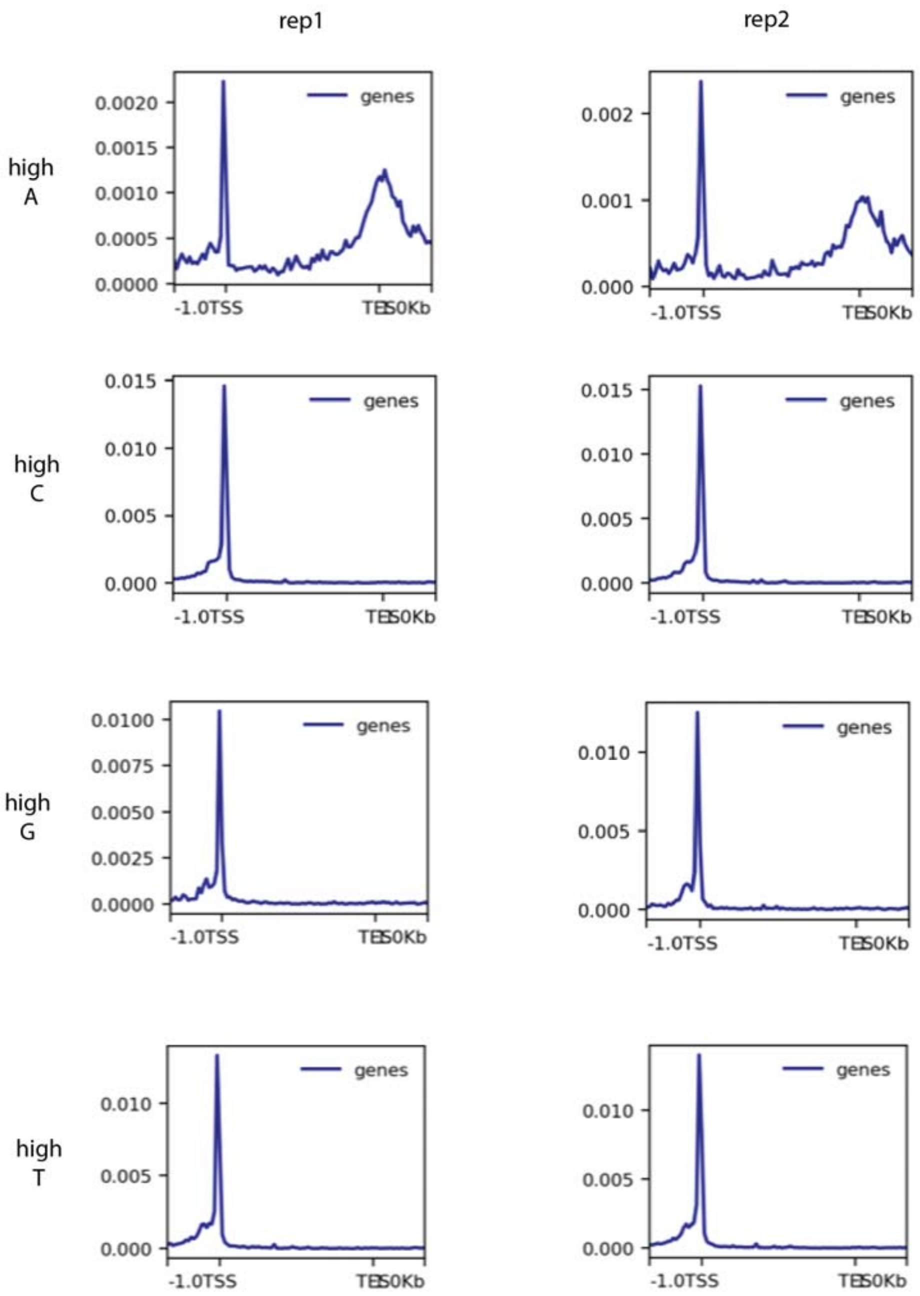

Supplementary Figure 9: Clustering of JACUSA2 hits around the TSS. Frequency of JACUSA2 hits plotted against their position on the transcript showing clustering ~5 nt around the TSS for terminations at all nucleoside identities in both replicates.

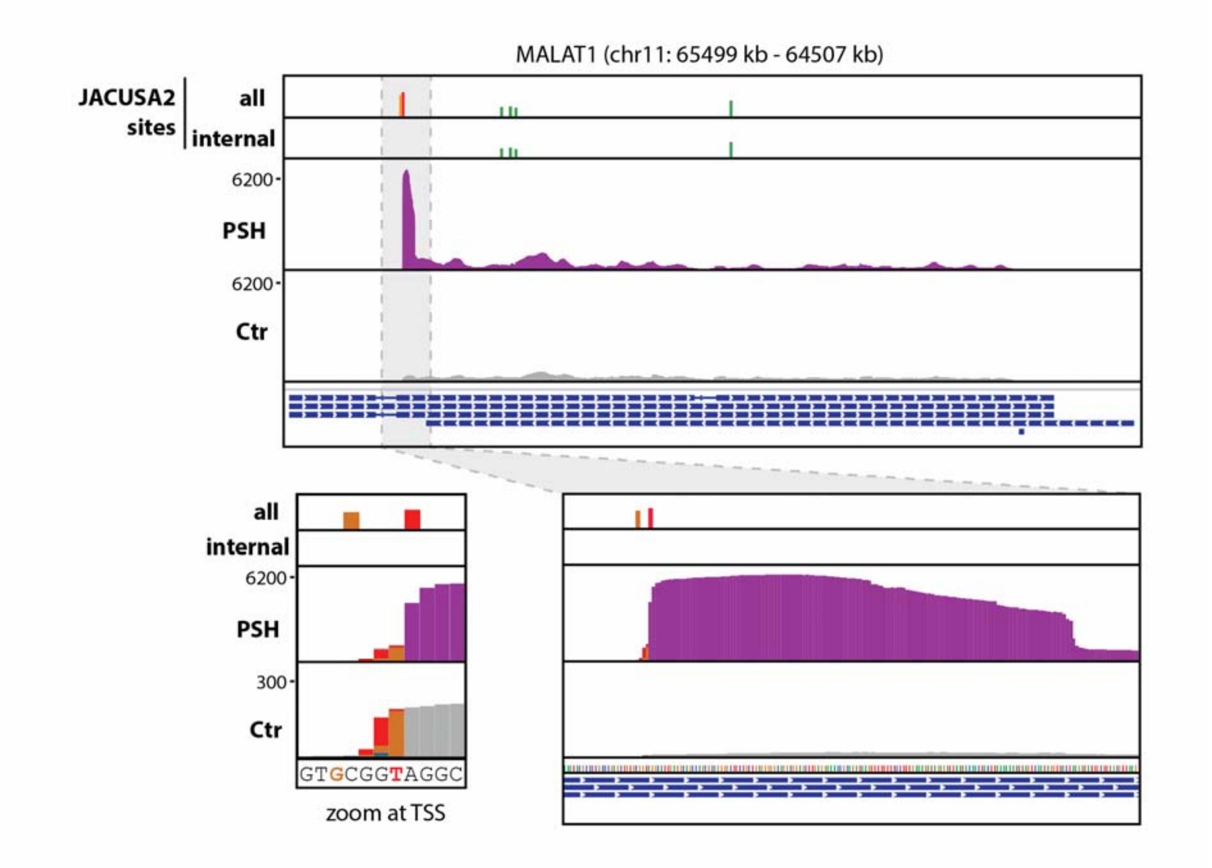

Supplementary Figure 10: Enrichment and terminations at 5′ end of transcripts. IGV browser coverage tracks of MePMe-seq data for MALAT1 RNA from cells labeled with PSH (purple) or methionine as control (gray). Colored bars represent terminations (red=T, orange=G, green=A,) identified by JACUSA2 hits (HS filtered).

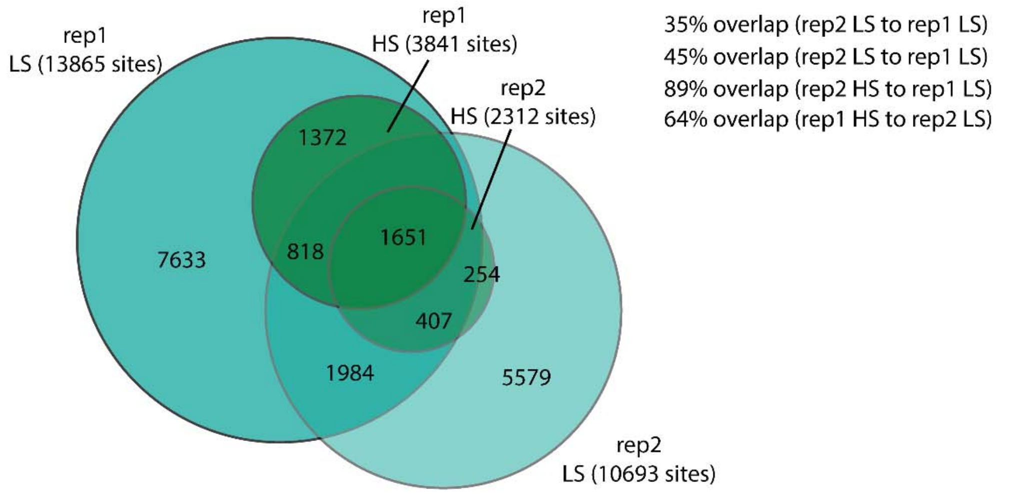

Supplementary Figure 11: Overlap of $m^6A$ sites identified in n=2 biological replicates of MePMe-seq experiments, when HS or LS filter settings are applied (% calculated as indicated).

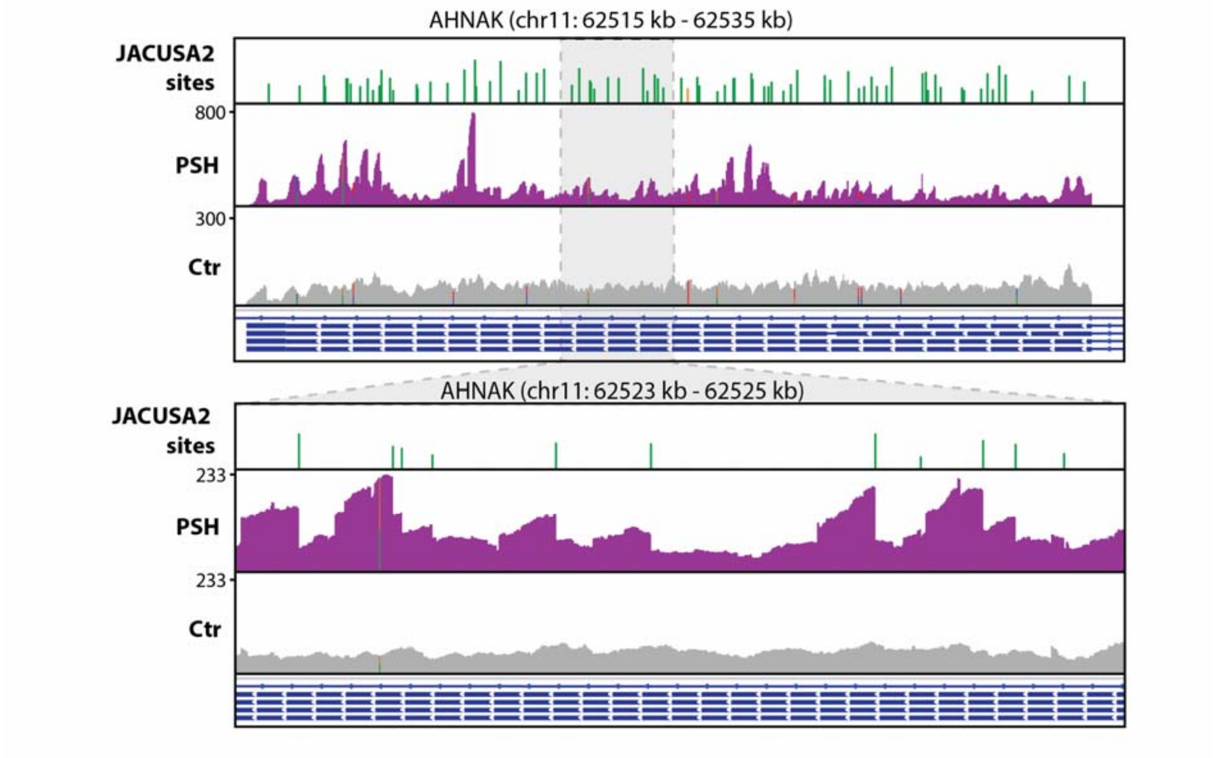

Supplementary Figure 12 Clustering of m$^6$A sites identified via MePMe-seq. IGV browser coverage tracks of MePMe-seq data for AHNAK mRNA from cells labeled with PSH (purple) or methionine as control (gray). Colored bars represent terminations (orange=G, green=A,) identified by JACUSA2 (HS filtered). One representative example of n=2 biological replicates is shown.

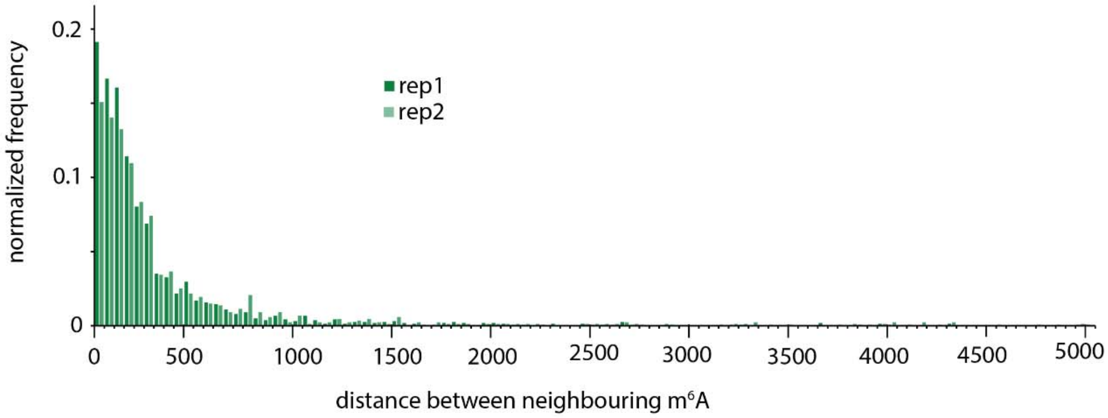

Supplementary Figure 13: Frequency of distance between neighboring m$^6$A sites. Full dataset of data shown in Figure 3e with a cutoff at distances >5000 nt.

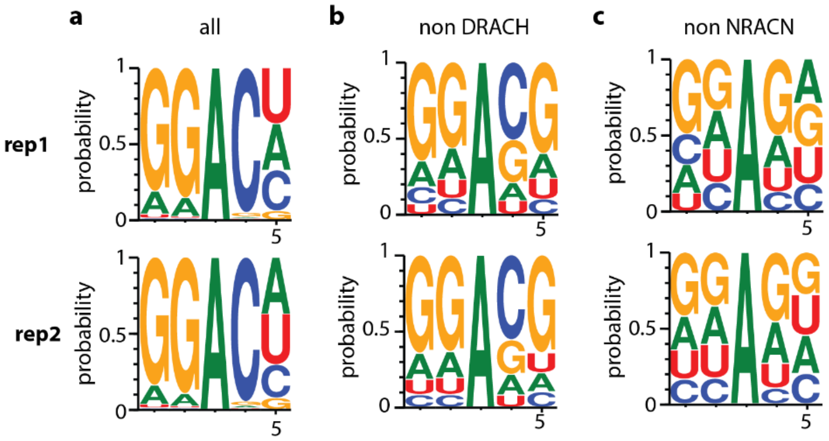

Supplementary Figure 14: Consensus motif for sequences surrounding identified m$^6$A (HS filtering) All 5 mers are considered as shown in Figure 3g for 1 representative dataset. (a), if DRACH sequences are excluded (b) and if NRACN sequences are excluded (c). Here, data for n=2 biologically replicates is shown.

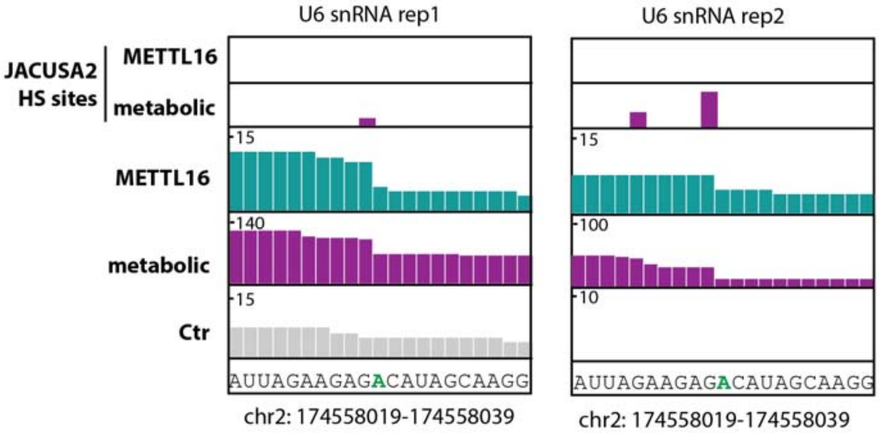

Supplementary Figure 15: IGV browser coverage tracks of U6 snRNA. Data for RNA labeled via METTL16 (turquoise), metabolic labeling (purple) or control (gray) are shown for n=2 biological replicates.

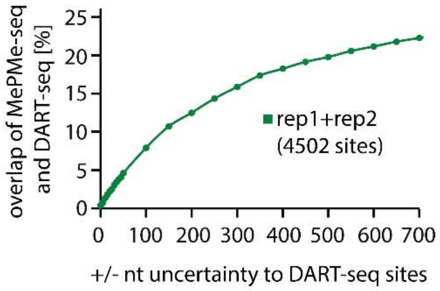

Supplementary Figure 16. Overlap of $m^6A$ sites identified by MePMe-seq and DART-seq depending on uncertainty allowed.

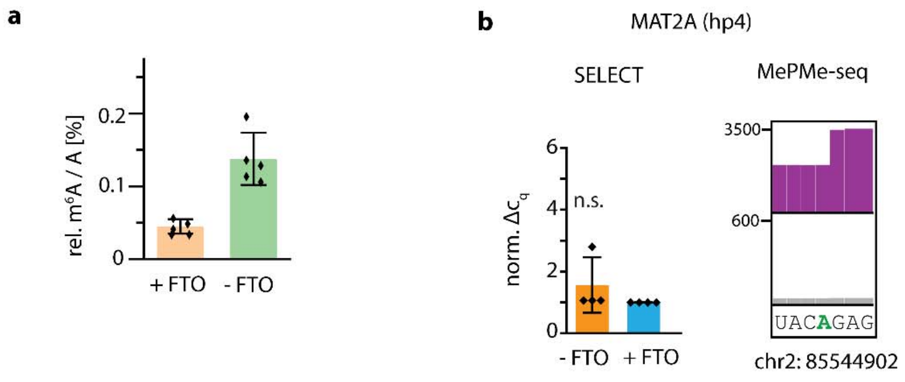

Supplementary Figure 17. SELECT experiments. a, LC-QqQ analysis of poly(A)$^+$ RNA with and without FTO treatment to verify demethylation. Mean values and SD for n=5 biological replicates are shown. b, The normalized $\Delta C_q$ values of SELECT qPCR measurements for reported m$^6$A in hp4 of MAT2A and the respective IGV browser coverage tracks of MePMe-seq data for the same site from cells grown with PSH (purple) and control (gray). Mean values and SD for n=4 biological replicates are shown. Statistical significance determined via one-sample one-tailed t-test (n.s. P>0.05; * P≤0.05; ** P≤0.01; *** P≤0.001).

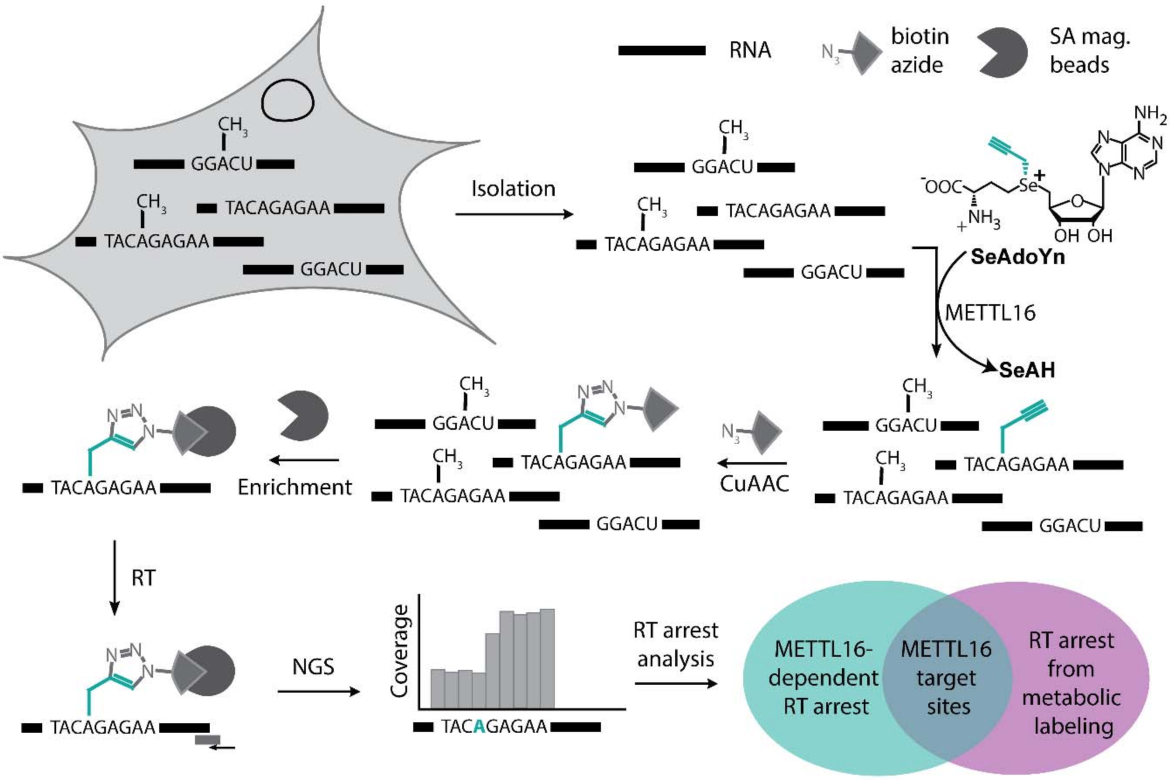

Supplementary Figure 18: Scheme illustrating *in vitro* METTL16-dependent labeling combined with metabolic labeling to identify METTL16 target sites.

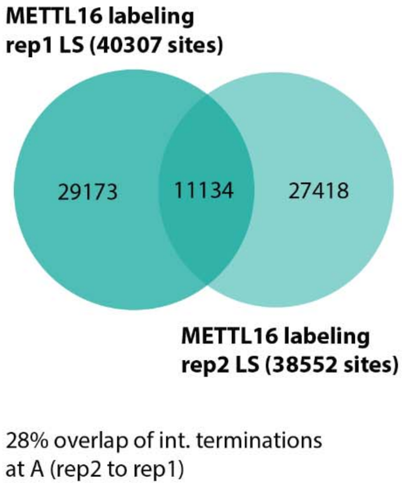

Supplementary Figure 19: Overlap of sites identified in n=2 biological replicates of *in vitro* METTL16 labeling experiments using low stringency filtering.

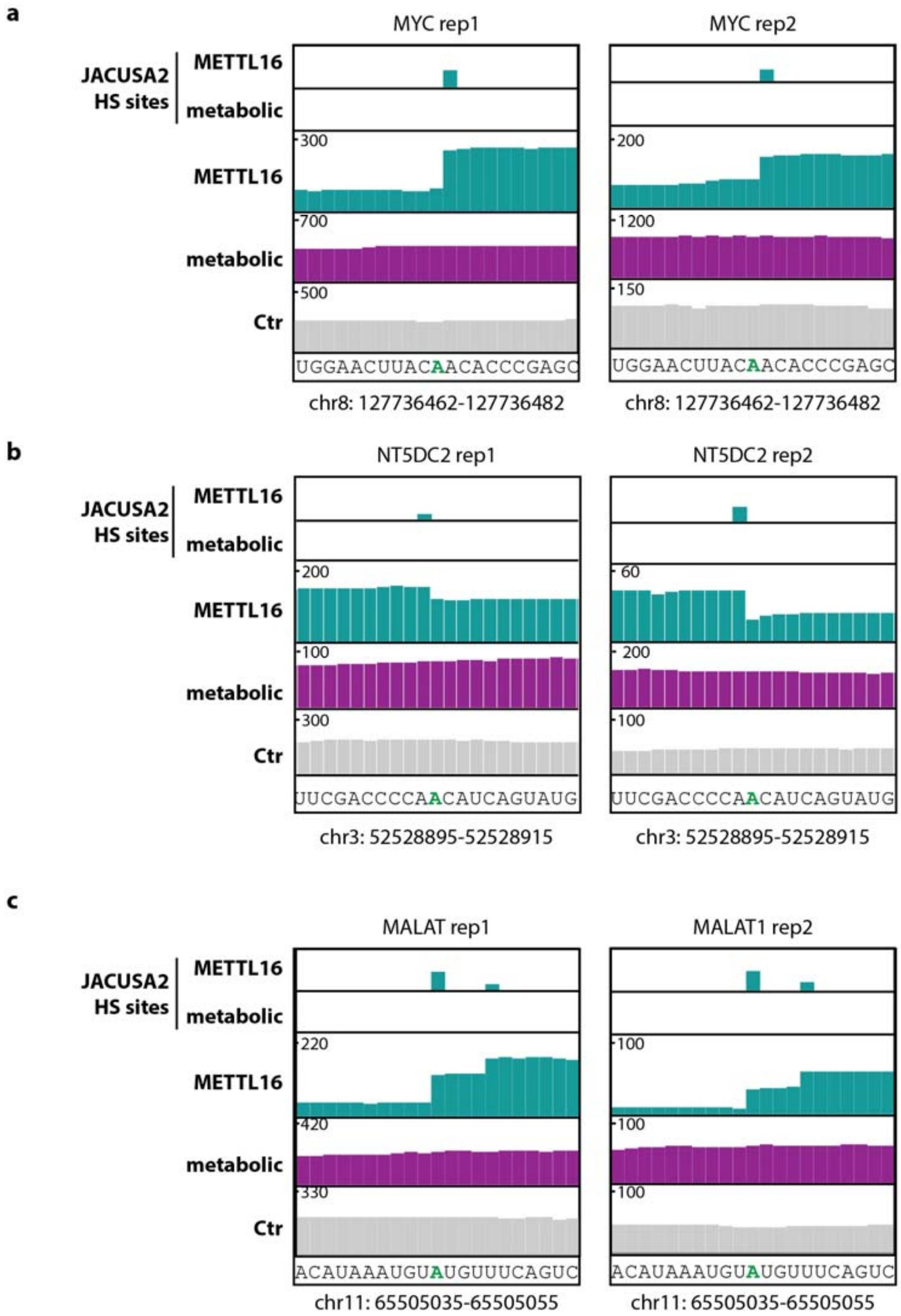

Supplementary Figure 20: IGV browser coverage tracks of *in vitro* METTL16 (cyan) and metabolic (purple) labeling data for indicated RNAs,(a) MYC, (b) NT5DC2 and (c) MALAT1 RNA. Control (Ctr) experiment is shown in gray. Data for n=2 biological replicates are shown.

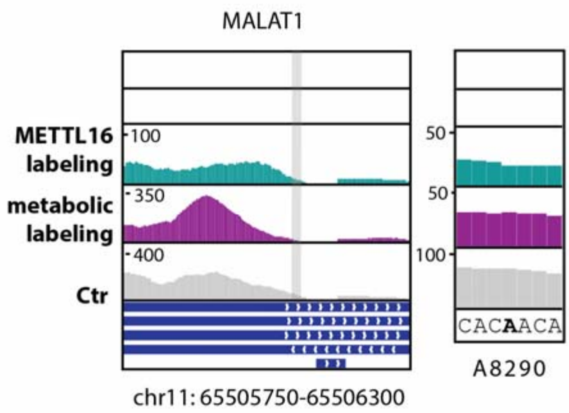

Supplementary Figure 21: IGV browser coverage tracks of *in vitro* METTL16 (cyan) and metabolic (purple) labeling data for featuring putative m$^6$A site at A8290 in MALAT1 RNA. Control (Ctr) experiment is shown in gray. One representative example of n=2 biological replicates is shown.

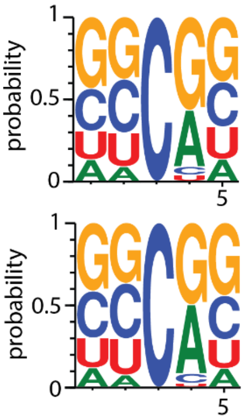

Supplementary Figure 22: Consensus motifs for modified C shown in Figure 6a for n=2 biological replicates. Consensus motif for 5mer sequences around identified terminations next to C when filtering with HS filtering. Results

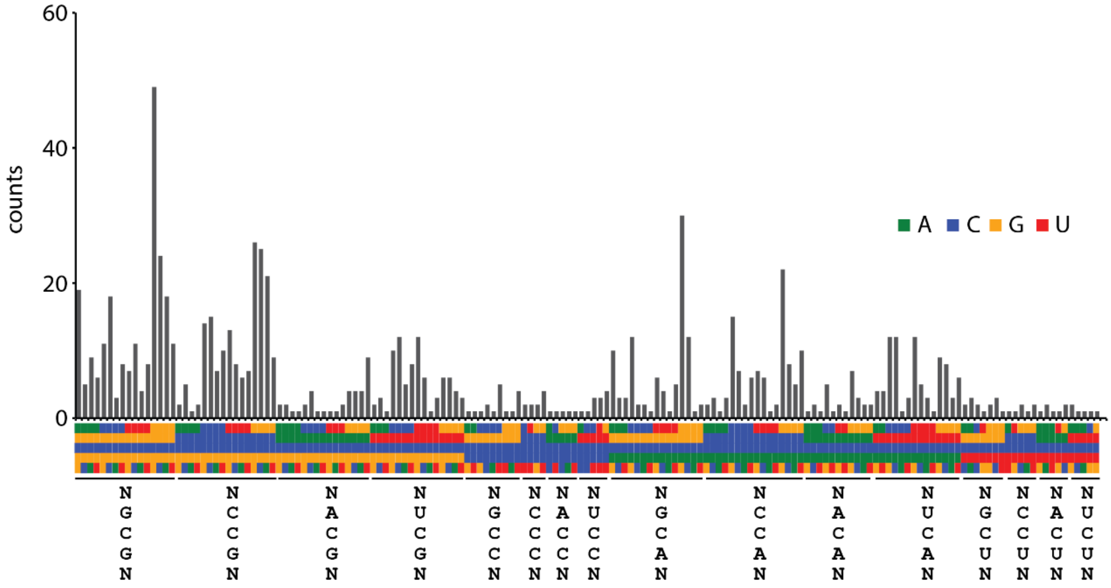

Supplementary Figure 23: Counts per sequence motif around identified m$^5$C sites (HS filtering), sorted by consensus motif.

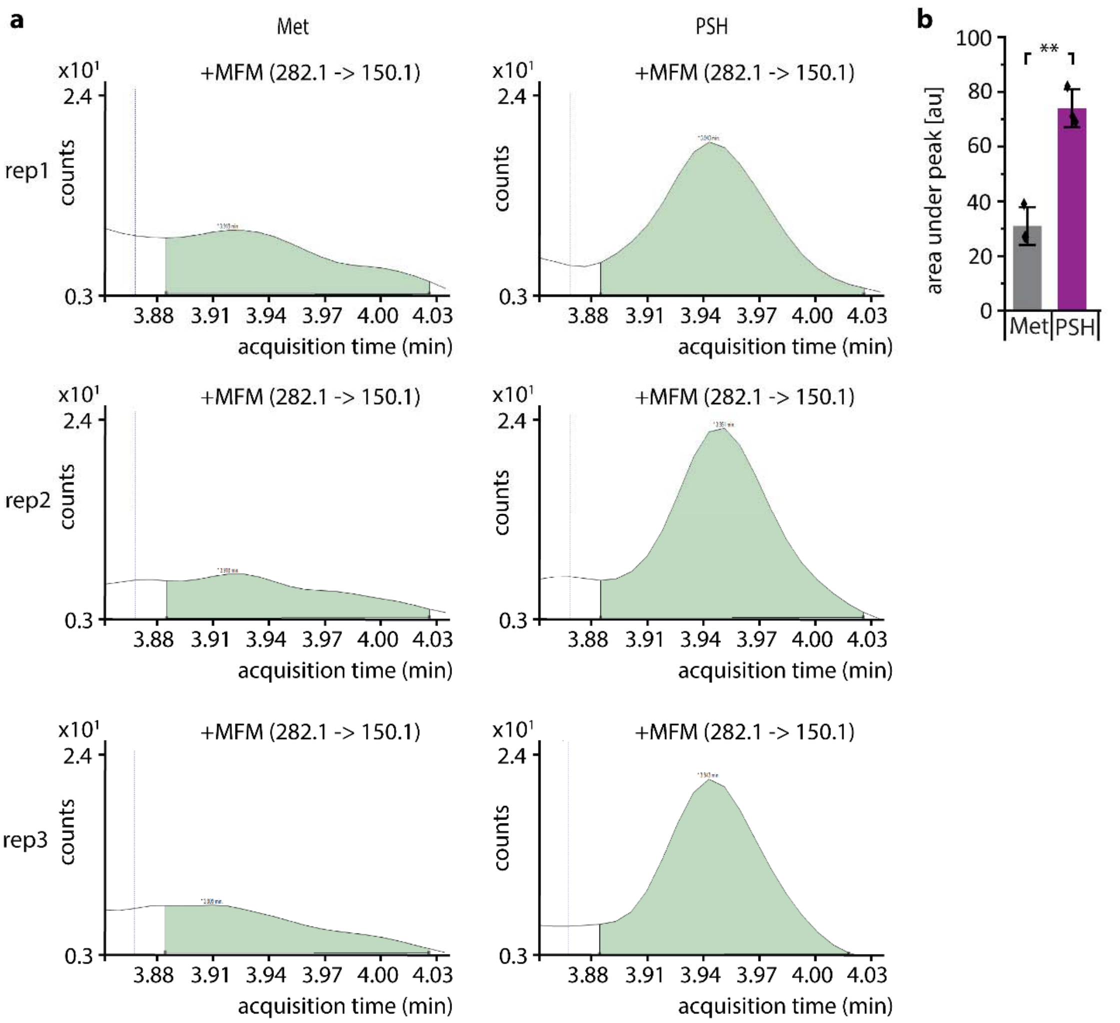

Supplementary Figure 24: LC-QqQ analysis of m5C in poly(A)$^+$-enriched RNA from HeLa cells after metabolic labeling with methionine (Met) or propargyl-selenohomocysteine (PSH), respectivel.a, Signal m$^5$C quantifier detected by dynamic MRM on LC-QqQ-MS from digested, dephosphorylated, poly(A)$^+$ enriched RNA from HeLa cells metabolically labeled with 2.5 mM PSH or Met as control. Area was chosen with exact same criteria for baseline and retention time frame. b, Area under peak for m$^5$C quantifier. Means values ± SD for n=3 biological replicates are shown. Statistical significance determined via independent two-tailed t-test (n.s. P>0.05; * P≤0.05; ** P≤0.01; *** P≤0.001).

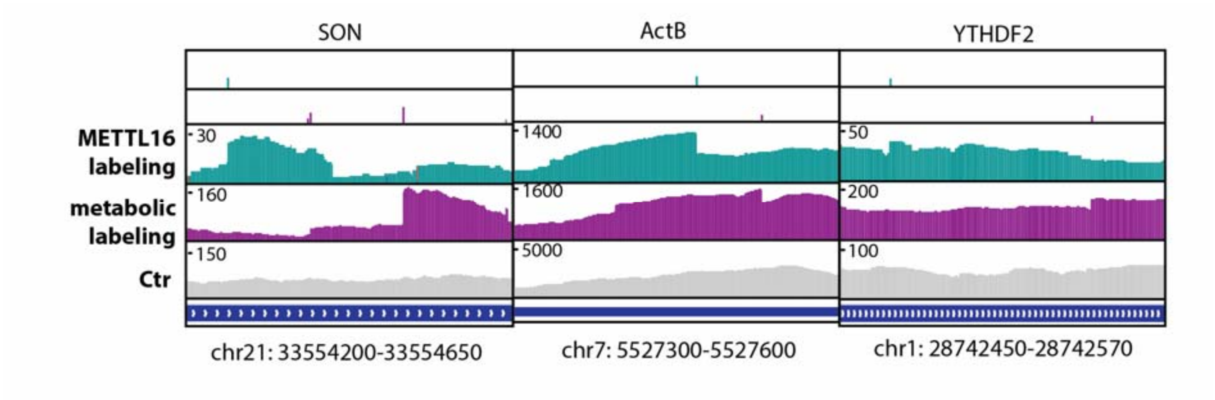

Supplementary Figure 25: IGV browser coverage tracks of *in vitro* METTL16 (cyan) and metabolic (purple) labeling data for indicated RNAs. Examples of mRNAs sites detected in the same transcript and region, but different positions. Control (Ctr) experiment is shown in gray. One representative example of n=2 biological replicates is shown.

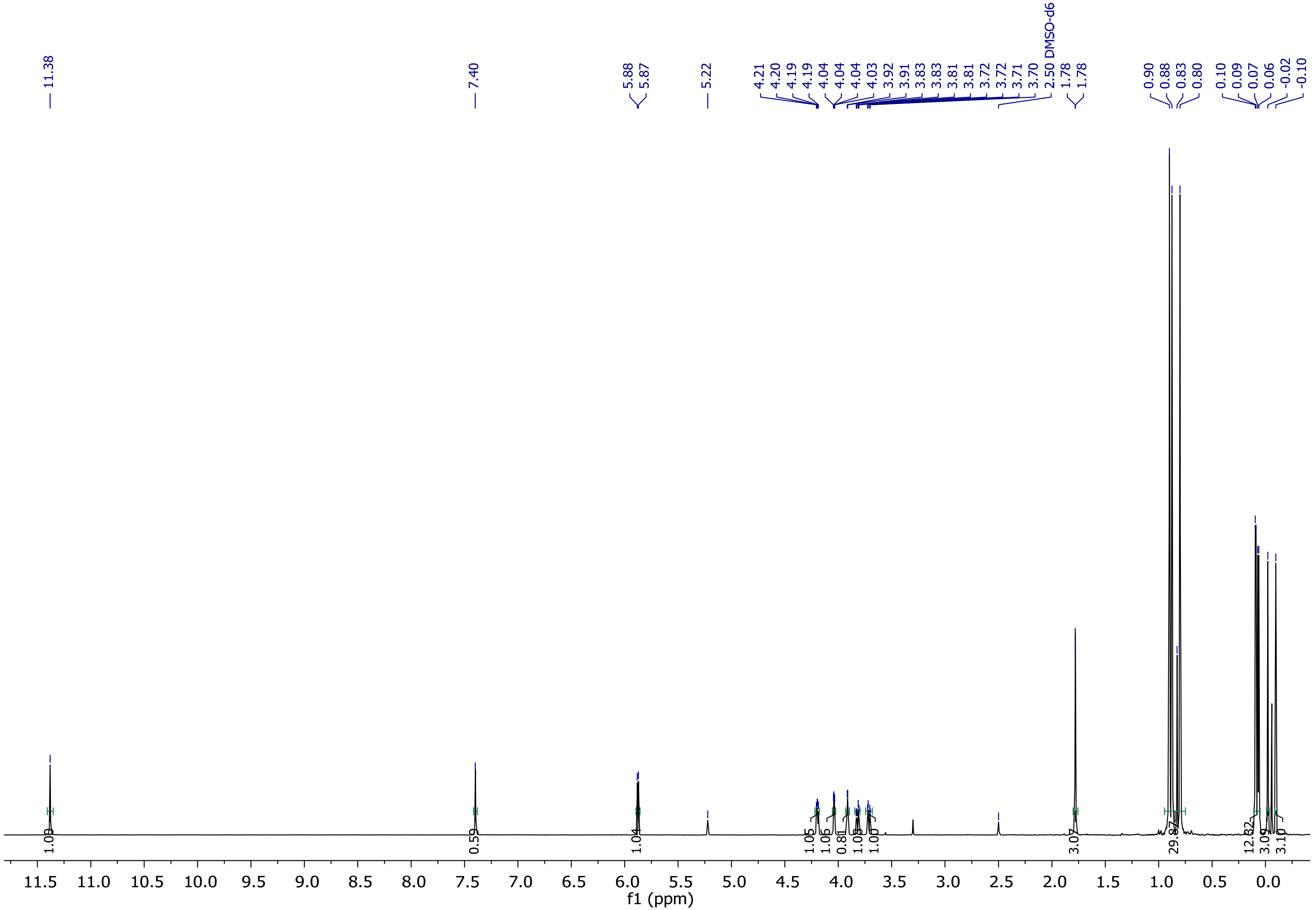

Supplementary Figure 26: [1]H-NMR spectrum of compound 1.

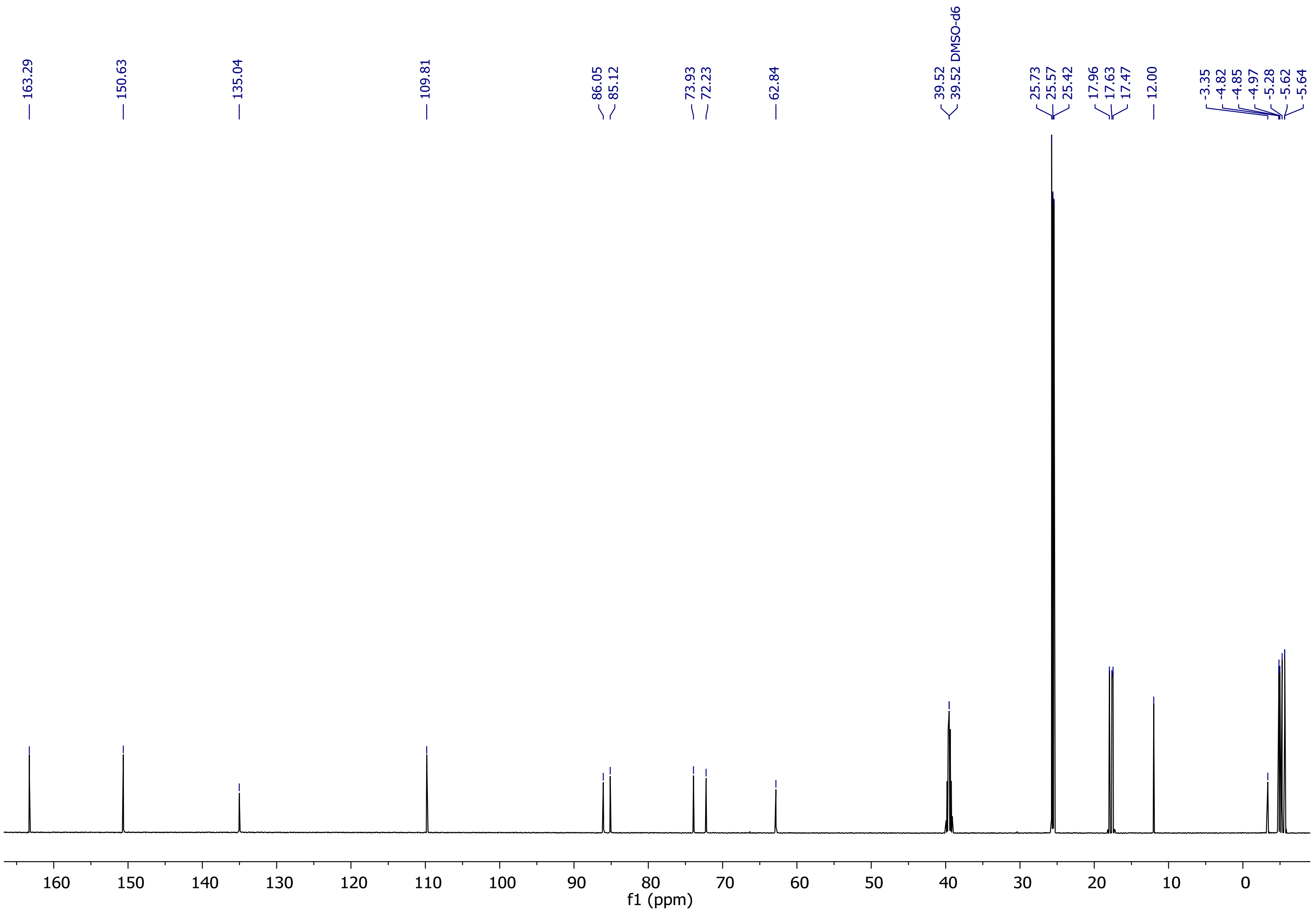

Supplementary Figure 27: $^{13}C$-NMR spectrum of compound 1.

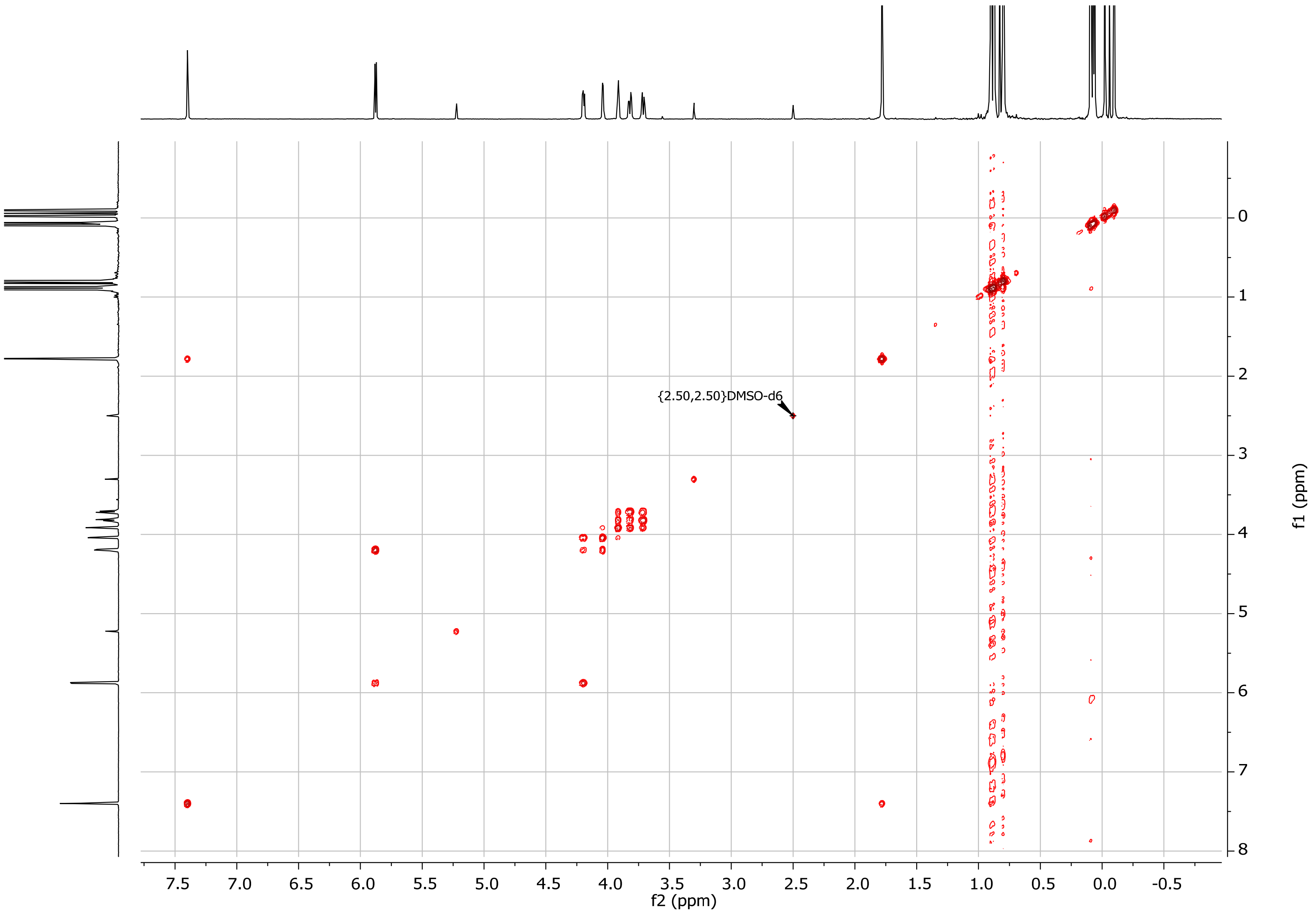

Supplementary Figure 28:  COSY spectrum of compound 1 .

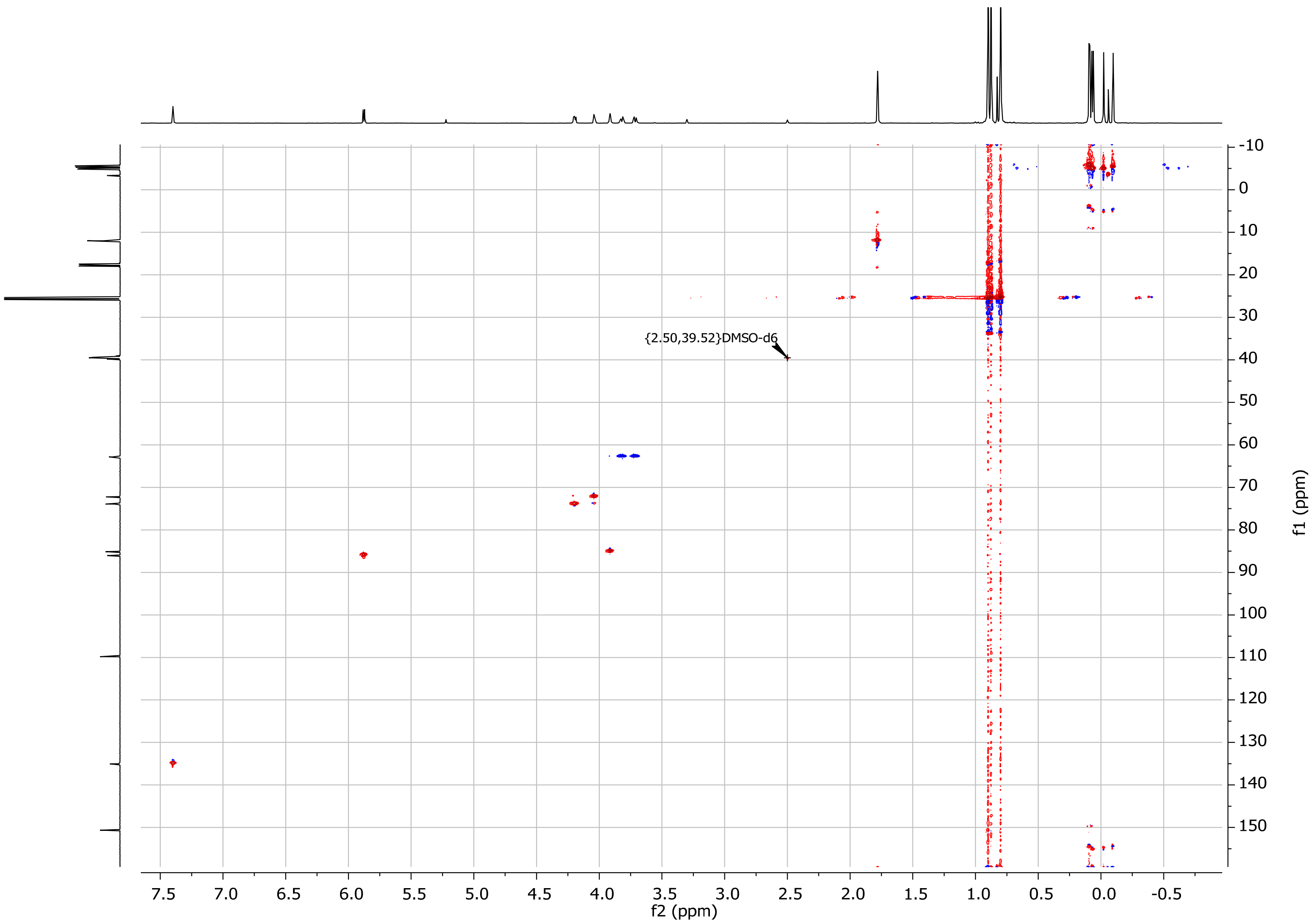

Supplementary Figure 29: HSQC spectrum of compound 1.

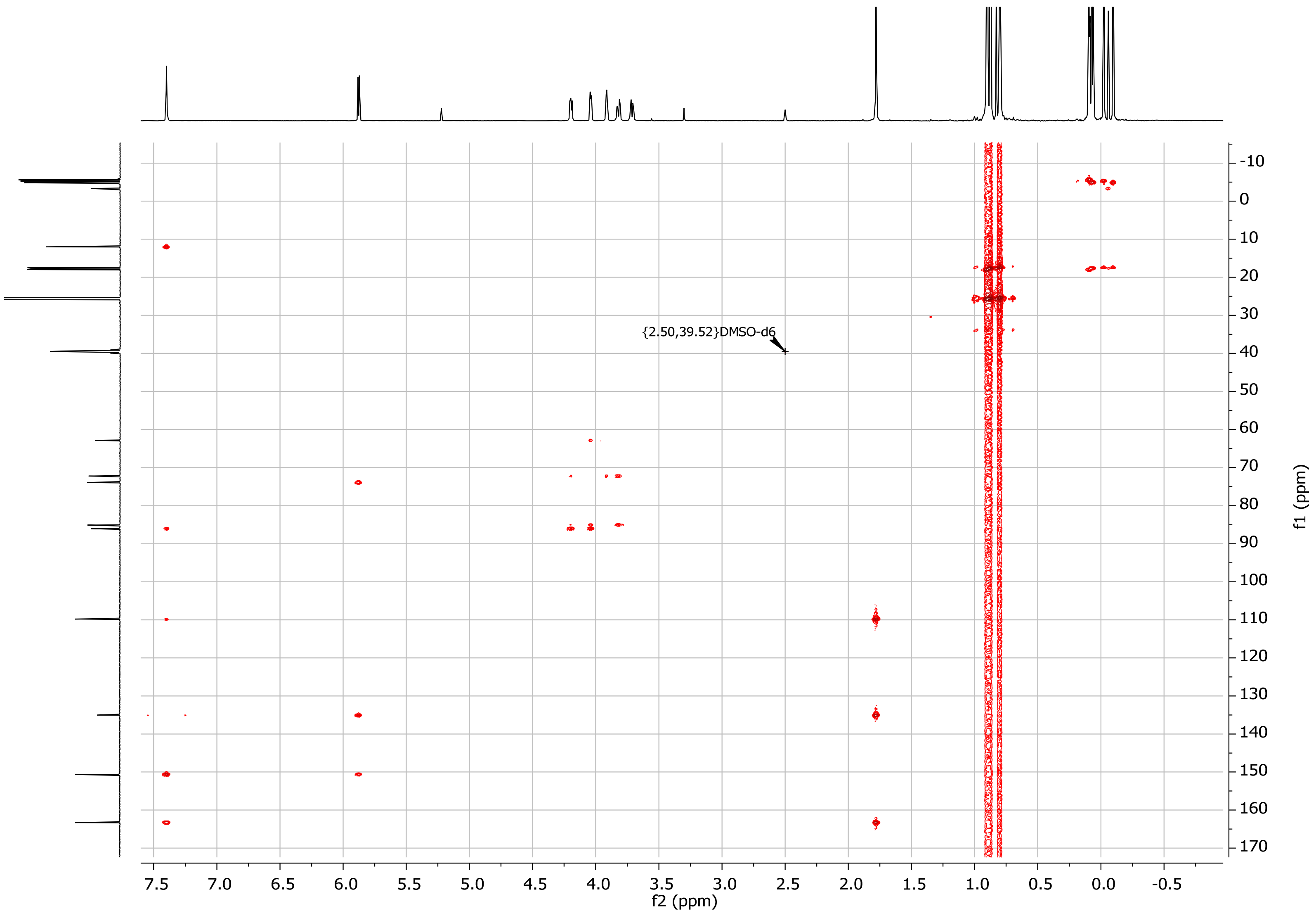

Supplementary Figure 30: HMBC spectrum of compound 1.

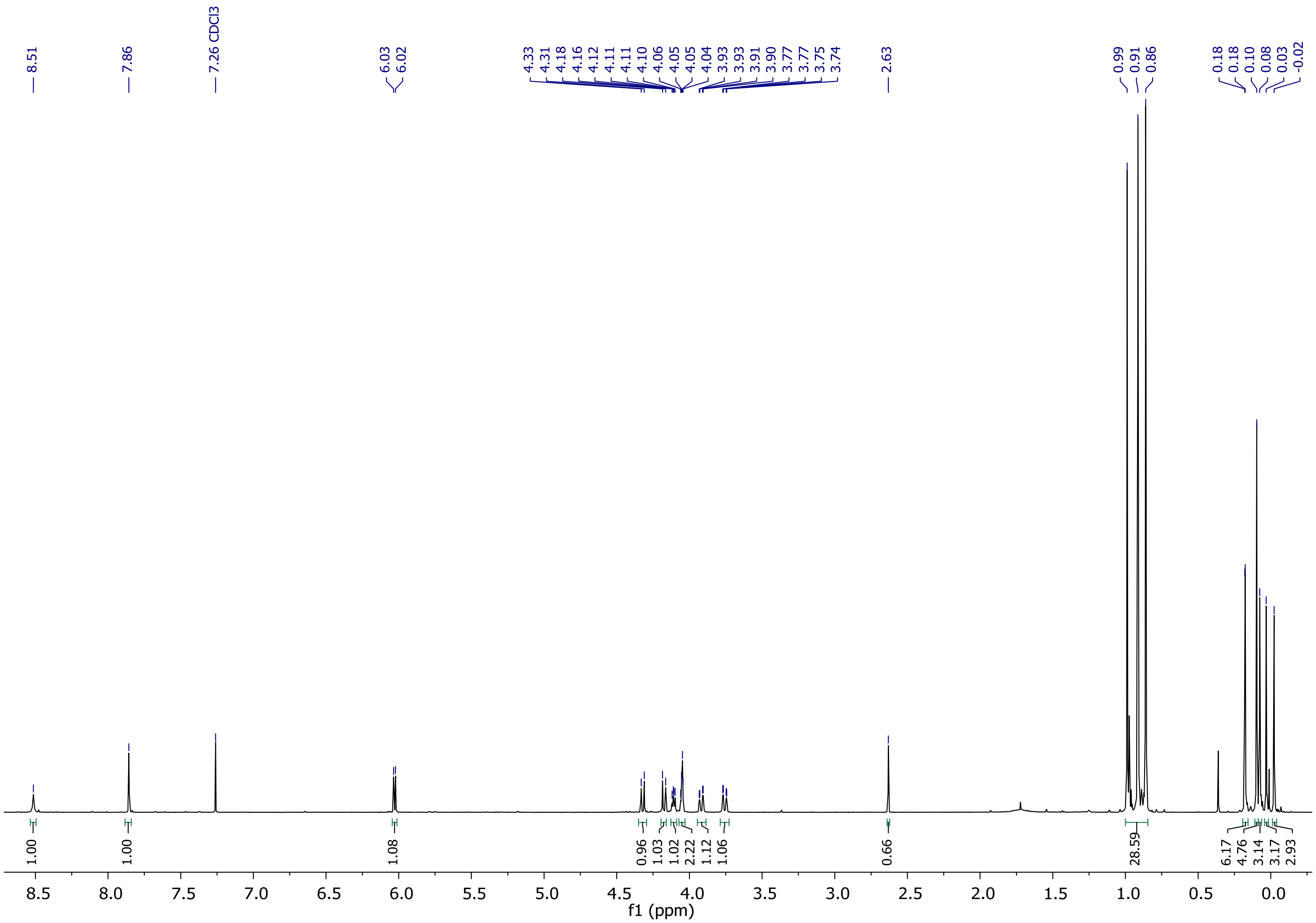

Supplementary Figure 31: $^{1}$H-NMR spectrum of compound 2.

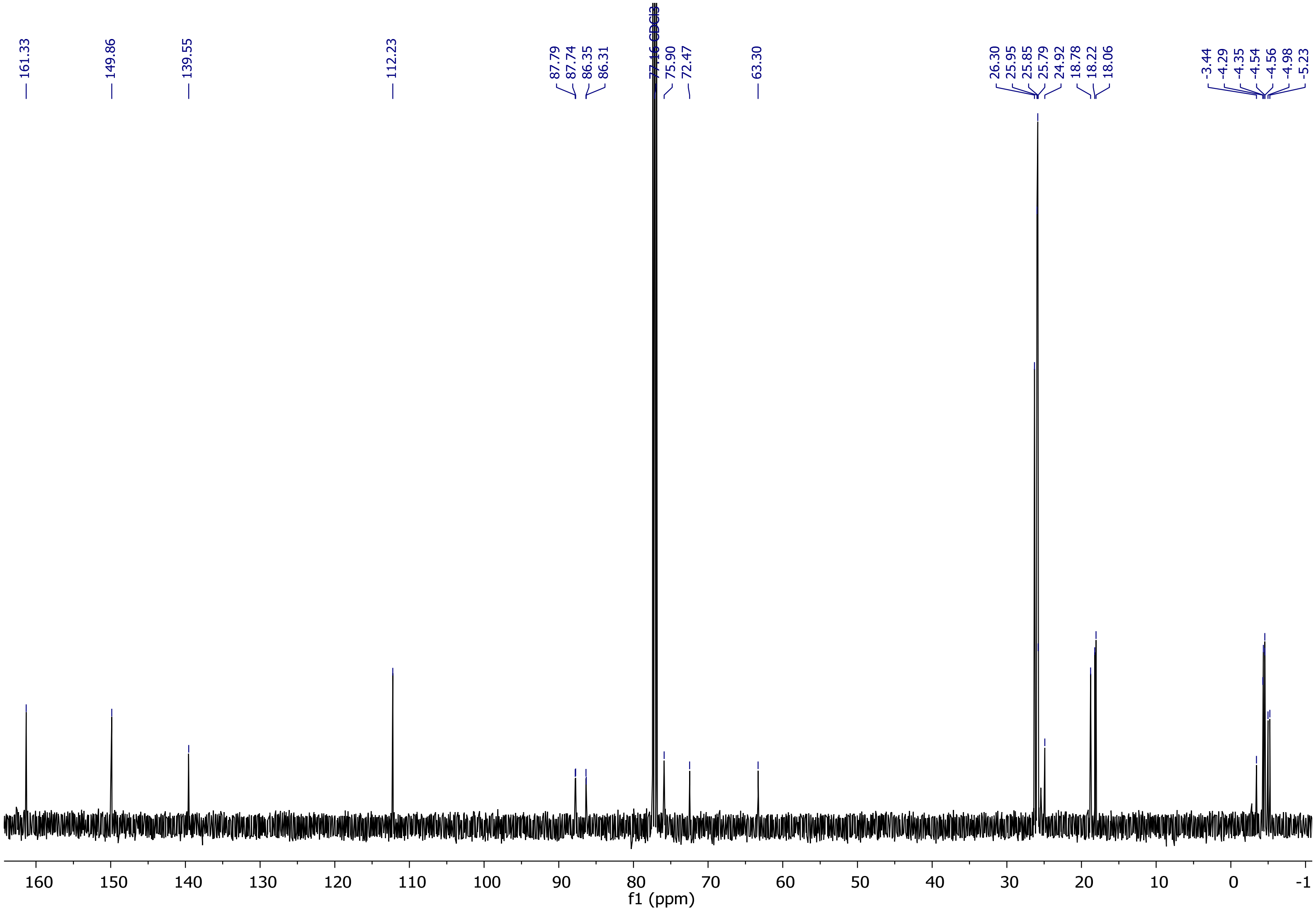

Supplementary Figure 32: $^{13}$C-NMR spectrum of compound 2.

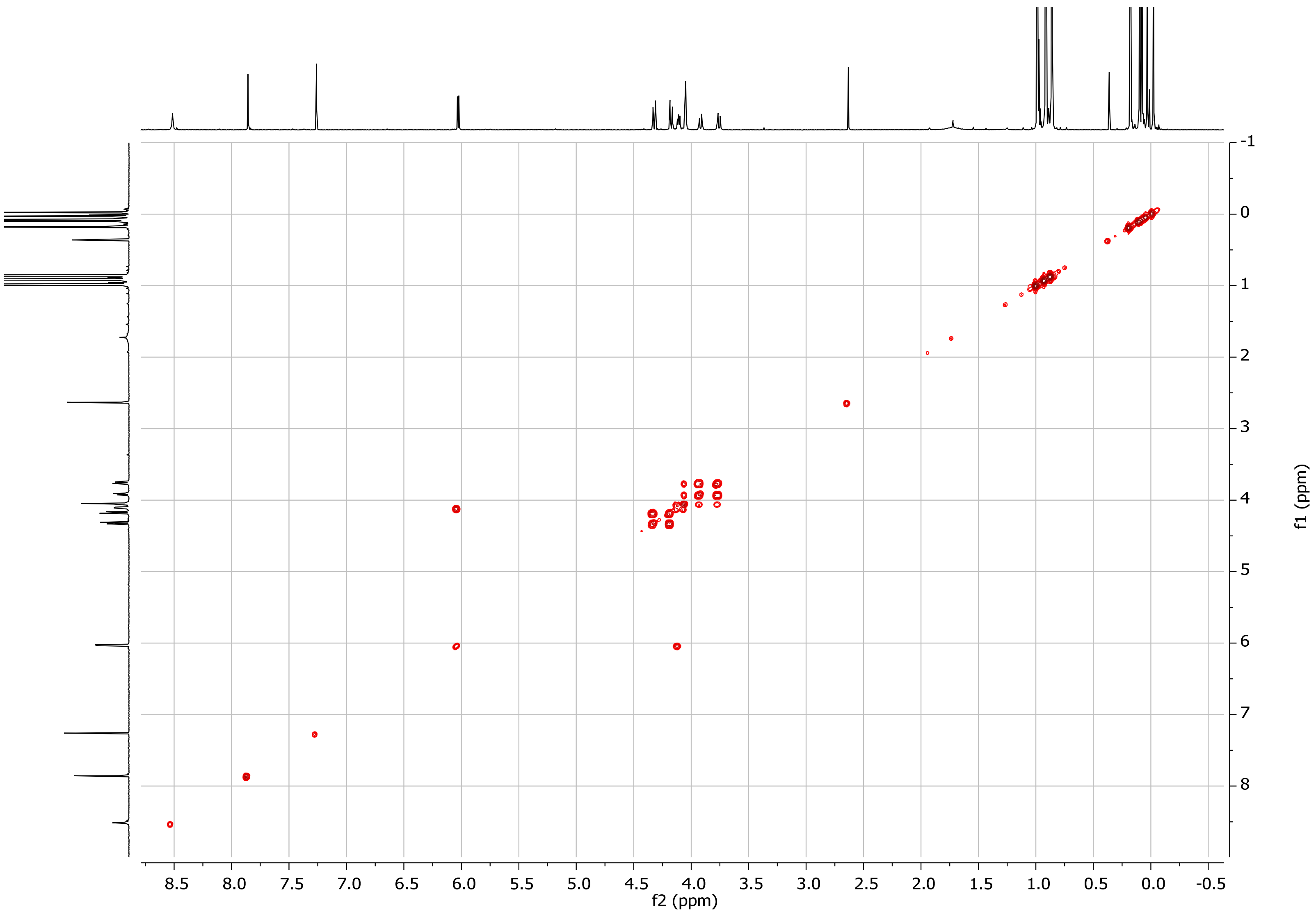

Supplementary Figure 33: COSY spectrum of compound 2.

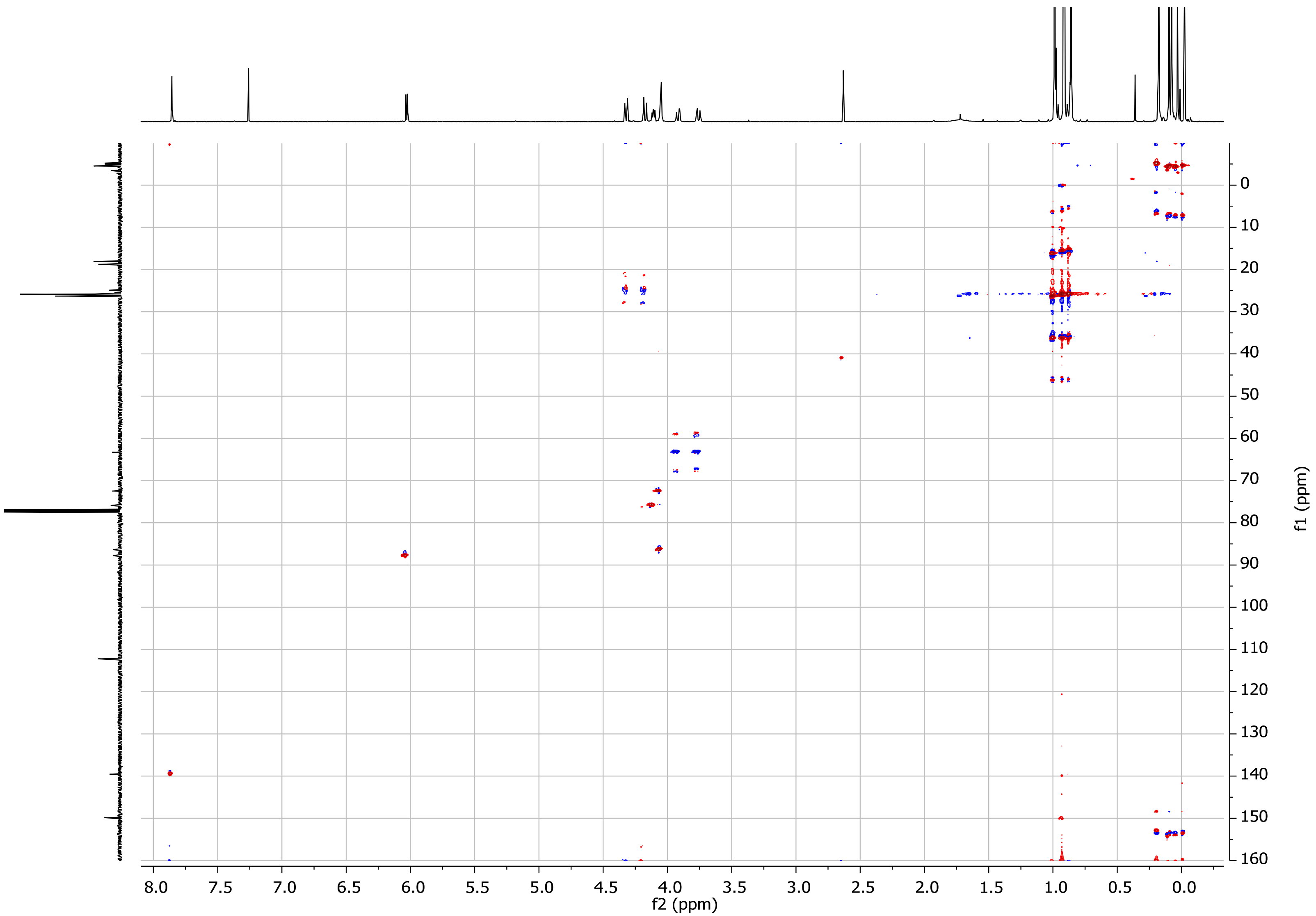

Supplementary Figure 34: HSQC spectrum of compound 2.

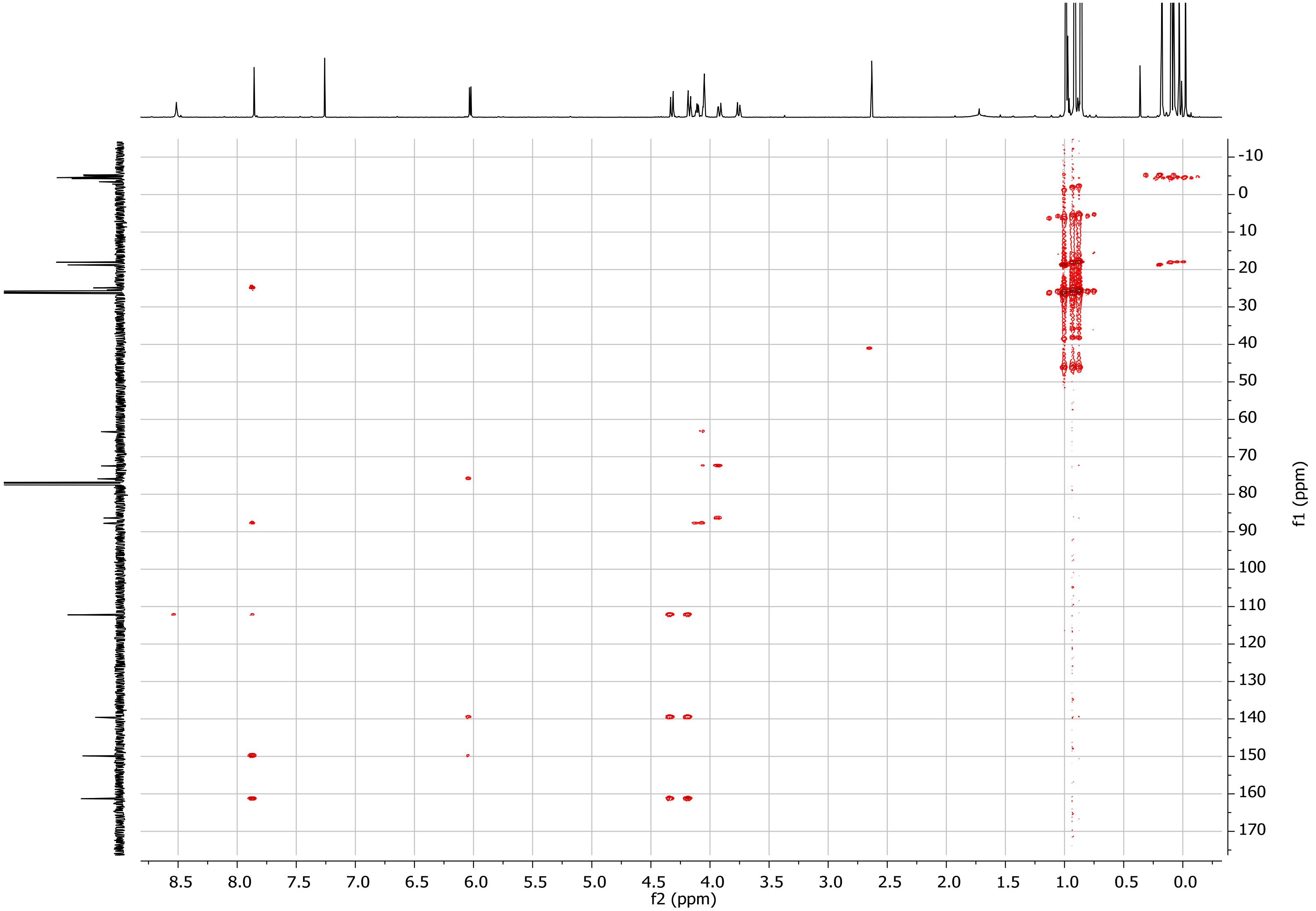

Supplementary Figure 35: HMBC spectrum of compound 2.

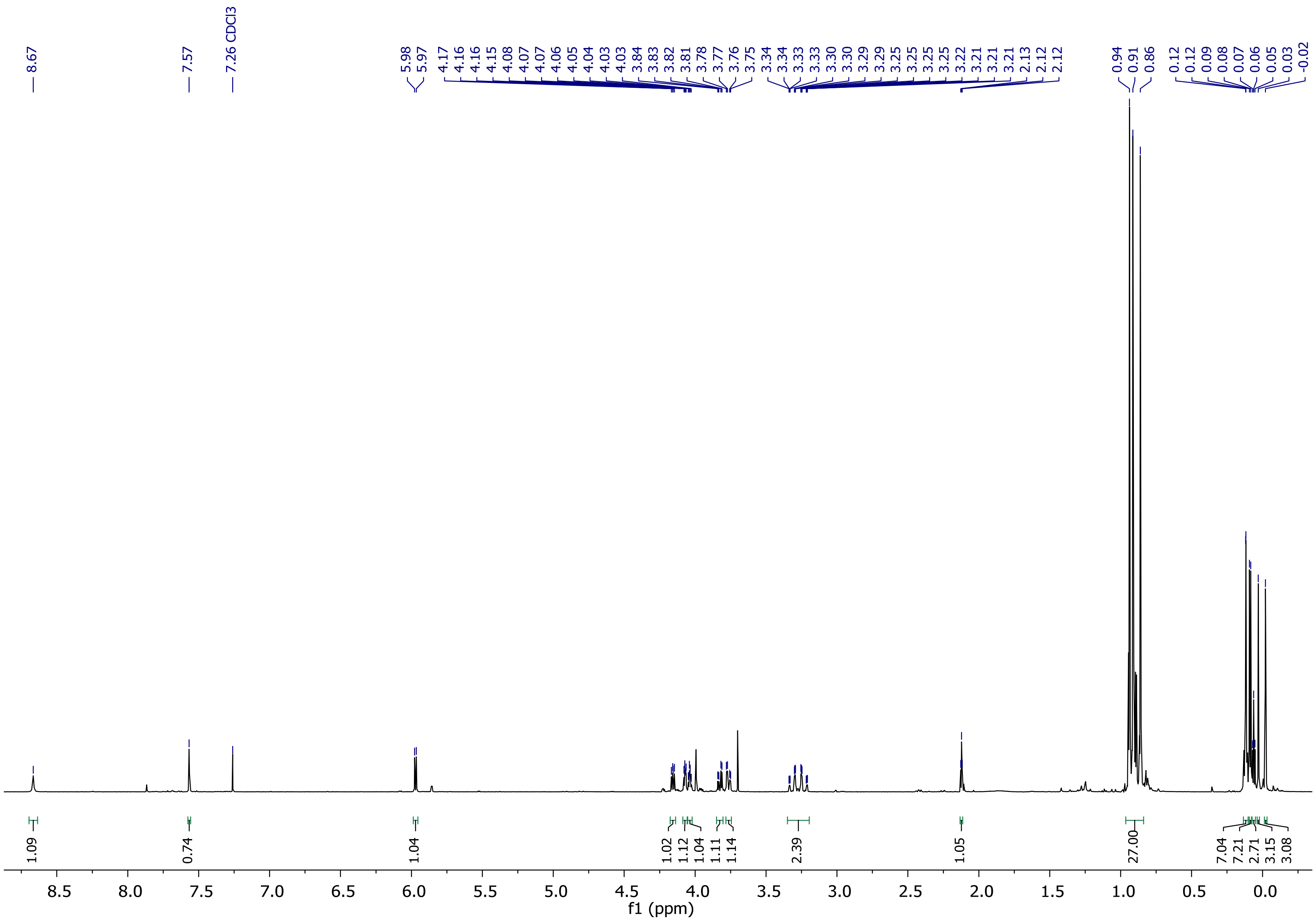

Supplementary Figure 36: $^{1}$H-NMR spectrum of compound 3.

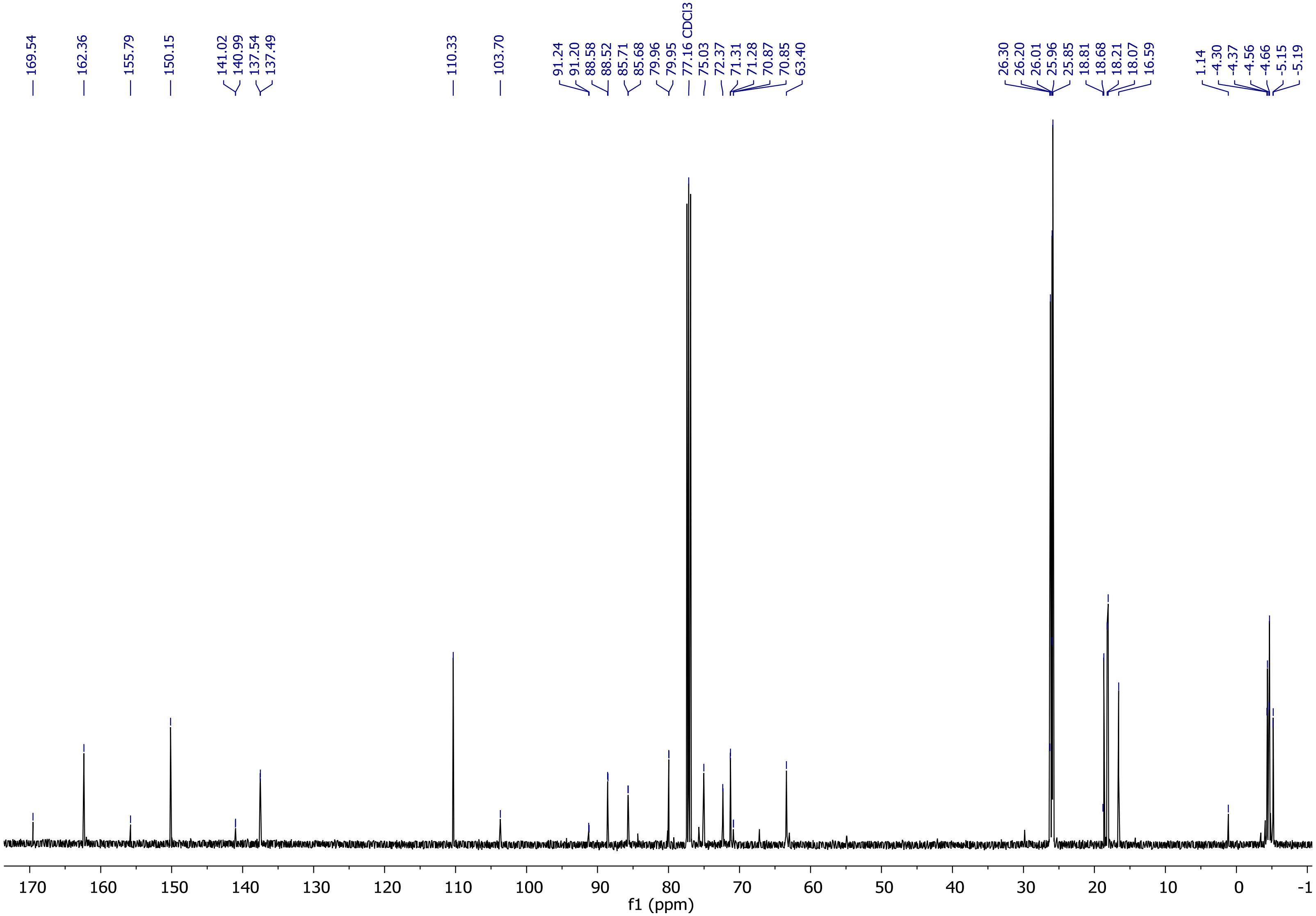

Supplementary Figure 37: $^{13}$C-NMR spectrum of compound 3.

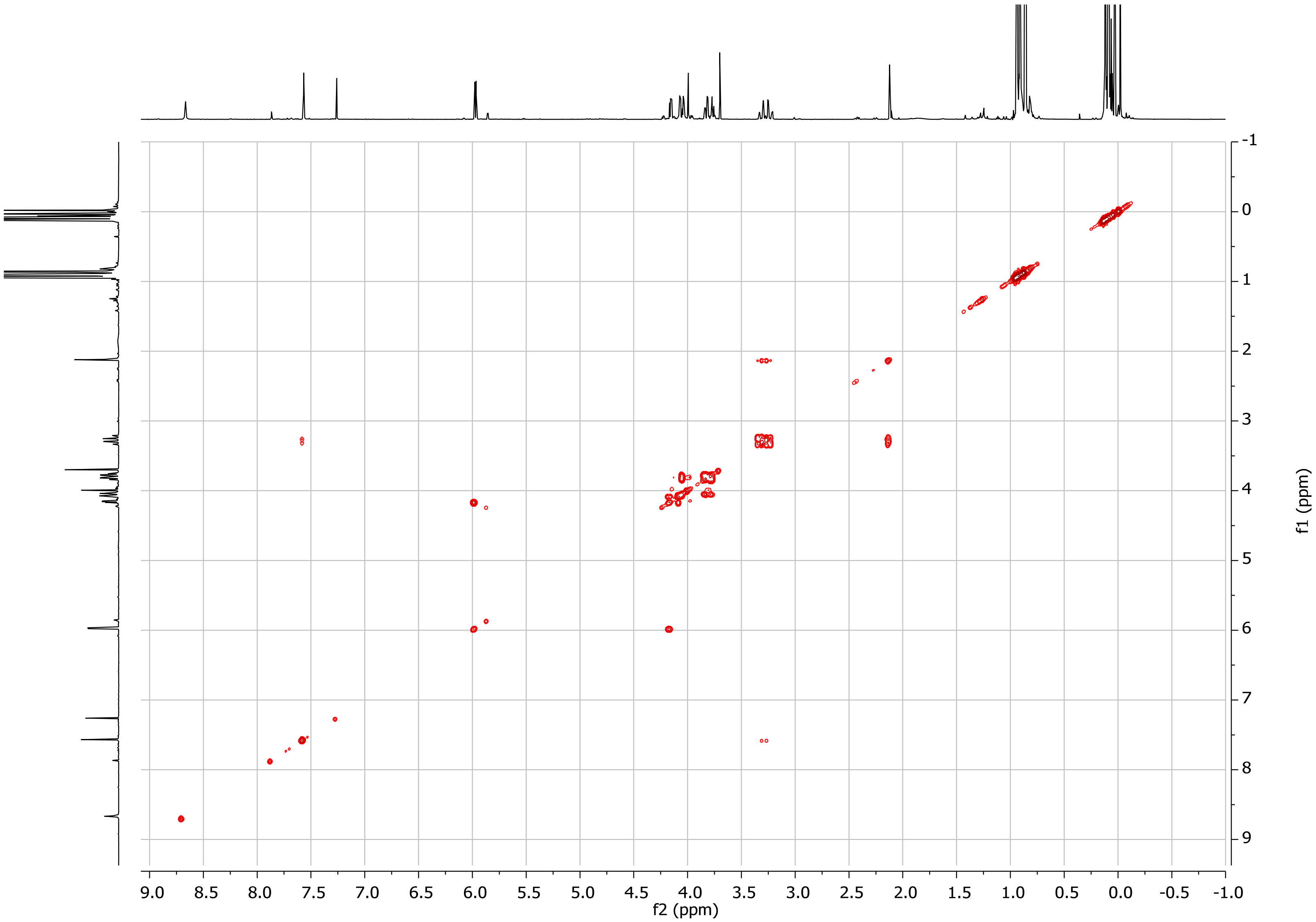

Supplementary Figure 38: COSY spectrum of compound 3.

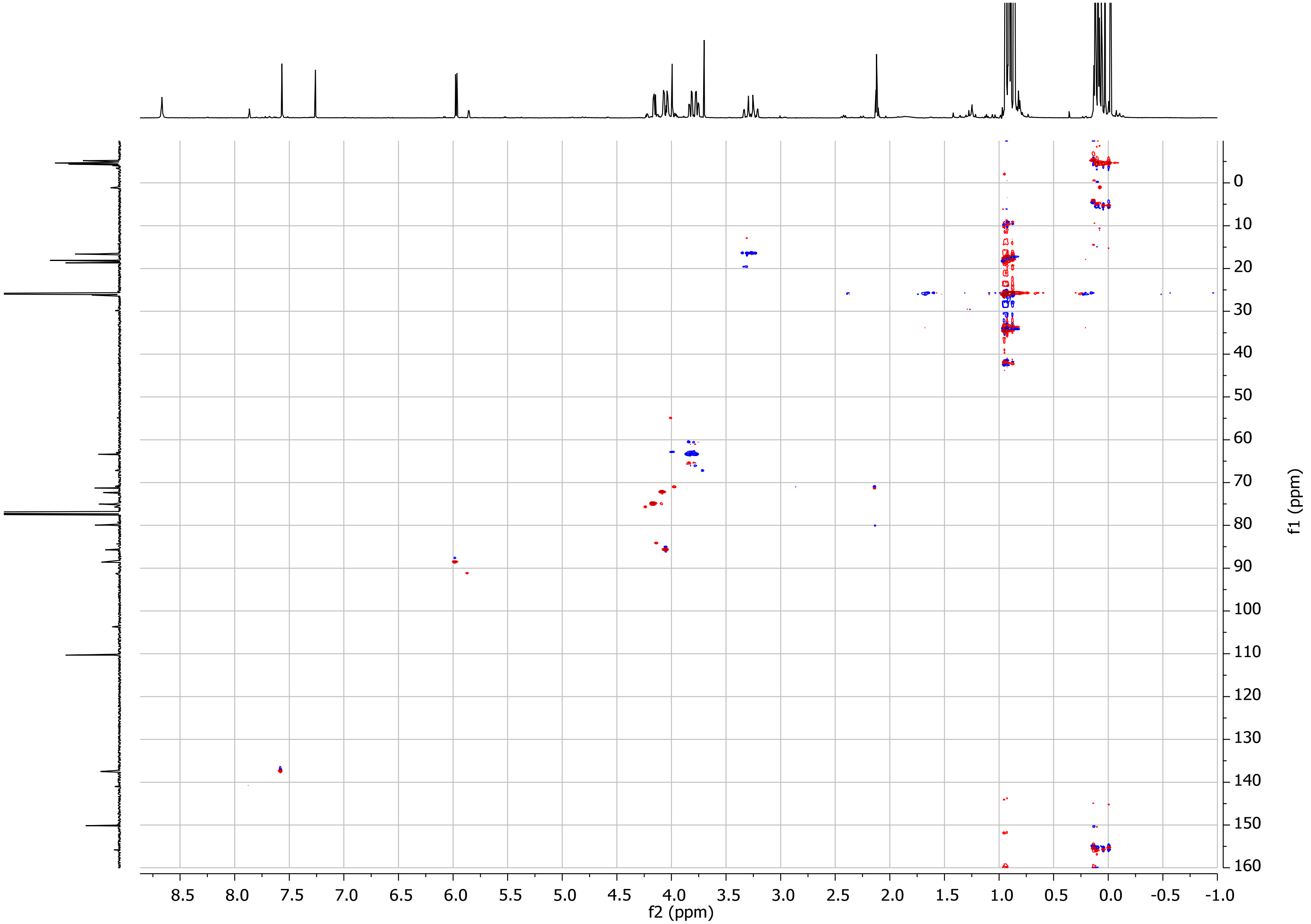

Supplementary Figure 39: HSQC spectrum of compound 3.

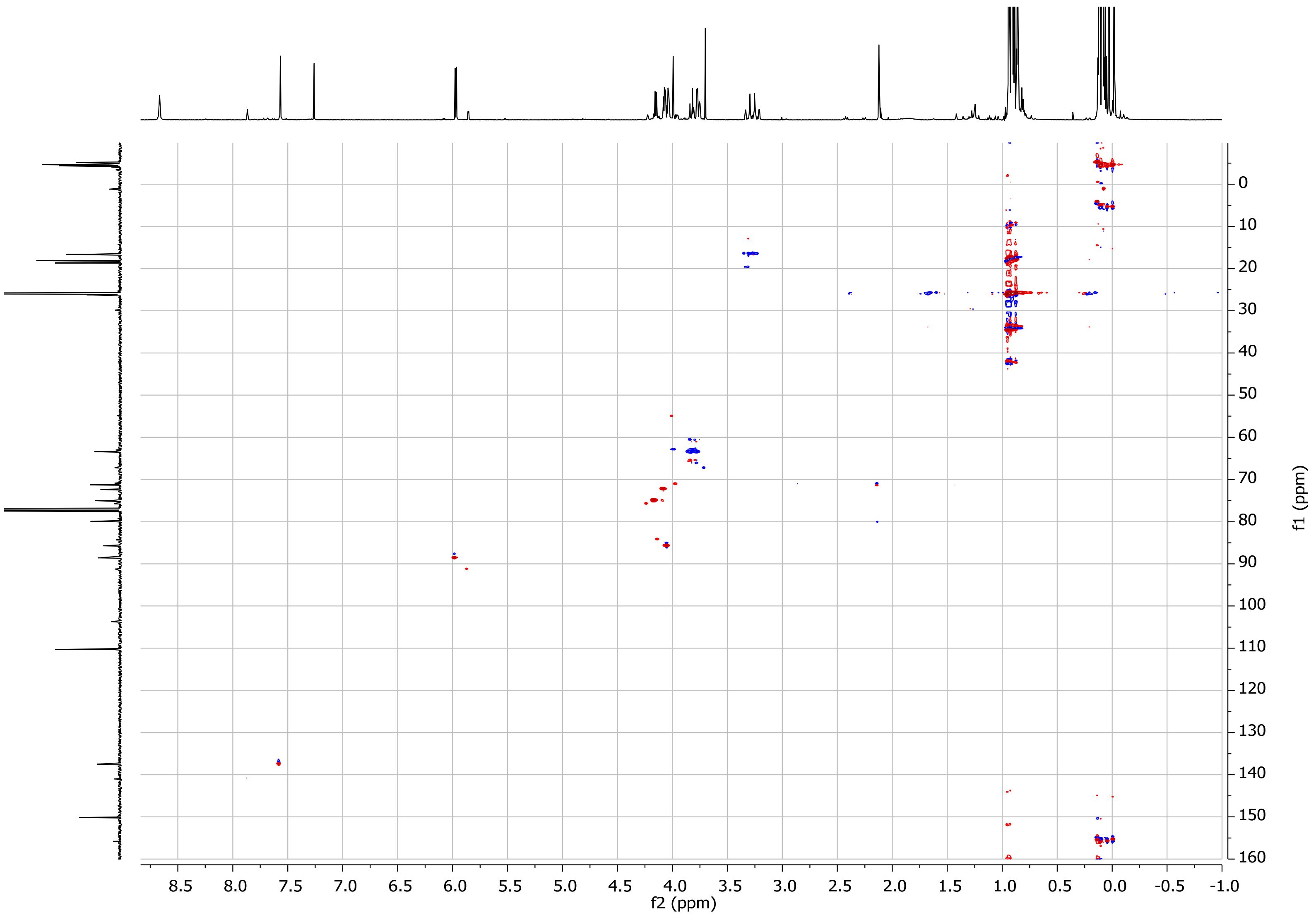

Supplementary Figure 40: HMBC spectrum of compound 3.

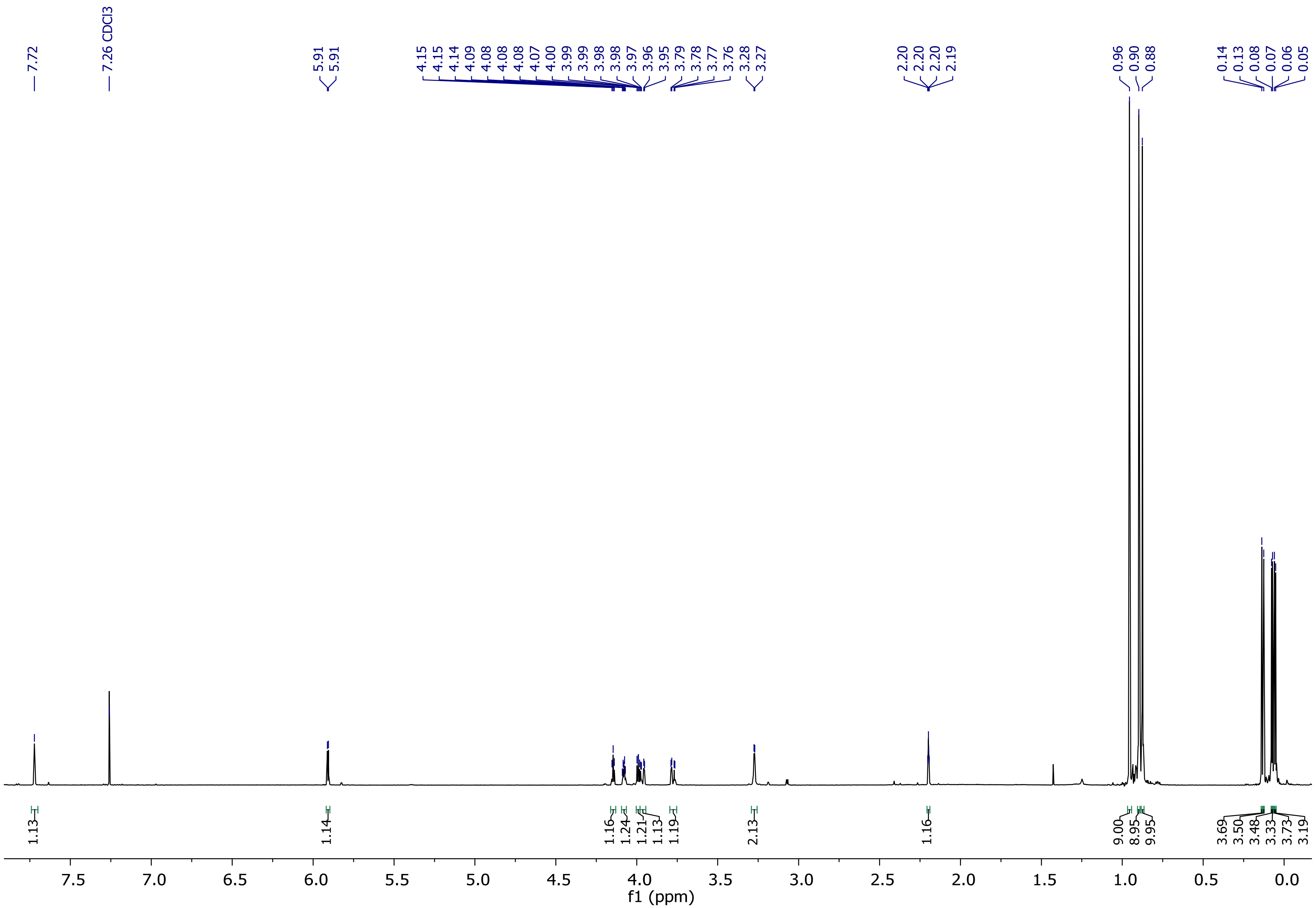

Supplementary Figure 41: $^{1}$H-NMR spectrum of compound 4.

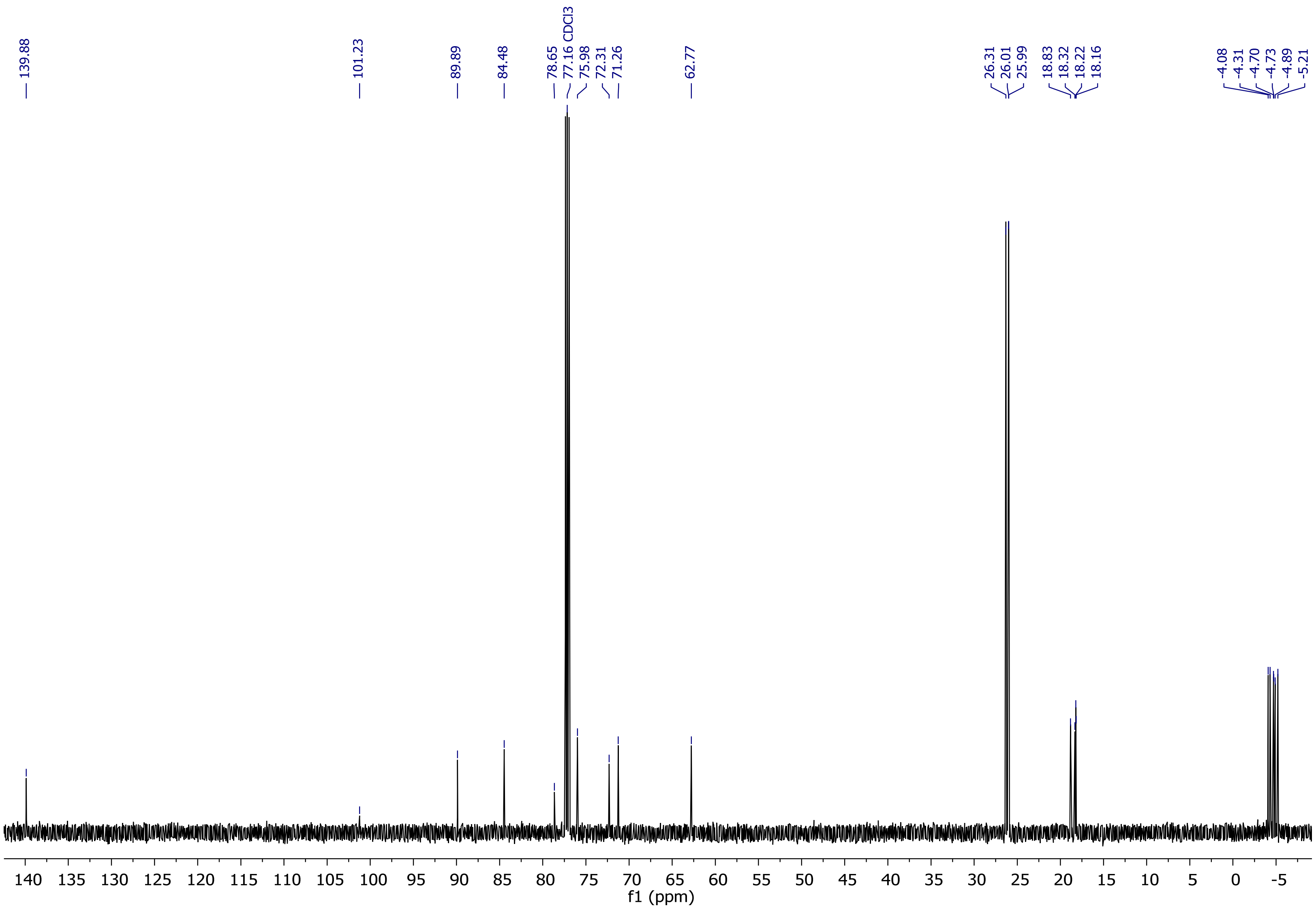

Supplementary Figure 42: $^{13}$C-NMR spectrum of compound 4.

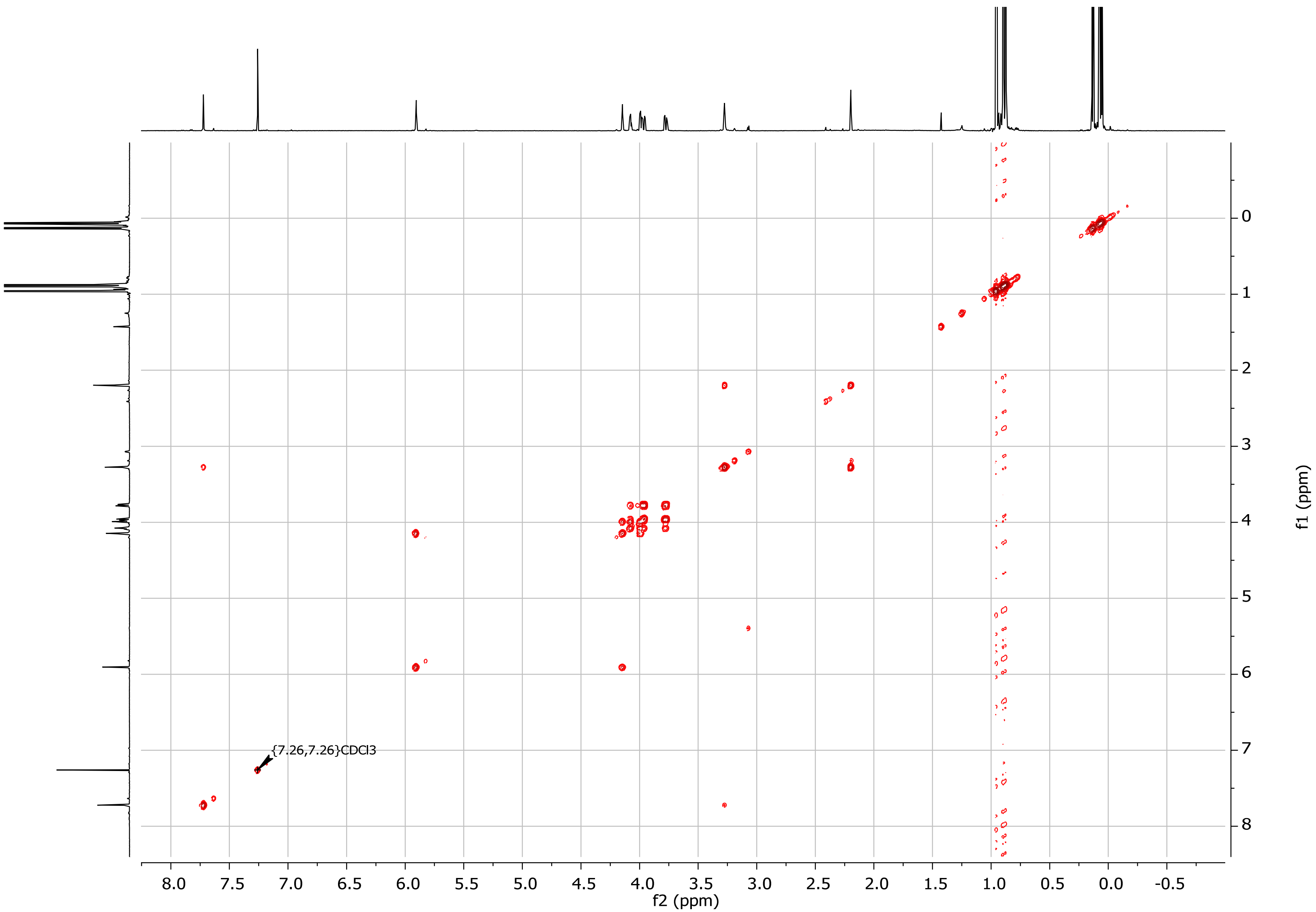

Supplementary Figure 43: COSY spectrum of compound 4.

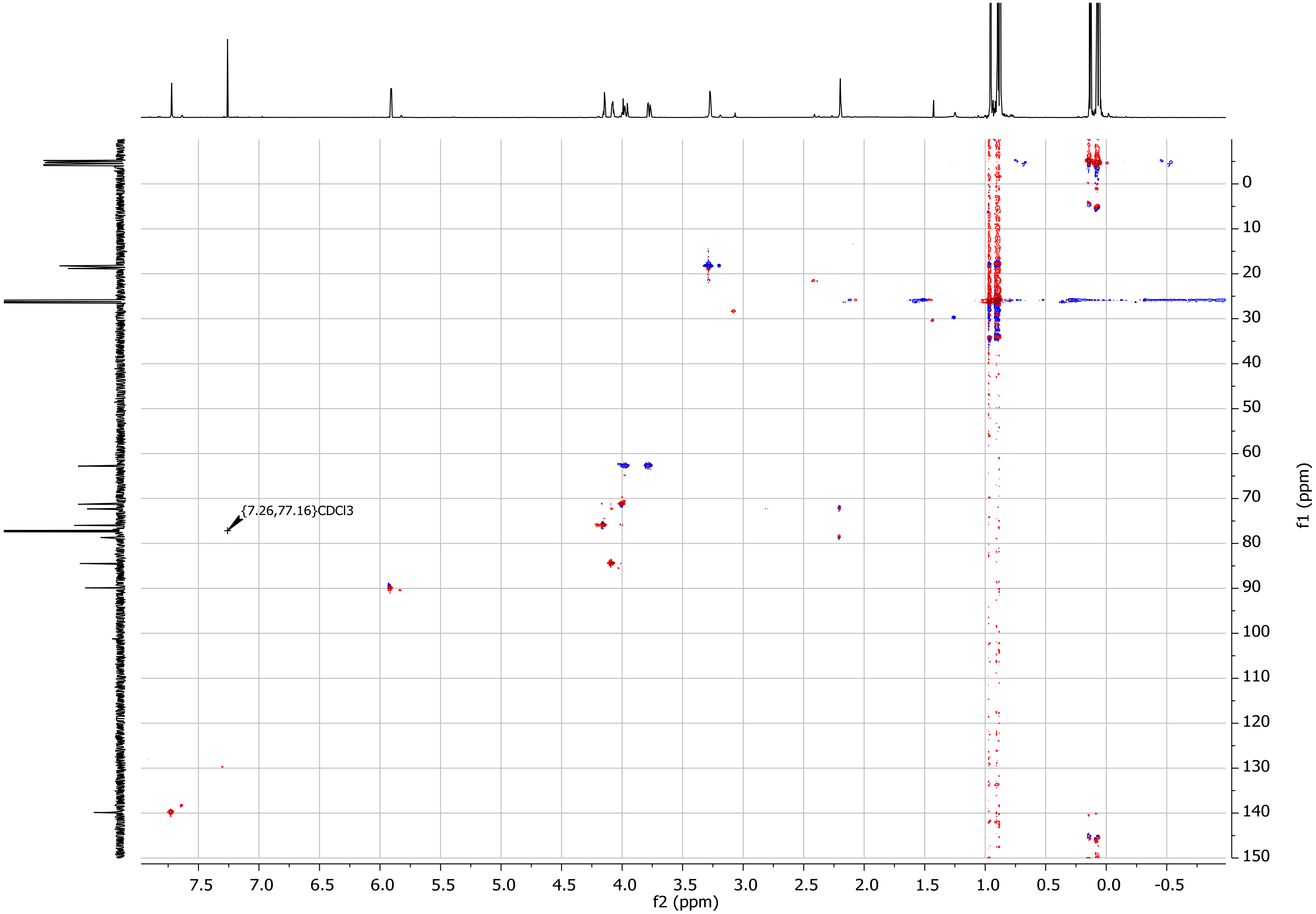

Supplementary Figure 44: HSQC spectrum of compound 4.

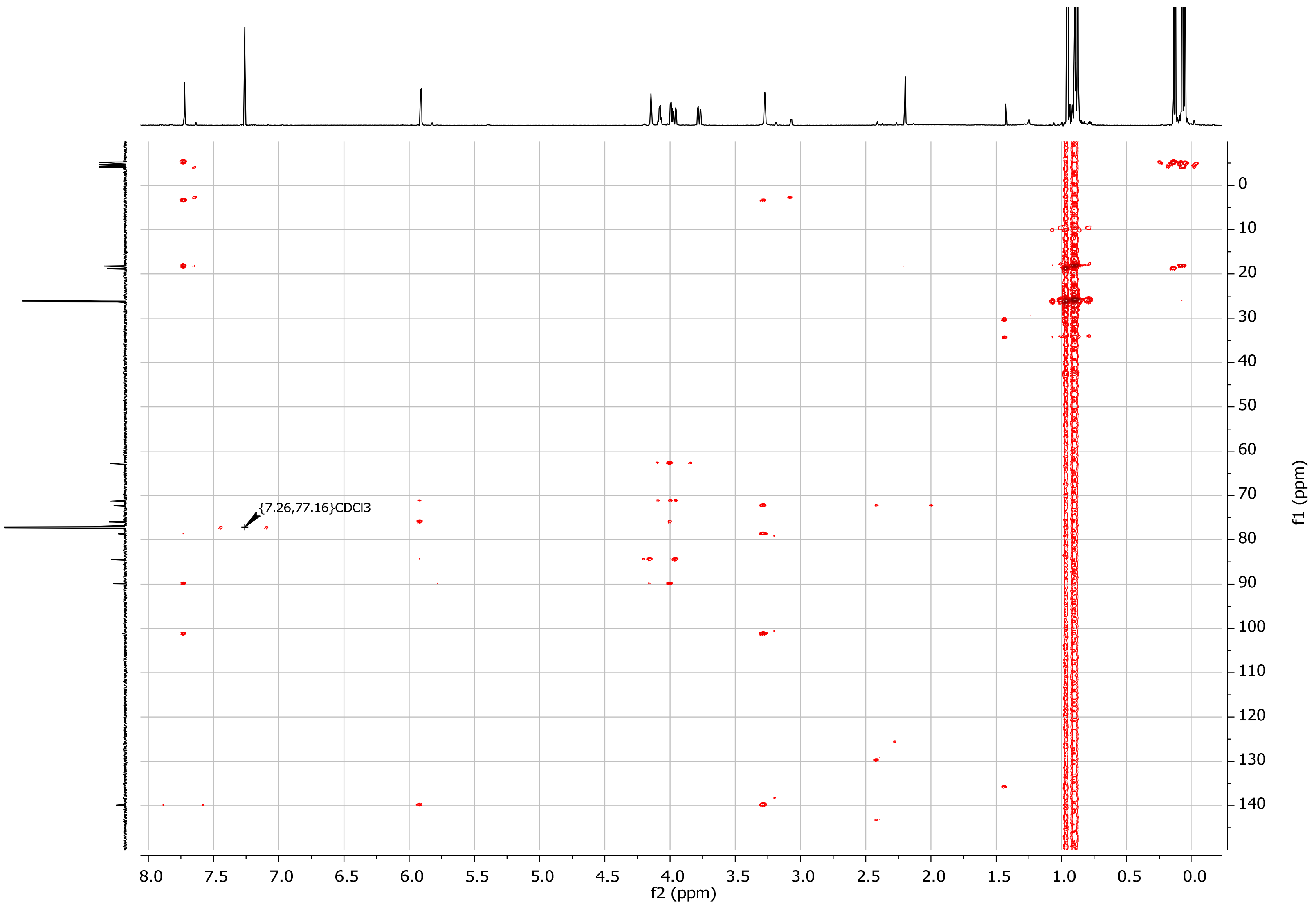

Supplementary Figure 45: HMBC spectrum of compound 4.

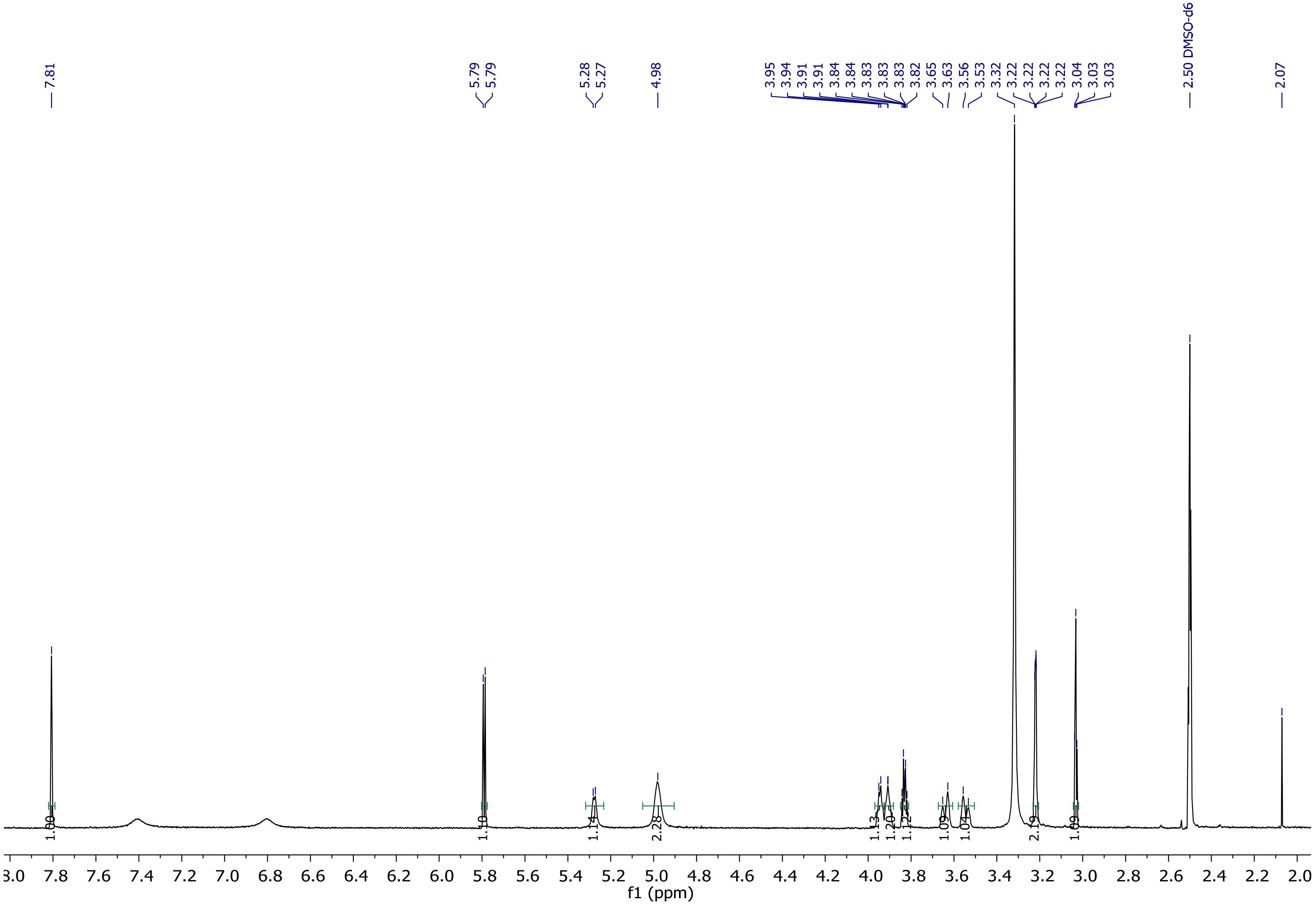

Supplementary Figure 46: $^{1}$H-NMR spectrum of compound 5.

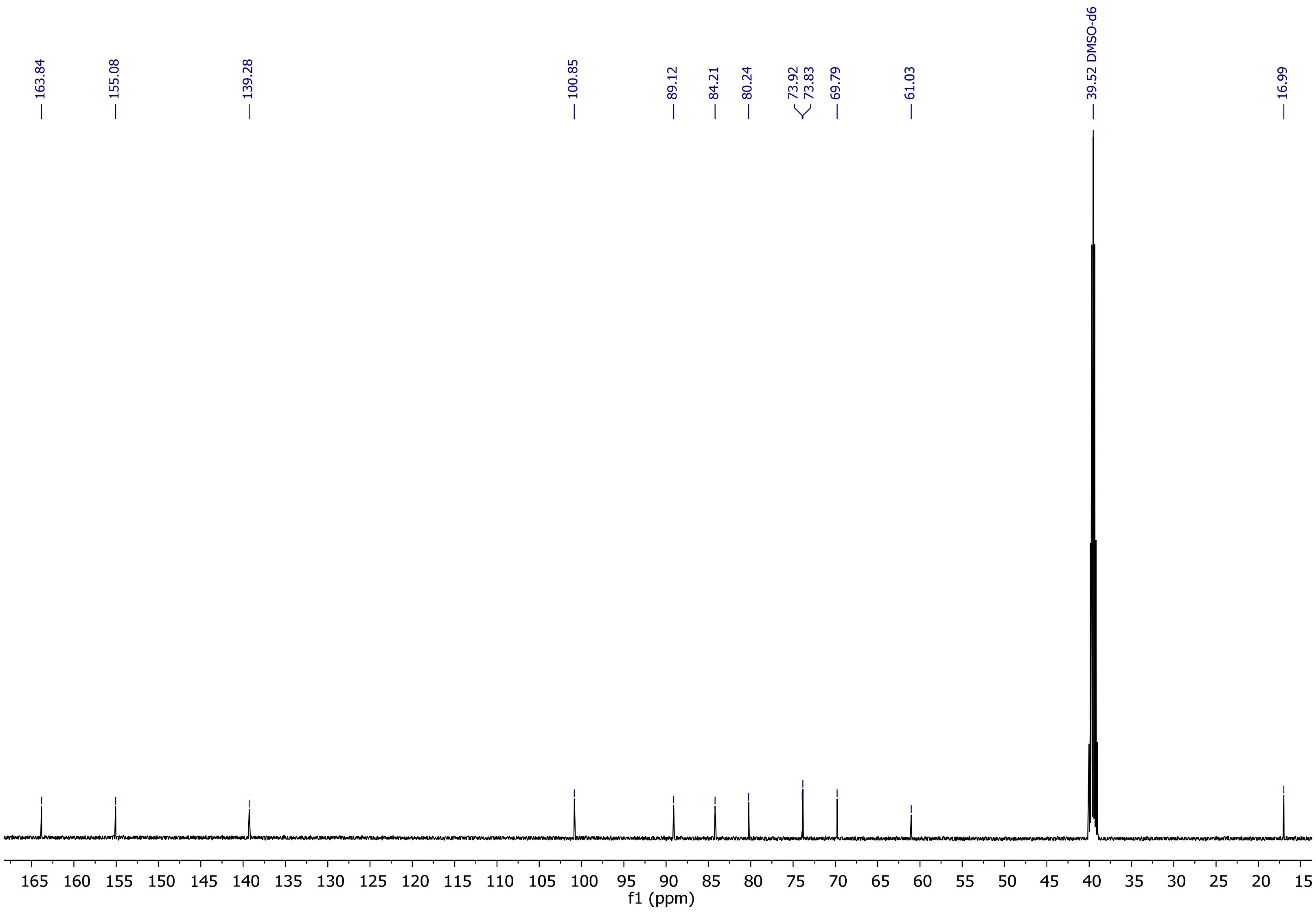

Supplementary Figure 47: $^{13}$C-NMR spectrum of compound 5.

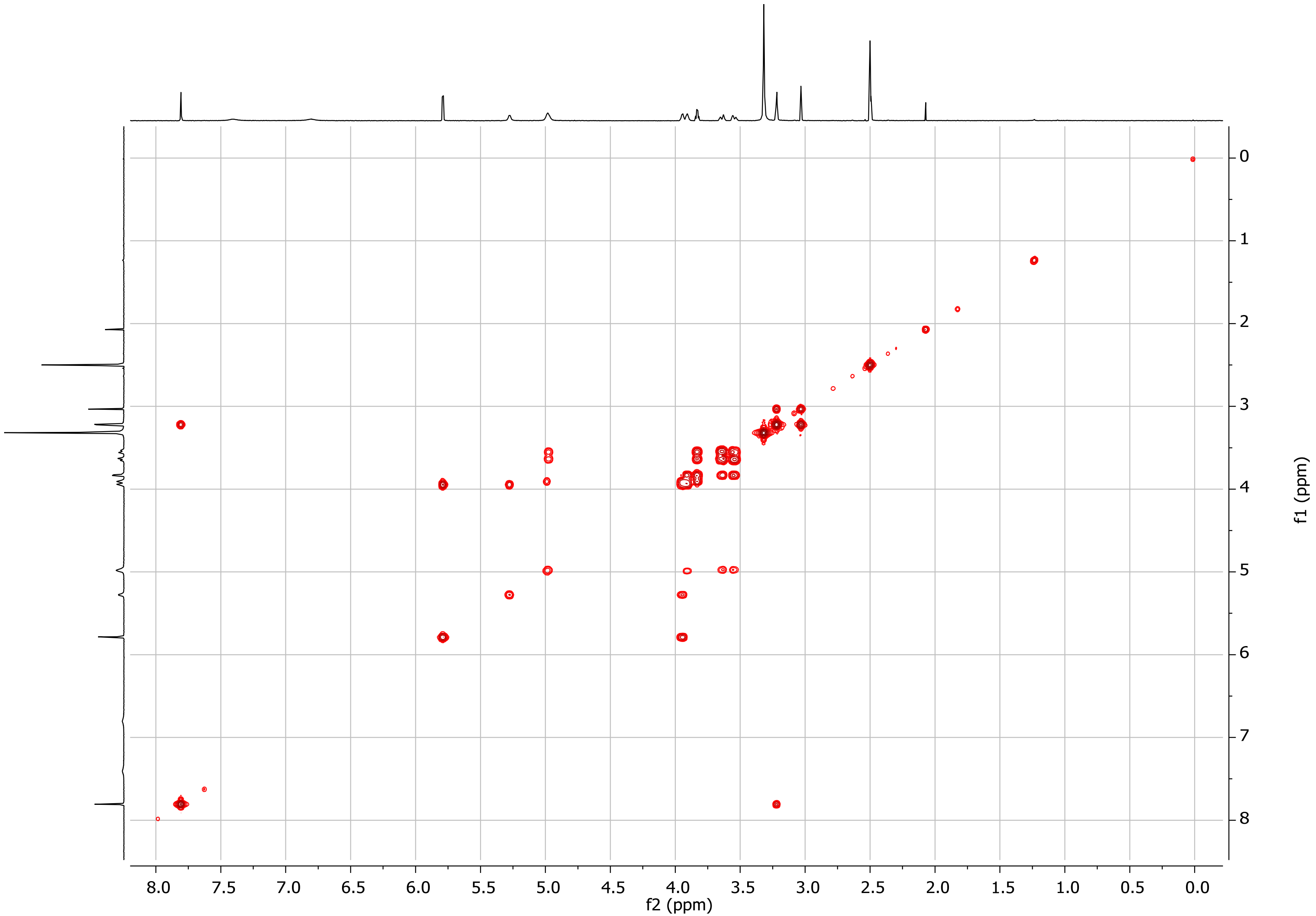

Supplementary Figure 48: COSY spectrum of compound 5.

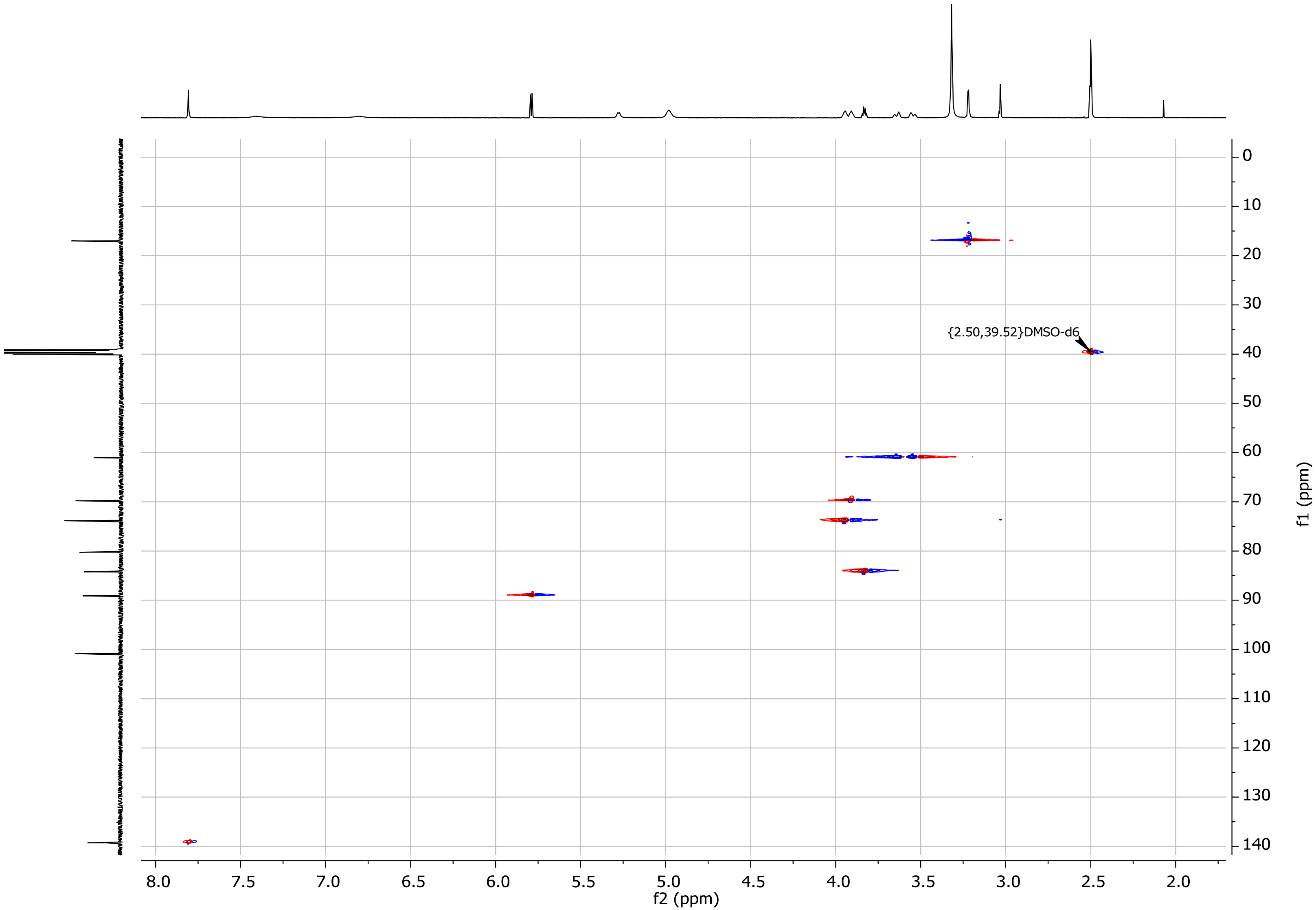

Supplementary Figure 49: HSQC spectrum of compound 5.

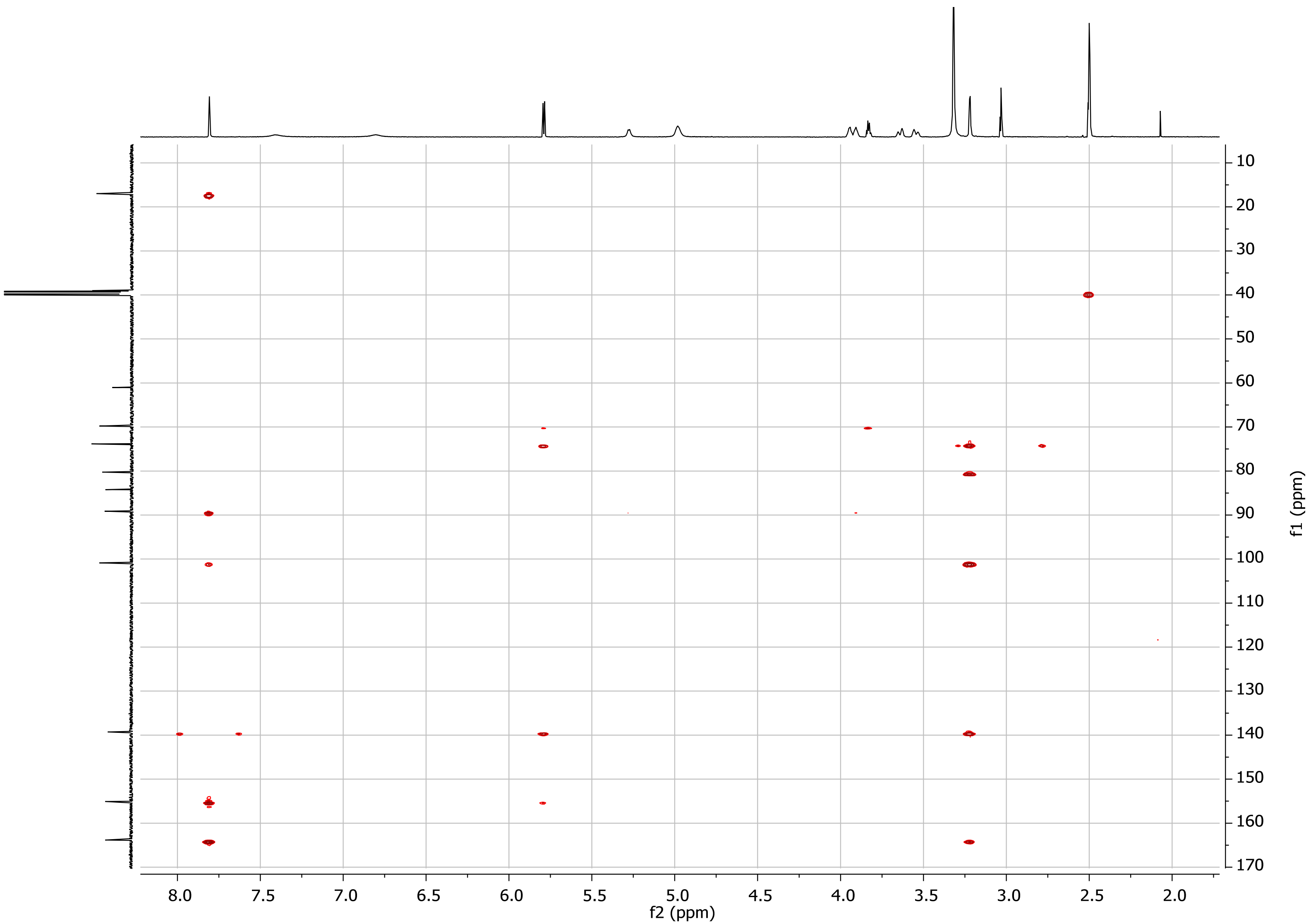

Supplementary Figure 50: HMBC spectrum of compound

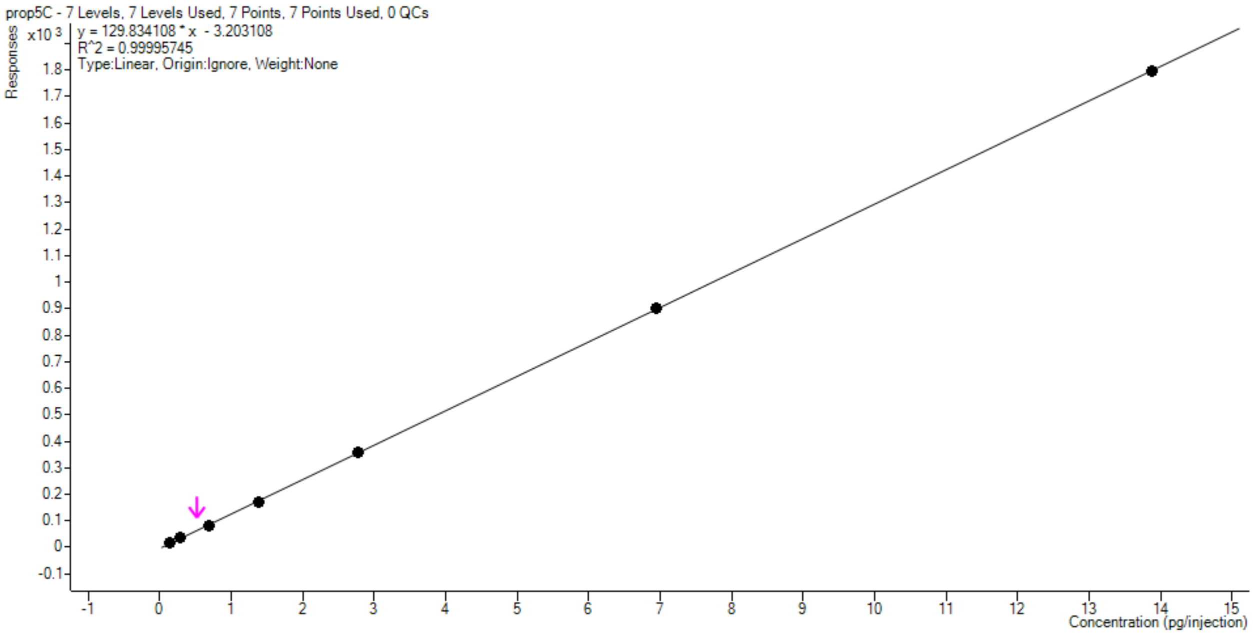

Supplementary Figure 51: The calibration curve of prop$^5$C standard The concentration of prop5C detected in samples from RNA isolated after metabolic labeling with PSH is indicated by the pink arrow and represents one example of n=3 biological replicates.

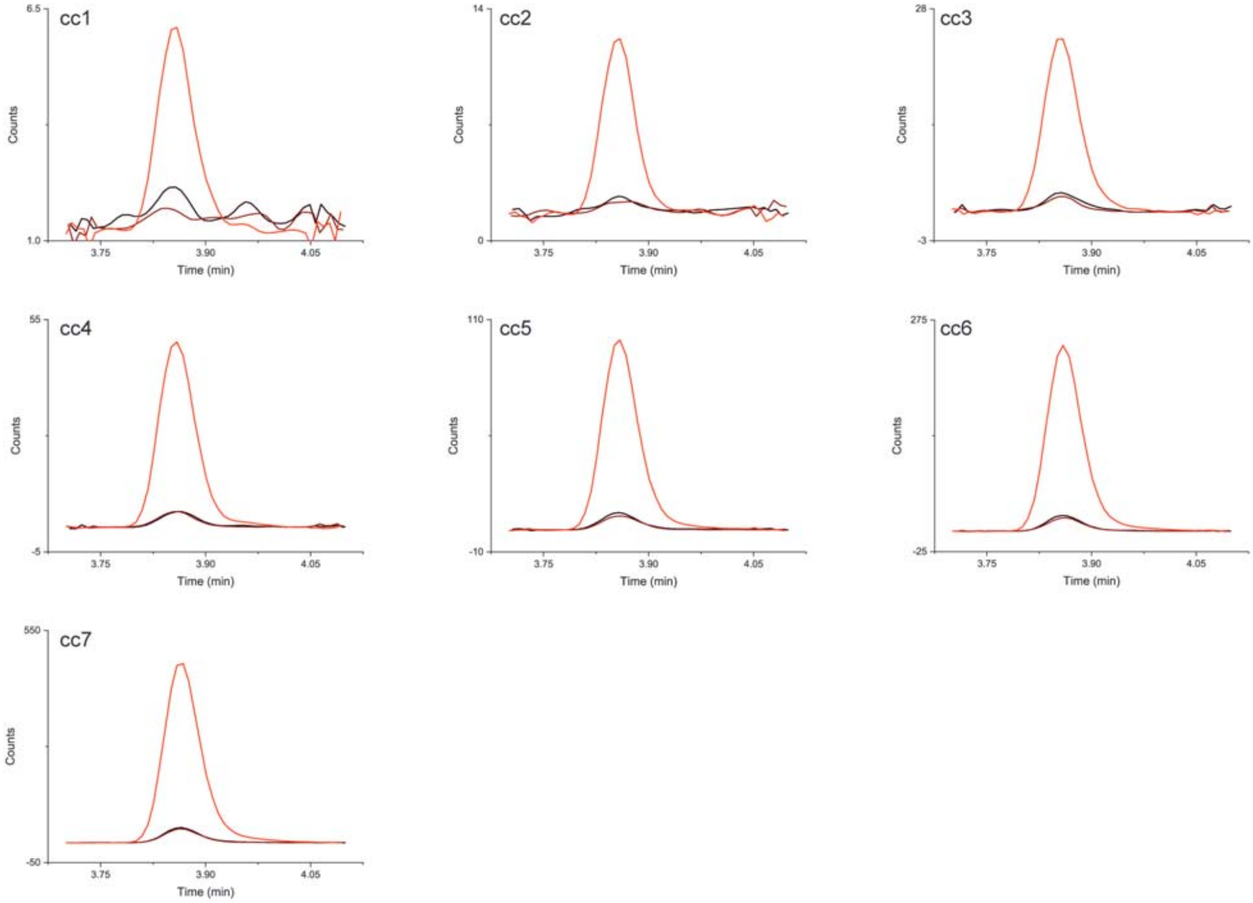

Supplementary Figure 52: The representation of all points of calibration (cc1 to cc7) curve of prop$^5$C. Quantifier (red line) and 2 qualifiers (brown and black lines) are shown.

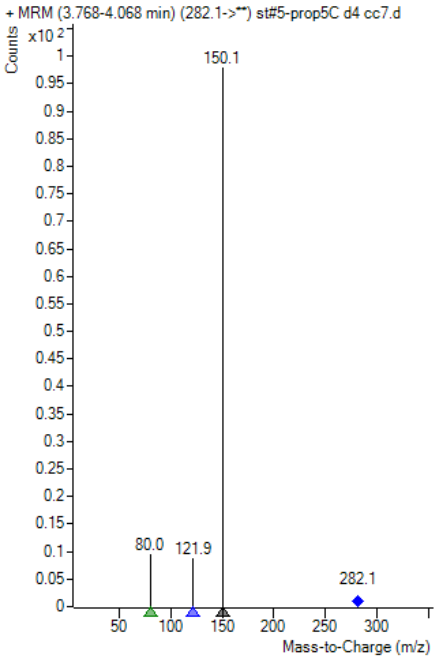

Supplementary Figure 53: The signal intensity of different MRM transition ions of prop$^5$C. The MRM 282.1 → 150.1 (quantifier) is much more abundant than the MRM 282.1 → 121.9 or 282.1 → 80.0 (qualifiers). The signal is from highest point (cc7) of calibration curve (10 µL injection = 13.9 pg of prop$^5$C).

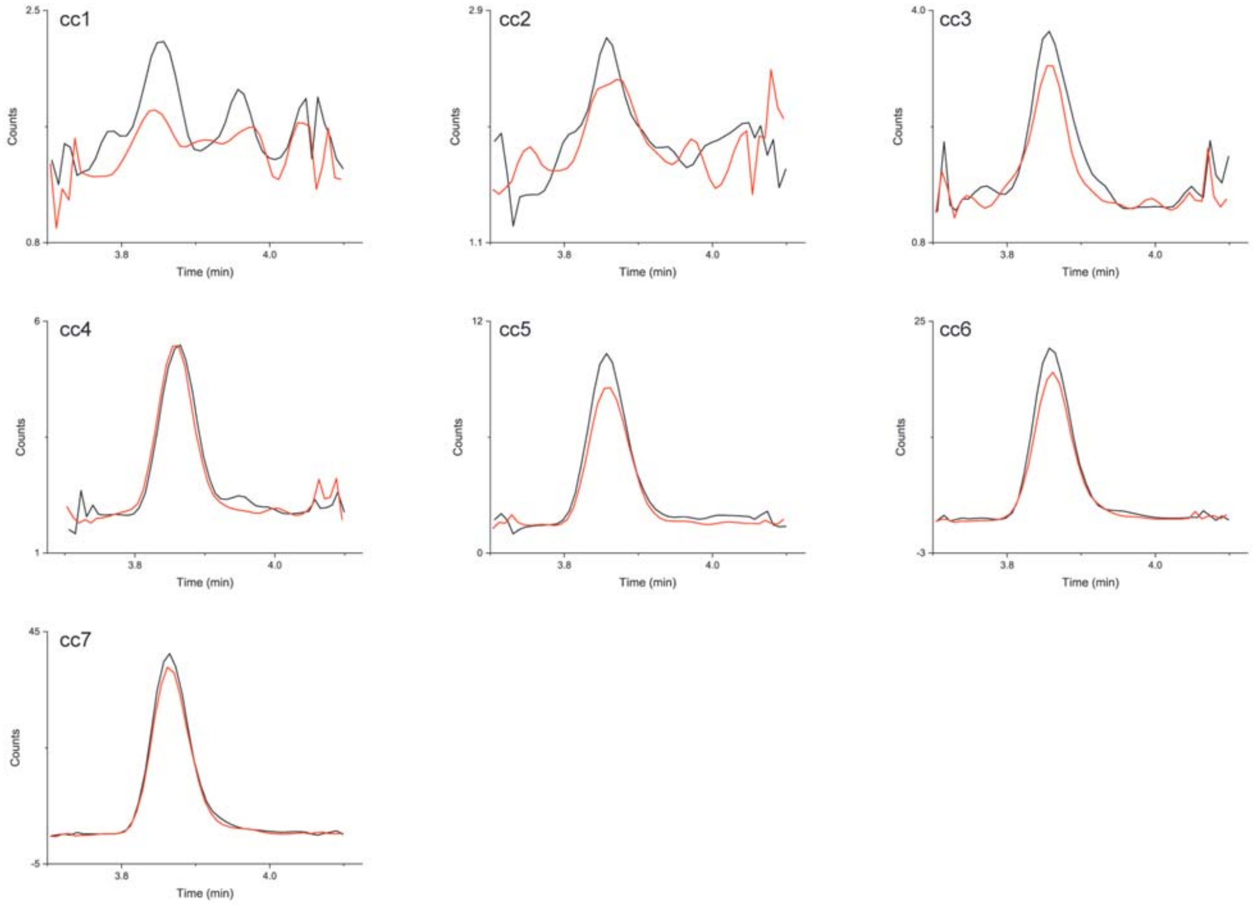

Supplementary Figure 54: Quantifier peaks of calibration curve of prop[5]CMagnification of the qualifiers from Supplementary Fig. 52. MRM 282.1 → 80.0 (black line) and 282.1 → 121.9 (red line). The figure shows the collapse of qualifier peaks between points 3 (cc3) and 2 (cc2) of calibration curve.

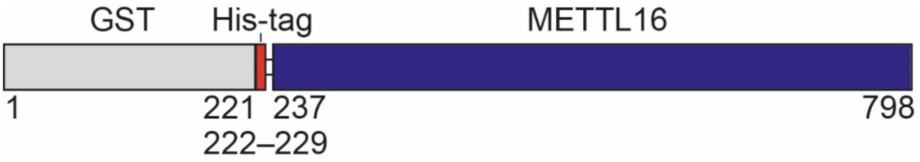

```
        10         20         30         40         50         60
MSPILGYWKI KGLVQPTRLL LEYLEEKYEE HLYERDEGDK WRNKKFELGL EFPNLPYYID

        70         80         90        100        110        120
GDVKLTQSMA IIRYIADKHN MLGGCPKERA EISMLEGAVL DIRYGVSRIA YSKDFETLKV

       130        140        150        160        170        180
DFLSKLPEML KMFEDRLCHK TYLNGDHVTH PDFMLYDALD VVLYMDPMCL DAFPKLVCFK

       190        200        210        220        230        240
KRIEAIPQID KYLKSSKYIA WPLQGWQATF GGGDHPPKSG SHHHHHHHHE NLYFQGMALS

       250        260        270        280        290        300
KSMHARNRYK DKPPDFAYLA SKYPDFKQHV QINLNGRVSL NFKDPEAVRA LTCTLLREDF

       310        320        330        340        350        360
GLSIDIPLER LIPTVPLRLN YIHWVEDLIG HQDSDKSTLR RGIDIGTGAS CIYPLLGATL

       370        380        390        400        410        420
NGWYFLATEV DDMCFNYAKK NVEQNNLSDL IKVVKVPQKT LLMDALKEES EIIYDFCMCN

       430        440        450        460        470        480
PPFFANQLEA KGVNSRNPRR PPPSSVNTGG ITEIMAEGGE LEFVKRIIHD SLQLKKRLRW

       490        500        510        520        530        540
YSCMLGKKCS LAPLKEELRI QGVPKVTYTE FCQGRTMRWA LAWSFYDDVT VPSPPSKRRK

       550        560        570        580        590        600
LEKPRKPITF VVLASVMKEL SLKASPLRSE TAEGIVVVTT WIEKILTDLK VQHKRVPCGK

       610        620        630        640        650        660
EEVSLFLTAI ENSWIHLRRK KRERVRQLRE VPRAPEDVIQ ALEEKKPTPK ESGNSQELAR

       670        680        690        700        710        720
GPQERTPCGP ALREGEAAAV EGPCPSQESL SQEENPEPTE DERSEEKGGV EVLESCQGSS

       730        740        750        760        770        780
NGAQDQEASE QFGSPVAERG KRLPGVAGQY LFKCLINVKK EVDDALVEMH WVEGQNRDLM

       790
NQLCTYIRNQ IFRLVAVN
```

Supplementary Figure 55: Sequence of METTL16-construct used in this work.

### Supplementary Notes

#### Supplementary Note 1: Synthesis of 5-propargylcytidine (prop$^5$C)

Materials and solvents

Unless specified, all reagents were used as supplied (see Supplementary Table 10) without further purification and solvents were evaporated on a rotary evaporator at 40 °C/2 kPa. Technical grade solvents were used for workup of reactions and for flash chromatography. Benzene (Carl Roth, 7173.1), 1.4-dioxane (Merck, 1.03115.1000) and chloroform (Fisher Scientific, C/4960/15) were used in analytical grade. For RP flash chromatography, the dd$H_2O$ and $CH_3CN$ (HPLC grade) as eluents were used. Dry solvents were purchased from the distributers mentioned in Supplementary Table 10.

General instrumentation, chromatography materials and methods for synthesis

$^1$H and $^{13}$C NMR spectra were measured on an Agilent DD2 600 spectrometer (600 MHz and 151 MHz) and an Agilent DD2 500 spectrometer (500 MHz and 126 MHz). The measurements were performed in DMSO-*d6* or $CDCl_3$ and referred to residual solvent signal. Complete assignment (if present) is based on heteronuclear correlation experiments HSQC and H, C-HMBC and COSY. Chemical shifts ($\delta$) are in ppm and coupling constants ($J$) in Hz. The numbering system for the assignment of NMR signals is given for the compounds individually. Spectra can be found as Supplementary Fig. 26-50.

High resolution mass spectra were measured on an Orbitrap LTQ XL (Thermo Fisher Scientific) spectrometer using ESI technique and an Orbitrap Velos Pro (Thermo Fisher Scientific) spectrometer using ESI technique.

For determination of low-resolution mass spectra, an HPLC - triple-quadrupole mass spectrometry (LC-QqQ) system was used. The system consists of an Agilent 1260 Infinity II with dual λ absorbance detector (G7114A) and an Agilent Ultivo mass spectrometer with JetStream ion source (type of ESI). Column: Agilent Poroshell 120 EC-C18 (3.0x150/2.7µm). For the very lipophilic compounds (**1**, **2**, **3** and **4**), the following LC method was used: eluent 20 mM $NH_4OAc$ buffer (pH = 6)/$CH_3CN$, gradient 50-100%, column temperature = 40 °C. For all other compounds (including compound **5**) the following universal LC method was used: eluent 20 mM $NH_4OAc$ buffer (pH = 6)/$CH_3CN$, gradient 0-100% or 0-60%, column temperature = 40 °C.. All eluents were used all in LC-MS grade..

For flash chromatography, an Interchim puriFlash® XS520Plus with multi λ absorbance detector and various types of columns were used: a) 330 g column, 50 µm spherical silica gel, Interchim [PF-50SIHP-F0330]; b) 120 g column, 50 µm spherical silica gel, Interchim [PF-50SIHP-F0120]; c) 80 g column, 25 µm spherical silica gel, Interchim [PF-25SIHC-F0080]. Solid injection (dry load) mode was used for all separations with this technique. We utilized adjustable solid load cartridge system (Teledyne ISCO, 605237048) with various amounts of silica gel (5 – 25g).

For reversed-phase (RP) flash chromatography, a Büchi PrepChrom C-700 with multi λ absorbance detector and Teledyne ISCO column RediSepRf® HP C18 Aq GOLD 50g were used. As eluents dd$H_2O$ and $CH_3CN$ (HPLC grade) were used.

Synthetic procedures

The synthetisis of **5** is shown in Supplementary Figure 56.

2',3',5'-Tris-*O*-(*tert*-butyldimethylsilyl)-5-methyluridine (**1**)

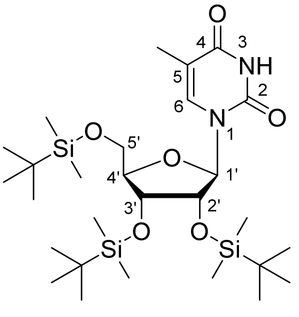

The 5-methyluridine ($m^5U$) (2.15 g, 8.3 mmol, 1 eq.) and 1*H*-imidazole (5.65 g, 83 mmol, 10 eq.) were suspended in dry DMF (28 mL) under argon. The suspension was cooled by an ice-water bath. *Tert*-Butyldimethylsilyl chloride (5 g, 33.2 mmol, 4 eq.) was added and the reaction mixture was stirred overnight to gradually warm to room temperature. The reaction mixture was transferred to a separation funnel with demineralized water ($dH_2O$) (150 mL) and extracted by EtOAc (2× 150 mL). The organic layer was extracted by $dH_2O$ (2× 150 mL) and brine (1× 150 mL) and the resulting EtOAc solution was dried by $MgSO_4$, filtered and evaporated *in vacuo*. Raw product was purified via flash chromatography (330 g $SiO_2$ column, eluent cyclohexane:EtOAc (2 % MeOH), gradient 90:10 to 30:70) to obtain the compound **1** as a white solid (4.55 g, 7.57 mmol, yield: 91 %).

$^1H$ NMR (600 MHz, DMSO-d6, ppm): δ 11.38 (s, 1H, NH), 7.40 (s, 1H, H-6), 5.88 (d, *J* = 6.9 Hz, 1H, H-1′), 4.20 (dd, *J* = 6.9, 4.6 Hz, 1H, H-2′), 4.04 (dd, *J* = 4.6, 1.8 Hz, 1H, H-3′), 3.93 – 3.89 (m, 1H, H-4′), 3.82 (dd, *J* = 11.5, 3.8 Hz, 1H, H-5′$^{a}$), 3.71 (dd, *J* = 11.5, 2.9 Hz, 1H, H-5′$^{b}$), 1.78 (s, 3H, C-5-$\underline{CH_3}$), 0.94 – 0.75 (m, 27H, Si-C-($\underline{CH_3}$)$_3$), 0.08 (dd, *J* = 14.7, 5.7 Hz, 12H, Si-$\underline{CH_3}$), -0.02 (s, 3H, Si-$\underline{CH_3}$), -0.10 (s, 3H, Si-$\underline{CH_3}$).

$^{13}C$ NMR (151 MHz, DMSO-d6, ppm): δ 163.29 (C-4), 150.63 (C-2), 135.04 (C-6), 109.81 (C-5), 86.05 (C-1′), 85.12 (C-4′), 73.93 (C-2′), 72.23 (C-3′), 62.84 (C-5′), 25.73, 25.57 and 25.42 (Si-C-($\underline{C}H_3$)$_3$), 17.96, 17.63 and 17.47 (Si-$\underline{C}$-($CH_3$)$_3$), 12.00 (C-5-$\underline{C}H_3$), -4.82, -4.85, -4.97, -5.28, -5.62 and -5.64 (Si-$\underline{C}H_3$).

HRMS (ESI+) m/z: calculated for $C_{28}H_{56}N_2O_6Si_3$: 623.33384 $[M + Na]^+$, found 623.33366.

2',3',5'-Tris-*O*-(*tert*-butyldimethylsilyl)-5-(bromomethyl)uridine (**2**)

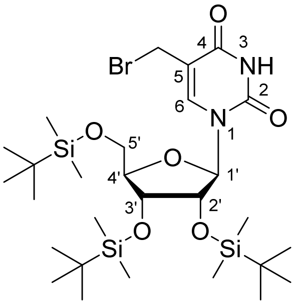

The compound **2** was synthesized according to literature procedure describing the radical bromination of 3′-5′-bis-*O*-(*tert*-butyldimethylsilyl)-thymidine reported by Xiaoxu Li et al. (2018)[1] The uridine derivative **1** (3.12 g, 5.19 mmol, 1 eq.), *N*-bromosuccinimide (1.11 g, 6.24 mmol, 1.2 eq.) and azobisisobutyronitrile (103 mg, 0.63 mmol, 0.12 eq.) were transferred to a round bottom flask. Benzene (55 mL) was added, and the reaction mixture was stirred and heated to 80 °C. The reaction mixture was tested by TLC (6:4 / cyclohexane:EtOAc) and indicated complete conversion of the starting material in 1 hour. The solution was cooled to 40 °C and evaporated *in vacuo*. The crude product was purified via flash chromatography (120 g $SiO_2$ column, eluent DCM:acetone, gradient 0-5 %). The pure fractions were evaporated *in vacuo* and freeze-dried (from 1,4-dioxane) to obtain 1.18 g (yield: 34 %) of compound **2** as a white solid lyophilizate.

$^1H$ NMR (500 MHz, $CDCl_3$): δ 8.51 (s, 1H, NH), 7.86 (s, 1H, H-6), 6.03 (d, *J* = 6.3 Hz, 1H, H-1′), 4.32 (d, *J* = 10.5 Hz, 1H, -$CH_2^{a}$-Br), 4.17 (d, *J* = 10.5 Hz, 1H, -$CH_2^{b}$-Br), 4.11 (dd, *J* = 6.3, 4.1 Hz, 1H, H-2′), 4.07 – 4.03 (m, 2H, H-3′ and H-4′), 3.92 (dd, *J* = 11.6, 2.0 Hz, 1H, H-5′$^{a}$), 3.76 (dd, *J* = 11.6, 1.8 Hz, 1H, H-5′$^{b}$), 0.99, 0.91 and 0.86 (s, 27H, Si-C-($\underline{CH_3}$)$_3$), 0.18 (d, *J* = 1.4 Hz, 6H, Si-$\underline{CH_3}$), 0.10, 0.08, 0.03 and -0.02 (s, 12H, Si-$\underline{CH_3}$).

$^{13}C$ NMR (126 MHz, $CDCl_3$): δ 161.33 (C-4), 149.86 (C-2), 139.55 (C-6), 112.23 (C-5), 87.77 (d, *J* = 6.7 Hz, C-1′), 86.33 (d, J = 4.8 Hz, C-4′), 75.90 (C-2′), 72.47 (C-3′), 63.30 (C-5′), 26.30, 25.95, 25.85 and 25.79 (Si-C-($\underline{CH_3}$)$_3$), 18.78, 18.22 and 18.06 (Si-$\underline{C}$-($CH_3$)$_3$), -4.29, -4.35, -4.54, -4.56, -4.98 and -5.23 (Si-$\underline{CH_3}$).

Compound **2** was very unstable in MS analyses. The LC-MS analysis (low resolution) performed in aqueous buffer (20 mM $NH_4OAc$, pH = 6.0) showed the mass of the hydroxy derivative ($ArCH_2$-OH instead of $ArCH_2$-Br). (ESI+) m/z: calculated for $C_{28}H_{56}N_2O_7Si_3$: 617.3468 $[M + H]^+$, found 617.3. The high resolution MS analysis (Orbitrap XL) performed in MeOH showed only a mass of the methyl ether derivative ($ArCH_2$-$OCH_3$ instead of $ArCH_2$-Br). HRMS (ESI+) m/z: calculated for $C_{29}H_{58}N_2O_7Si_3$: 653.34495 $[M + Na]^+$, found 653.34507. Only the high resolution MS analysis performed in a $CHCl_3$/$CH_3CN$ mixture provided a low abundant but correct molecular ion.

HRMS (ESI+) m/z: calculated for $C_{28}H_{55}BrN_2O_6Si_3$: 701.24435 $[M + Na]^+$, found 701.24460.

2',3',5'-Tris-*O*-(*tert*-butyldimethylsilyl)-5-propargyluridine (**3**)

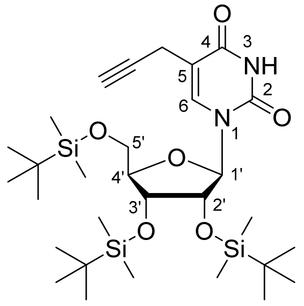

The bromo derivative **2** (900 mg, 1.32 mmol, 1 eq.) and CuCN (39 mg, 0.437 mmol, 0.33 eq.) were dissolved in dry THF (20 mL) in the Schlenk tube under argon atmosphere to form a dark green solution. The solution of ethynylmagnesium bromide in THF (5.8 mL of 0.5 M solution = 2.9 mmol, 2.2 eq.) was added dropwise at room temperature. The color of the solution changed to ochre. The reaction mixture was stirred and heated to 65 °C for 21 hours and then quenched by adding d$H_2O$ (50 mL) and a saturated aqueous solution of $NH_4Cl$ (50 mL) to the mixture. The resulting mixture was extracted by EtOAc (2× 100 mL). The organic layer was extracted by d$H_2O$ (2× 200 mL) and brine (1× 200 mL). The resulting EtOAc solution was dried by $MgSO_4$, filtered and evaporated *in vacuo*. Raw product was purified via flash chromatography (120 g $SiO_2$ column, eluent cyclohexane:EtOAc(2 % MeOH), gradient 95:5 to 60:40) to obtain the propargyl derivative **3** as a white solid (600 mg, 0.96 mmol, yield: 72 %).

$^1$H NMR (500 MHz, $CDCl_3$): δ 8.67 (s, 1H, NH), 7.58 – 7.55 (m, 1H, H-6), 5.97 (d, *J* = 6.0 Hz, 1H, H-1′), 4.16 (dd, *J* = 6.0, 4.5 Hz, 1H, H-2′), 4.07 (dd, *J* = 4.5, 2.9 Hz, 1H, H-3′), 4.04 (q, *J* = 3.7 Hz, 1H, H-4′), 3.83 (dd, *J* = 11.3, 4.0 Hz, 1H, H-5′[a]), 3.76 (dd, *J* = 11.3, 3.5 Hz, 1H, H-5′[b]), 3.35 – 3.20 (m, 2H, -$CH_2$-C≡C), 2.12 (t, *J* = 2.7 Hz, 1H, -C≡C-H), 0.96 – 0.84 (m, 27H, Si-C-($CH_3$)$_3$), 0.12 (d, *J* = 1.2 Hz, 6H, Si-$CH_3$), 0.09, 0.08, 0.07, 0.06, 0.05, 0.03 and -0.02 (s, 12H, Si-$CH_3$).

$^{13}$C NMR (126 MHz, $CDCl_3$): δ 162.36 (C-4), 150.15 (C-2), 137.52 (d, *J* = 5.4 Hz, C-6), 110.33 (C-5), 88.55 (d, *J* = 7.3 Hz, C-1′), 85.70 (d, *J* = 4.9 Hz, C-4′), 79.96 (-C≡C-H), 75.03 (C-2′), 72.37 (C-3′), 71.30 (d, *J* = 3.9 Hz, -C≡C-H), 63.40 (C-5′), 26.30, 26.20, 26.01, 25.96 and 25.85 (Si-C-($CH_3$)$_3$), 18.68, 18.21 and 18.07 (Si-C-($CH_3$)$_3$), 16.59 (-$CH_2$-C≡C), -4.30, -4.37, -4.56, -4.66, -5.15 and -5.19 (Si-$CH_3$).

HRMS (ESI+) m/z: calculated for $C_{30}H_{56}N_2O_6Si_3$: 663.30778 $[M + K]^+$ and 647.33384 $[M + Na]^+$, found 663.30819 and 647.33439.

2',3',5'-Tris-*O*-(*tert*-butyldimethylsilyl)-5-propargylcytidine (**4**)

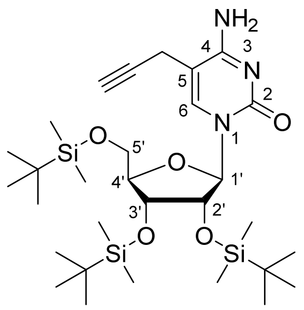

For the preparation of compound **4**, the procedure describing the conversion of a uridine derivative to a cytidine derivative reported by Qi Sun et al. [2] was considerably modified. The propargyl derivative **3** (900 mg, 1.44 mmol, 1 eq.) was dissolved in dry DCM (20 mL) under argon. *N*-Methylpiperidine (460 µL, ~ 2.6 eq.) and triethylamine ($Et_3N$, 460 µL, ~ 2.3 eq.) were added. Subsequently, 4-methylbenzene-1-sulfonyl chloride (800 mg, 4.2 mmol, 2.9 eq.) in dry DCM (6 mL) was added dropwise. The reaction mixture was stirred at room temperature for 3.5 hours and then cooled by an ice-water bath. The solution of $NH_3$ (25 mL of 7N $NH_3$ in MeOH) was added and formation of a temporary precipitate was observed. The solution was stirred for 2 hours and then transferred to a separation funnel with d$H_2O$:$CHCl_3$ (400 mL, 1:1) and extracted. The organic layer was extracted by d$H_2O$ (4×) and brine (1×) and evaporated *in vacuo*. The crude product was purified via flash chromatography (80 g $SiO_2$ column neutralized by DCM (2 % $Et_3N$), eluent cyclohexane:EtOAc (with 2 %MeOH):MeOH, gradient from 90:10:0 to 45:45:10). The pure fractions were evaporated *in vacuo* and freeze-dried (from 1,4-dioxane) to obtain 333 mg (yield: 37 %) of pure compound **4** as a white solid lyophilizate.

$^1$H NMR (600 MHz, $CDCl_3$): δ 7.72 (m, 1H, H-6), 5.91 (d, *J* = 4.0 Hz, 1H, H-1′), 4.15 (t, *J* = 4.1 Hz, 1H, H-2′), 4.08 (dt, *J* = 5.7, 3.0 Hz, 1H, H-4′), 3.99 (dd, *J* = 5.2, 4.3 Hz, 1H, H-3′), 3.96 (dd, *J* = 11.5, 2.9 Hz, 1H, H-5′[a]), 3.78 (dd, *J* = 11.5, 3.1 Hz, 1H, H-5′[b]), 3.27 (d, *J* = 2.7 Hz, 2H, -$CH_2$-C≡C), 2.20 (t, *J* = 2.7 Hz, 1H, -C≡C-H), 0.96, 0.90 and 0.88 (s, 27H, Si-C-($CH_3$)$_3$), 0.14, 0.13, 0.08, 0.07, 0.06, and -0.05 (s, 18H, Si-$CH_3$).

$^{13}$C NMR (151 MHz, $CDCl_3$): δ 139.88 (C-6), 101.23 (C-5), 89.89 (C-1′), 84.48 (C-4′), 78.65 (-C≡C-H), 75.98 (C-2′), 72.31 (-C≡C-H), 71.26 (C-3′), 62.77 (C-5′), 26.31, 26.01 and 25.99 (Si-C-($CH_3$)$_3$), 18.83, 18.32 and 18.16 (Si-C-($CH_3$)$_3$), 18.22 (-$CH_2$-C≡C), -4.08, -4.31, -4.70, -4.73, -4.89 and -5.21 (Si-$CH_3$).

HRMS (ESI+) m/z: calculated for $C_{30}H_{57}N_3O_5Si_3$: 646.34982 $[M + Na]^+$, found 646.35015.

5-Propargylcytidine (**5**) (**prop$^5$C**)

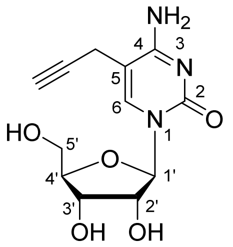

For removing the *tert*-butyldimethylsilyl protection groups, the procedure described by Venkatesham et al. [3] was adapted. The propargyl derivative **4** (22 mg, 35 µmol, 1 eq.) was dissolved in MeOH (4 mL) and $NH_4F$ was added (18 mg, 486 µmol, 14 eq.). The reaction vessel was closed by a screw cap and the reaction mixture was stirred and heated to 65 °C (oil bath). The reaction mixture was repeatedly tested by LC-MS (50 µL of reaction mixture diluted in 1 mL MeOH, 6 µL injection) and indicated almost full conversion of the starting material in 22 hours. The reaction was stopped after 27 hours. The solvent was evaporated *in vacuo*, the residue was suspended in DMSO (1 mL) and injected onto a chromatographic column and purified by RP flash chromatography (C18-aq 50 g; eluent dd$H_2O$/$CH_3CN$, gradient 0–20 %). The pure fraction was freeze-dried to obtain 3 mg (yield: 30 %) of compound **5** as a white lyophilizate. The purity was tested by LC-MS.

$^1$H NMR (500 MHz, DMSO-$d_6$): δ 7.81 (s, 1H, H-6); 5.79 (d, *J* = 4.6 Hz, 1H, H-1′); 5.28 (d, *J* = 5.3 Hz, 1H, -OH-2′); 4.98 (s, 2H, -OH-3′ and -OH-4′); 3.97 – 3.93 (m, 1H, H-2′); 3.93 – 3.88 (m, 1H, H-3′); 3.83 (dt, *J* = 4.8, 3.6 Hz, 1H, H-4′); 3.64 (d, *J* = 11.9 Hz, 1H, H-5′$^a$); 3.54 (d, J = 11.9 Hz, 1H, H-5′$^b$); 3.22 (dd, *J* = 2.7, 1.0 Hz, 2H, -$\underline{CH_2}$-C≡C); 3.03 (t, *J* = 2.6 Hz, 1H, -C≡C-H).

$^{13}$C NMR (126 MHz, DMSO-$d_6$): δ 163.84 (C-4); 155.08 (C-2); 139.28 (C-6); 100.85 (C-5); 89.12 (C-1′); 84.21 (C-4′); 80.24 (-$\underline{C}$≡C-H); 73.92 (-C≡$\underline{C}$-H); 73.83 (C-2′); 69.79 (C-3′); 61.03 (C-5′); 16.99 (-$\underline{C}H_2$-C≡C).

HRMS (ESI+) m/z: calculated for $C_{12}H_{15}N_3O_5$: 304.09039 $[M + Na]^+$, found 304.09049.

Supplementary Note 2: Details about the LC-QqQ-MS instrument and software setup

The LC-QqQ-MS system we used herein is an Agilent 1260 Infinity II LC-system coupled with and an Agilent Ultivo Triple Quadrupole mass spectrometer equipped with an advanced electrospray ionization (ESI) source (JetStream). Chromatographic separation was performed on an Agilent Poroshell 120 EC-C18 3 × 150 mm 2.7 µm column with complementary pre-column Agilent EC-C18 3 x 5 mm 1.9 µm. Dynamic MRM (dMRM) mode was used with defined time windows for each analyte to prevent potential problems with partially overlapping analytes [4].

The system was controlled by the MassHunter Data Acquisition Software (edition for Ultivo, version C.01.00). For basic analysis the MassHunter Qualitative Analysis Navigator (version B.08.00) was used and for quantification the MassHunter Quantitative Analysis Software (edition for QqQ, version B.09.00) was used. For optimization of MRM parameters, the manual methods developing and the MassHunter Optimizer (version C.01.00) were used complementary.

MS data were conducted via ESI in positive ion mode and multiple reaction monitoring (MRM) was employed for the detection and quantification. The ion source parameters used herein include: Main gas temperature = 250 °C, Sheath gas temperature = 375 °C and Capillary Voltage = 2400 V. The mass spectrometr Ultivo has the Cell Acceleration Voltage (CAV) hard set at 9 V.

Supplementary Note 3: Details to prop$^5$C quantification via LC-QqQ-MS.
The calibration curve of prop$^5$C was determined using a synthetic standard (Supplementary Fig. 51) . From a stock solution of prop$^5$C (1.39 pg/µL),0.1, 0.2, 0.5, 1, 2, 5, 10 µL were injected.

For all points of the calibration curve of prop$^5$C, the quantifier peaks showed the characteristic shape and were of very good quality (Supplementary Fig. 52). However, both qualifiers showed a much lower signal cin comparison to the quantifier (Supplementary Fig. 53). At very low concentrations of prop$^5$C, the qualifier peaks collapse (between points 3 (cc3) and 2 (cc2) of the calibration curve inSupplementary Fig. 54). , indicating that the detection limit of the system was reached. On the other hand, the quantifier signal is very good. Thus, for the quantification process itself (which is based on quantifier) the setup was considered sufficient. Taking into account the above, we decided to use the analytical data of prop$^5$C standard and perform the measuring and quantification of the real samples.

Supplementary Note 4: NGS library preparation for MePMe-seq

Two times poly(A)$^+$ enriched RNA (5 µg) was fragmented at 88 °C (25 ng/µL mRNA) in 1× first strand buffer (Invitrogen$^{TM}$) for 4 min, immediately cooled on ice (5 min) and precipitated in EtOH. Successful fragmentation was confirmed via Bioanalyzer RNA analysis. Samples were biotinylated in a CuAAC reaction by combining the preincubated copper catalyst (1:5 $CuSO_4$:THPTA, 5 min at rt) with RNA and biotin azide in sodium-phosphate buffer. The reaction was started by addition of sodium ascorbate (final concentrations: 5 mM $CuSO_4$, 25 mM THPTA, 50 ng/µL RNA, 30 µM biotin azide, 60 mM sodium phosphate buffer, 100 mM sodium ascorbate), incubated for 30 min at 37 °C, stopped by addition of EDTA (10 mM) and directly purified using Microspin G-25, following the manufacturer's instructions. Biotinylated RNA was bound to 500 µg streptavidin-coated magnetic beads (Dynabeads$^{TM}$ M-280 Streptavidin, Invitrogen$^{TM}$), following the manufacturer's instructions for coupling nucleic acids except for two changes: 1) After initial washing, beads were preincubated with 0.2 µM poly(dT)-Oligo (10 min, rt, constant rotation) before adding the RNA (to reduce unspecific background during binding). 2) Incubation time of RNA with the streptavidine beads was prolonged (1 h, rt, constant rotation).

Next, immobilized RNA was dephosphorylated at the 3′ end with T4 PNK (1× T4 PNK buffer A, 0.8 U/µL Ribolock, 0.4 U/µL T4 PNK in 50 µL reaction volume, 30 min, 37 °C, shaking at 1100 rpm) followed by a washing step for which the bead-bound RNA was seperated with a magnet to remove the supernatant, and then washed with 500 µL 1× B&W buffer, followed by 500 µL dd$H_2O$. Subsequently, the L3-Adapter was ligated at the 3′end (1× Ligation buffer, 0.4 U/µL Ribolock, 1.5 µM L3-Adapter, 20 % PEG and 0.5 U/µL T4 RNA ligase 1 in 20 µL final reaction volume, overnight, 16 °C, shaking at 1100 rpm). Beads were magnetically seperated after adding 500 µL 1× B&W buffer to reduce viscosity of the mixture followed by a washing step as described before. Bead-bound RNA was hybridized with the RT primer (0.05 µM) in 10 µL reaction volume for 5 min at 70 °C followed by cooling on ice for 5 min. RT was performed with SSIV (hybridized RNA+primer mix, 1× RT buffer, 5 mM DTT, 0.4 U/µL Ribolock, 1 mM of each dNTP, 1 U/µL SSIV, in 20 µL reaction volume, 30 min, 50 °C). RNA was hydrolyzed (80 mM NaOH, 10 min, 98 °C), the reaction was neutralized (1 eq. HCl) and beads were removed by magnetic precipitation. Resulting cDNA was purified using silane-coated magnetic beads following the protocol described in [5]. Briefly, 10 µL of (MyOne$^{TM}$ Silane Dynabeads$^{TM}$, Invitrogen$^{TM}$) were washed once with RLT buffer (Quiagen$^{TM}$). cDNA was mixed with the washed beads in 125 µL of RLT buffer and precipitated with 1 vol-euivalent of EtOH. After incubation at room temperature (2× 5 min, resuspending in between) beads were washed with ethanol (80 %) trice, precipitated and dried (5 min, rt).

For PCR amplification, a second linker is ligated. The resuspended, immobilized cDNA (in dd$H_2O$) was heated with the respective barcode-linker (L##clip2.0) in DMSO to denature any secondary structures (2 min, 70 °C followed by 1 min, 0 °C) and 1 mM ATP, 0.75 U/µL RNA Ligase (high conc.) and 22.5 % PEG8000 in 1× RNA ligase buffer with DTT (20 µL reaction volume total) were added. After homogenizing the viscous mixture, another 0.3 U RNA Ligase (high conc.) were added and reacted overnight at 16 °C (shaking at 1,100 rpm). Following the ligation, a second clean-up using silane-coated magnetic beads following the protocol from the iCLIP2 protocol[5] was performed adding fresh beads (5 µL). After the washing steps and precipitation, RNA was eluted from the beads in 23 µL dd$H_2O$ (incubation for 5 min at rt) and beads were removed by magnetic precipitation.

Purified cDNA was used in a first PCR amplification (0.2 mM of each dNTP, 0.5 µM of each Solexa_s primer (rev and fwd) and 0.02 U/µL Phusion DNA polymerase in 1× HF buffer, 98 °C for 30 s, 6x[98 °C for 10 s, 5 °C for 30 s, 72 °C for 30 s], 72 °C for 3 min) to reduce loss in the first size amplification step. To remove adapter and primer contaminations from previous steps, reaction was size-selected using the ProNex size-selective purification system (Promega) following the manufacturer's instructions using a 3:1 v/v ratio of ProNex Chemistry to sample. Deviating from the protocol, the pre-library was eluted from the beads in 23 µL ddH2O instead of the provided elution buffer.

Cycle number for the final amplification of the libraries was determined by performing test-PCR reactions using 1 µL of the pre-library for 10 µL of reaction split to 3 aliquots (0.2 mM of each dNTP, 0.5 µM of each Solexa primer (rev and fwd) and 0.02 U/µL Phusion DNA polymerase in 1× HF buffer amplified with different cycle numbers (98 °C for 30 s, 15-19 x [98 °C for 10 s, 5 °C for 30 s, 72 °C for 30 s], 72 °C for 3 min). Checking the size profile of these libraries either on a nat. PAGE or via bioanalyzer, cycle numbers were chosen to ensure sufficient amplification without overamplification (increasing background amplification) or adapter-dimers (defined bands at ~ 170 bp typically separate from the broader library peak at 250-600 bp). Final PCR was performed with the determined cycle number (reduced by one cycle because the input is increased) in two separate reactions to ensure backup in case of unforeseen problems (10 µL of pre-library, 0.2 mM of each dNTP, 0.5 µM of each Solexa primer (rev and fwd) and 0.02 U/µL Phusion DNA polymerase in 1× HF buffer). The final library was purified by size-selection using the ProNex size-selective purification system (Promega) following the manufacturer's instructions using a 2.4:1 v/v ratio of ProNex Chemistry to sample and eluted from the beads in 20 µL dd$H_2O$ instead of the provided elution buffer. Quality of the finished libraries was checked via bioanalyzer-High sensitivity DNA assay (Agilent) and concentration was determined by Quant it PicoGreen Assay (Thermo Fisher).
